# The making of a Lewy body: the role of α-synuclein post-fibrillization modifications in regulating the formation and the maturation of pathological inclusions

**DOI:** 10.1101/500058

**Authors:** Anne-Laure Mahul-Mellier, Melek Firat Altay, Johannes Burtscher, Niran Maharjan, Nadine Ait-Bouziad, Anass Chiki, Siv Vingill, Richard Wade-Martins, Janice Holton, Catherine Strand, Caroline Haikal, Jia-Yi Li, Romain Hamelin, Marie Croisier, Graham Knott, Georges Mairet-Coello, Laura Weerens, Anne Michel, Patrick Downey, Martin Citron, Hilal A. Lashuel

**Author notes:** To whom correspondence should be addressed at: Laboratory of Molecular and Chemical Biology of Neurodegeneration, Brain Mind Institute, Ecole Polytechnique Fédérale de Lausanne, 1015 Lausanne.

## Abstract

Although converging evidence point to α-synuclein (α-syn) aggregation and Lewy body (LB) formation as central events in Parkinson’s disease (PD), the molecular mechanisms that regulate these processes and their role in disease pathogenesis remain poorly understood. Herein, we applied an integrative biochemical, structural and imaging approach to elucidate the sequence, molecular and cellular mechanisms that regulate LB formation in primary neurons. Our results establish that post-fibrillization C-terminal truncation mediated by calpains 1 and 2 and potentially other enzymes, plays critical roles in regulating α-syn seeding, fibrillization and orchestrates many of the events associated with LB formation and maturation. These findings combined with the abundance of α-syn truncated species in LBs and pathological α-syn aggregates have significant implications for ongoing efforts to develop therapeutic strategies based on targeting the C-terminus of α-syn or proteolytic processing of this region.

## Introduction

The intracellular accumulation of aggregated forms of alpha-synuclein (α-syn) occurs in neurons and glial cells, and represents one of the main pathological hallmarks of Parkinson’s disease (PD) and related synucleinopathies, including dementia with Lewy bodies (DLB) and multiple system atrophy (MSA)^1^. PD and DLB are characterized by the presence of Lewy bodies (LB) in subcortical and cortical neurons, while in MSA, α-syn inclusions are mainly detected in glial cells and are referred to as glial cytoplasmic inclusions (GCIs). Although it is widely believed that α-syn plays a central role in the pathogenesis of synucleinopathies, there is a poor understanding of the molecular and cellular processes that trigger and govern the misfolding, fibrillization, inclusion formation, and spread of α-syn in the brain. Furthermore, there is a debate regarding whether the processes of α-syn fibrillization and inclusion formation are protective or neurotoxic, which remains a subject of active investigation. Addressing this knowledge gap should allow us to advance our understanding of the disease, improve our preclinical models, and develop more effective diagnostic and therapeutic strategies for PD and other synucleinopathies.

Post-mortem observations of brain samples from PD patients who had undergone embryonic graft transplantation have shown that LB pathology can spread from the host brain to the transplant^2^. In experimental systems, exogenously added misfolded α-syn can act as a seed to initiate the misfolding and aggregation of endogenous α-synuclein in both cellular and animal models^3^. This has led to the intriguing hypothesis that α-syn pathology may spread throughout the brain in a prion-like manner^4^. These discoveries provide an attractive mechanism to explain the progression of α-syn pathology in PD, and have fuelled the pharmaceutical industry to develop intervention strategies based on targeting the secretion of α-syn and/or blocking its uptake and cell-to-cell transmission.

The primary event responsible for the initiation and propagation of this pathogenic process involves the interaction between misfolded seeds of α-syn and endogenous proteins in recipient cells. Therefore, we hypothesized that a better understanding of the sequence, molecular and cellular determinants of α-syn seeding and inclusion formation could offer insight into more effective strategies to inhibit both α-syn aggregation and the spread of pathology at different stages of disease progression. Given that α-syn in pathological aggregates is known to be modified at multiple residues, we sought to investigate the role of post-translational modifications (PTMs) in 1) the processing of α-syn pre-formed fibrils (PFFs); 2) α-syn seeding capacity and fibril formation; and 3) the formation and maturation of LB-like inclusions in a primary neuronal model of synucleinopathies. In this model, once internalized into the cells, small amounts of extracellular α-syn PFF seeds (PFFs) are sufficient to induce the aggregation of endogenous soluble α-syn^3^ into fibrils and to initiate the formation of cytoplasmic inclusions. These structures share many of the features which characterize pathological α-syn inclusions in PD and synucleinopathies. These similarities include the formation of fibrillar aggregates that are immunoreactive to α-syn phosphorylated at residue serine 129 (pS129), ubiquitin and p62^5^. Further advantages of this model are: 1) α-syn seeding and fibril formation can be induced at nanomolar concentrations and without the need to overexpress the protein; 2) PFFs, the primary initiators of the process, are added exogenously to the neurons and can be prepared using α-syn monomers bearing site-specific modifications, thus enabling systematic investigation of the role of PTMs in α-syn seeding; and 3) it enables detailed studies to monitor changes in α-syn biochemistry, localization, modifications, interactome, and aggregation over time using advanced methods such as mass-spectrometry and correlative light electron microscopy (CLEM). To achieve in-depth characterization of the mechanisms of seeding and pathology formation in this neuronal model system, we investigated the sequence, as well as molecular and cellular determinants of these processes using integrative cutting-edge approaches, including biochemical characterization, confocal and CLEM imaging, and quantitative proteomic analyses. We uncovered novel mechanisms governing α-syn seeding activity and formation of LB-like inclusions and demonstrate that post-fibrillization PTMs (e.g. C-terminal truncations) are not only markers of α-syn pathology, but also serve as master regulators of α-syn seeding, fibril growth, and the formation and maturation of LB-like inclusions. The results have significant implications for the development of α-syn-targeted diagnostics and therapies and underscore the importance of expanding the existing tool box that is commonly used to investigate α-syn pathology formation and pathology spreading in humans and animal models of synucleinopathies. Accurate assessments and profiling of the pathological diversity of α-syn can only be achieved by using a combination of well-characterized tools that enable detection and characterization of the different aggregated and modified forms of the protein.

## Results

### C-terminal truncation of PFFs is an early event during the α-syn seeding process after internalization

To investigate the role of PTMs in regulating the seeding and formation of α-syn inclusions, we took advantage of the neuronal seeding model developed by Virginia Lee and coworkers^6,7^. In this model, α-syn intracellular aggregation is triggered by the addition of a nanomolar concentration (70 nM) of extracellular PFFs (Figure S1), which induce the formation of intracellular LB-like inclusions in a time-dependent manner (over 7-21 days) upon internalization (Figure 1A). Immunocytochemistry (ICC) revealed that these inclusions contained α-syn pS129 (Figure 1B). The aggregates were also positive for two other well-established LB markers, namely ubiquitin (Ub) and p62^5^, which are two central players in the proteasomal and autophagic degradation pathways (Figures 1D-E, S2D-E).

**Figure 1.**
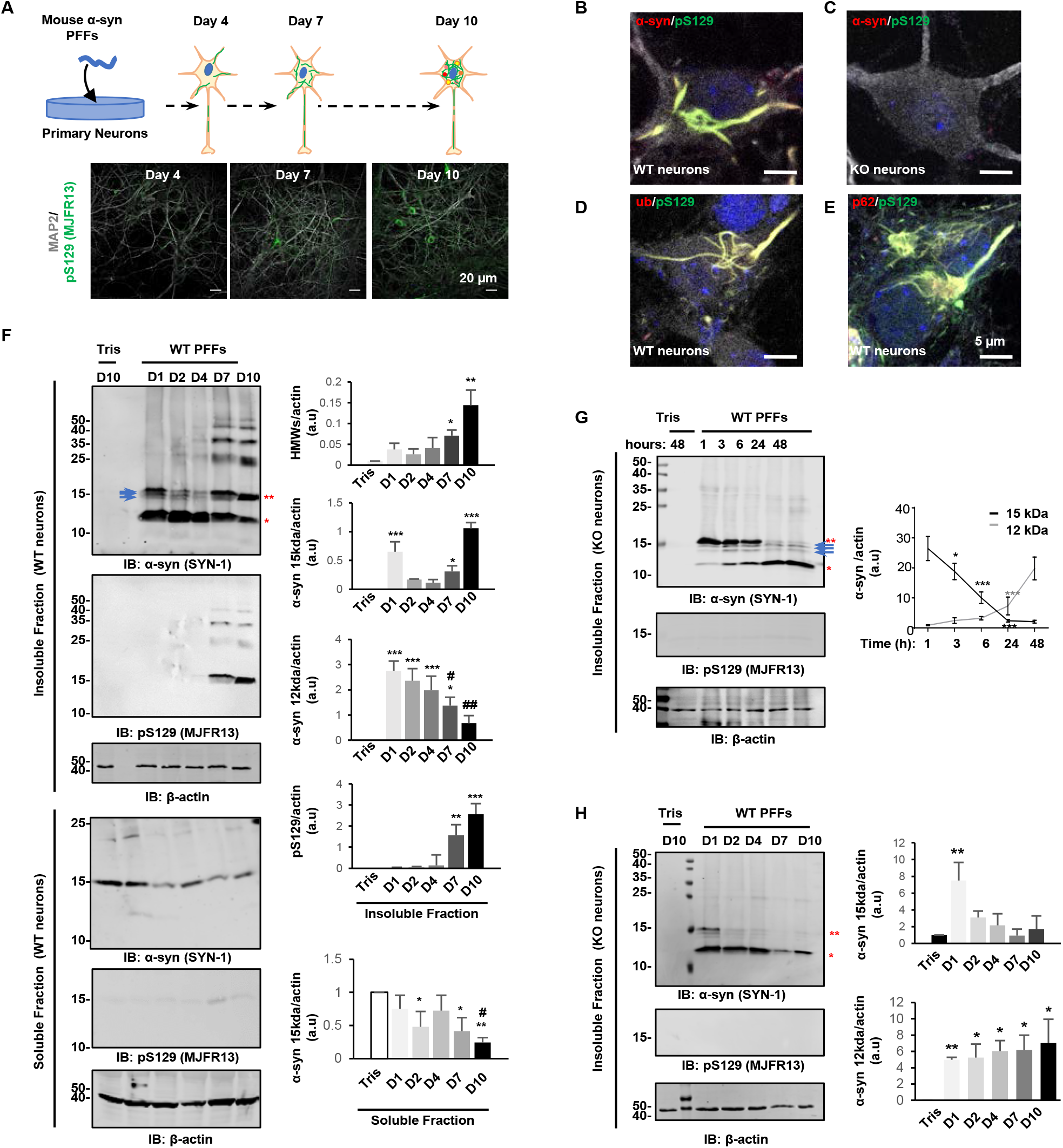
C-terminal truncation is an early event during the seeding process. **A.** Seeding model in hippocampal primary neurons. 70 nM of mouse PFFs were added to neurons at DIV 5 (day *in vitro).* Control neurons were treated with Tris buffer used to prepare PFFs. After 4 days of treatment, positive pS129-α-syn aggregates were detected in the extension of the neurons. After 7 days of treatment, the aggregates appeared in the cytosol of the neurons. The number of LB-like inclusions increased over time, as shown at 10 days of treatment. Scale bars = 20 μm. **B-E.** ICC analysis of the LB-like inclusions that formed at 10 days after adding mouse PFFs to WT neurons (B, D-E) or in KO neurons (C). Aggregates were detected using pS129 (MJFR13) in combination with total α-syn (SYN-1), p62, or ubiquitin antibodies. Neurons were counterstained with microtubule-associated protein (MAP2) antibody, and the nucleus was counterstained with DAPI staining. Scale bars = 5 μm. **F-H**. WB analyses of the LB-like inclusions that formed over time after adding mouse PFFs to WT neurons **(F)** or KO neurons **(G-H)**. Control neurons were treated with Tris buffer (Tris). After sequential extractions of the soluble and insoluble fractions, cells lysates were analysed by immunoblotting. Total α-syn, pS129 and actin were respectively detected by SYN-1, pS129 (MJFR13), and actin antibodies. Levels of total α-syn (15 kDa, indicated by a double red asterisks; 12 kDa indicated by a single red asterisk or HMW) or pS129-α-syn were estimated by measuring the WB band intensity and normalized to the relative protein levels of actin. Blue arrows indicate the intermediate α-syn-truncated fragments. The graphs represent the mean +/−SD of 3 independent experiments. **(F)** p<0.01=*, p<0.001=**, p<0.0001=*** (ANOVA followed by Tukey HSD post-hoc test, Tris vs. PFF-treated neurons) and p<0.01=#, p<0.001=## (ANOVA followed by Tukey HSD post-hoc test, PFF-treated neurons D10 vs. D7 or D4 or D1). **(G-H)** p<0.01=*, p<0.0001=*** (ANOVA followed by Tukey HSD post-hoc test, level of α-syn 15 kDa at 1 hour vs. other time-points or levels of α-syn 12 kDa at 1 hour vs. other time-points or Tris vs. PFF-treated neurons).

Within the same population of treated neurons, we observed different types of α-syn inclusions that appear to differ in shape and subcellular localization over time (Figures S2A-E), despite similar immunoreactivity of the LB markers (α-syn-pS129, Ub, and p62) (Figures 1D-E). In our hands, addition of 70 nM of PFFs barely induced toxicity in neurons, even after 21 days of treatment (Figures S2F-H).

To determine the role of PTMs in regulating α-syn seeding and aggregation, we first sought to achieve a better understanding of the major types of PTMs that occur during the early seeding events and explored their role in regulating α-syn inclusion formation and maturation. We first monitored changes in α-syn modifications and aggregation species over time by western blotting (WB) and showed that the endogenous α-syn shifted from the soluble to the insoluble fraction (Figure 1F) over time in the WT primary culture treated with WT PFFs.

Consistent with the ICC data in Figure 1B, the monomeric (15 kDa) and high-molecular-weight (HMW) bands (~23, 37, 40, and 50 kDa) derived from α-syn inclusions and stained with a pan-synuclein antibody (SYN-1) (Figure 1F, top panel) were also detected with a specific antibody against pS129 (Figure 1F, middle panel). This suggests that α-syn in the inclusion fraction is phosphorylated at S129. The epitopes of the antibodies used are summarized in supplemental Figure 3.

Interestingly, we did not observe the accumulation of pS129 α-syn immunoreactive species in the insoluble fractions during the first 4 days post-seeding, suggesting that phosphorylation at S129 is not required for the initial aggregation events. This could also be explained by the great majority of α-syn species that accumulate in the insoluble fractions between 1 and 4 days being truncated and running with an apparent molecular weight (MW) of 12 kDa. The 12 kDa band was detected by the pan α-syn antibody (SYN-1, epitope 91-99) (Figure 1F, top panel) but not using antibodies against pS129 (Figure 1F middle panel) or against the C-terminal residues 116 to 138 (Figure S6G). This suggests that these species correspond to C-terminally truncated forms of α-syn. Notably, cleaved α-syn was not detected in the insoluble fraction extracted from pure primary cultures of mouse cortical astrocytes treated for 10 days with 70 nM of PFFs (Figure S2I), which confirms that the truncation of α-syn occurs mainly in the neuronal population of the hippocampal primary culture.

We consistently observed that C-terminal truncation fragments appeared rapidly after the internalization of PFFs into the neurons, so we determined whether truncation occurred first on the α-syn PFF seeds and whether it could represent an early key step in the seeding process. We therefore monitored the extent of PFF cleavage after internalization into α-syn knockout (KO) neurons by confocal imaging and WB approaches. The use of α-syn KO neurons allowed us to monitor the fate of PFFs without interference from the formation of new inclusions since these processes do not occur in the absence of endogenous α-syn^6^ (Figures 1G-H).

Using fluorescently labelled PFFs (Figure S1, mouse PFFs^488^), we confirmed by confocal imaging that the seeds were internalized through the endolysosomal pathway and accumulated in late endosomes that were positively stained for LAMP1 marker, as shown previously ^8^ (Figures S2J-K).

During the first hours after internalization in the KO neurons, WB analyses revealed that PFFs were truncated into four fragments with apparent MWs of 15 to 12 kDa. This resulted in the complete loss of full-length α-syn (15 kDa) within 24 hours (Figure 1G). This disappearance correlated well with the appearance of the 12 kDa band, which was the main species 24 hours after addition to the cells. Three additional bands were observed just below the α-syn full-length band within the first hour but did not seem to increase in intensity with time, suggesting that these fragments are rapidly cleaved to generate the 12 kDa fragment (Figures 1G-H, top panel, arrows).

Next, we compared the level of full-length and truncated α-syn over time in both WT (Figure 1F) and KO (Figure 1G-H) neurons. In KO neurons, α-syn PFF seeds were completely cleaved after 24 hours. In WT neurons, the level of intact α-syn at 15 kDa was also massively reduced during the first four days after adding α-syn PFF seeds. However, it markedly increased again after day 7, which was concomitant with the appearance of the HMW bands and the positive signal for pS129 (Figure 1F) associated with the aggregation of the endogenous protein.

### PFF cleavage and generation of α-syn truncation after internalization is a general phenomenon that occurs in different models of α-syn pathology formation and spreading and in human MSA brains

To our knowledge, no report has investigated the role of post-fibrillization α-syn cleavage during the process of LB-like aggregate formation in neuronal seeding models^6,7,9,10^. However, it is important to note that in the great majority of the relevant studies (Figure S4), the detection of total α-syn by the WB approach was solely based on the use of C-terminal antibodies, which do not allow the identification of C-terminally truncated fragments lacking the S129 or other key residues within the epitopes of these antibodies. In studies where N-terminal or NAC antibodies were used, α-syn truncated species could be detected, but their sequences were not defined. Thus, to determine the pathological relevance of C-terminal cleavage, we compared the extent of α-syn truncation and mapped the cleavage products in different models of α-syn pathology.

Since α-syn seeding has been previously shown to occur in mammalian cell lines ^9–11^, we verified the presence of α-syn truncation in HeLa cells overexpressing human α-syn and treated with human PFFs (Figures S1, human WT PFFs and S5A-C). Similar to our observations in neurons, different types of pS129 positive inclusions were observed in the same population of treated HeLa cells (Figure S5B). As early as 4 hours after the delivery of the human PFFs, α-syn-truncated fragments similar to those found in the neuronal seeding models were detected in the insoluble fraction of HeLa cells with a gradual increase in the amount of the main truncated form of α-syn (~12 kDa) over time (Figure S5C).

We next assessed the extent of truncation in several neuronal seeding models. WB analyses clearly showed a similar pattern of truncation in all types of neurons with α-syn cleaved into ~12 kDa fragments and only a low amount of full-length α-syn remaining at 15 kDa. This occurred 10 days after the addition of PFFs into hippocampal or cortical primary neurons from mice (Figure 2A) or hippocampal, cortical, or striatal primary neurons from rats (Figure 2B). Interestingly, the HMW bands positively stained with SYN-1 antibody in the insoluble fraction were detected at the same size (~23, 37, 40, and 50 kDa) in all types of neurons.

**Figure 2.**
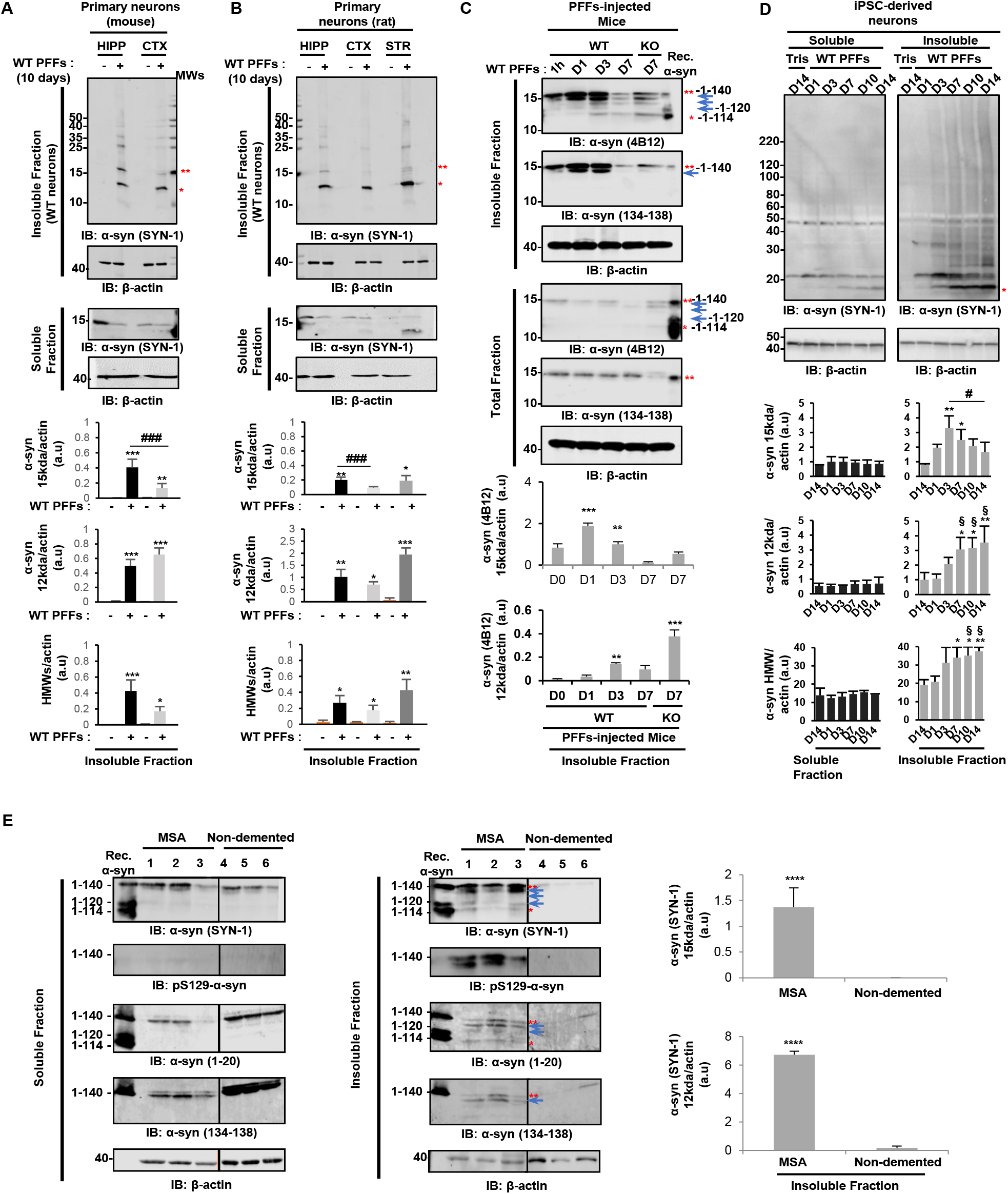
C-terminal truncation is a general phenomenon in α-syn seeding and inclusion formation in cells. WB analyses of the truncation pattern of α-syn in primary neurons **(A-B)**, *in vivo* after injection of human PFFs in the striatum of WT and KO mice **(C)**, in iPSC-derived neurons from a healthy control individual transduced with human PFFs, (D) or in human brain tissue from MSA and non-demented patients **(E)**. Hippocampal (HIPP) and cortical (CTX) primary neurons from mice or **(A)** hippocampal (HIPP), cortical (CTX), or striatal (STR) primary neurons from rats **(B)** were treated for 10 days with 70 nM of mouse PFFs. iPSC-derived neurons were treated with 70 nM of human PFFs for 1, 3, 7, 10, and 14 days **(D)**. Control neurons were treated with Tris buffer (-). Striatum of WT or α-syn KO mice were dissected after 1 hour or 1, 3, or 7 days after injection with human PFFs^WT^ **(C)**. After sequential extractions of the soluble and insoluble fractions, cells lysates were analysed by immunoblotting. The levels of total α-syn (SYN-1, 4B12, or 134-138 antibodies) (15 kDa, indicated by a double red asterisks; 12 kDa indicated by a single red asterisk or HMW) were estimated by measuring the WB band intensity and normalized to the relative protein levels of actin. Blue arrows indicate the intermediate α-syn-truncated fragments. The graphs represent the mean +/− SD of 3 independent experiments. p<0.01=*, p<0.001=**, p<0.0001=*** (ANOVA followed by Tukey HSD post-hoc test, Tris vs. PFF-treated neurons). p<0.0001= ### (ANOVA followed by Tukey HSD post-hoc test, hippocampal vs. cortical neurons). p<0.01=§ (ANOVA followed by Tukey HSD post-hoc test, D1 vs. D7, D10 or D14, iPSC-derived neurons). p<0.01=# (ANOVA followed by Tukey HSD post-hoc test, D3 vs. D14, iPSC-derived neurons). p<0.00001=**** (ANOVA followed by Tukey HSD post-hoc test, MSA vs. non-demented patients).

Strikingly, a similar truncation pattern was also observed *in vivo* after injecting human PFFs in the striatum of C57BL/6J WT mice or α-syn KO mice (Figure 2C). To follow the fate of the seeds specifically, we used an antibody raised against human α-syn (clone 4B12). As early as 1 day after injection, four cleaved fragments running at sizes similar to those observed in primary neurons were detected in the insoluble striatal fractions of the rodent brains using α-syn antibody raised against the epitope 103-108 (clone 4B12), but not with an antibody specific for the C-terminal domain (epitope: 134-138). This suggests that these fragments could result from C-terminal truncations.

To explore the pathophysiological relevance of our findings, we next investigated the processing of fibrillar α-syn in human induced pluripotent stem cells (iPSCs) that were differentiated into dopaminergic neurons. We used a line derived from a healthy control individual, which showed dense processes and expressed markers consistent with human midbrain neurons (Figure S5D). 24 hours after adding human PFFs (Figure S1, human WT PFFs) to the iPSC-derived human neuronal culture (50 days *in vitro*), α-syn was cleaved into a ~12 kDa fragment (Figure 2D), which was similar to our observations of mouse neurons. Finally, WB analyses of human tissues extracted from MSA brain patients (Figure S5F) clearly revealed C-terminally truncated α-syn, which was detected with an N-terminal antibody (epitope: 1-20) or the human specific antibody (clone 4B12, epitope: 103-108), but not with an antibody specific for the C-terminal domain (epitope: 134-138) (Figure 2E). Altogether, our data demonstrate that the truncation of α-syn fibrils is a general and early event that occurs during the formation of intracellular LB-like α-syn inclusions in all neuronal seeding models tested and, even more importantly, in brain tissues from MSA patients. Our results suggest that C-terminal truncation may play a central role in regulating the seeding capacity of α-syn and/or the maturation of α-syn aggregates into LB.

### α-syn PFF seeds C-terminal cleavage occurs primarily at residue 114

To investigate the mechanisms underlying α-syn cleavage in the neuronal seeding model, we next used quantitative proteomic analyses to determine the exact truncation sites of mouse α-syn PFF seeds. N-terminal truncation has been reported previously for cell culture seeding models^11,12^, but our proteomic analyses showed that the N-terminal domain of α-syn was intact in both the 15 and 12 kDa bands sliced from the SDS-PAGE gel (Figure S6A). These findings are consistent with our observation that antibodies raised against N-terminal residues 1-5 or 1-20 were still able to detect α-syn-cleaved products (~12 kDa) in the insoluble fraction of KO neurons (Figures S6E-F). In line with the WB data in Figure 1H, our proteomic analyses demonstrated that PFFs were cleaved at several sites within the C-terminal domain: Asp-135, Ser-129, and Asp-119 were detected in the upper band extracted at ~13-15 kDa, whereas a cleavage occurred at Glu-114 in the band around ~12 kDa (Figures 3A-B and S6B-D). Therefore, liquid chromatography-tandem mass spectrometry (LC-MS/MS) analyses clearly established that the smallest and main fragment that accumulates as the predominant species in neurons seeded with PFFs ends at residue 114 (1-114, MW =11567.20 Da) (Figure 3B). Strikingly, the cleavage sites identified in mouse α-syn were close to those detected in the LBs from human brain tissue^13,14^ or in an α-syn neuroblastoma cell line seeding model^11^. This suggests the involvement of specific proteases in cellular responses to PFFs.

**Figure 3.**
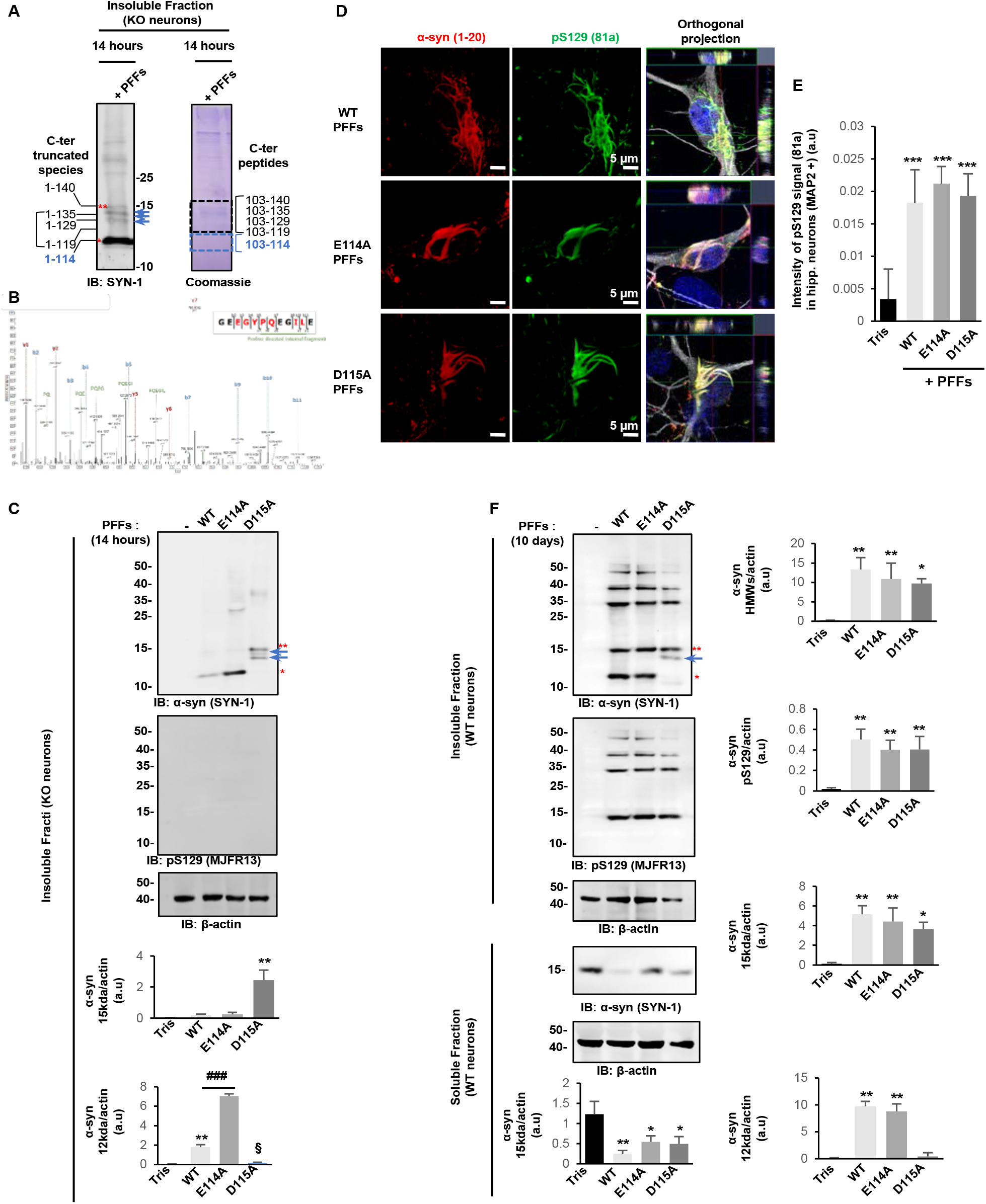
Preventing α-syn cleavage at residue 114 does not impact the seeding capacity of PFFs in primary neurons and mice. **A**. Insoluble fractions of α-syn KO primary neurons treated with 70 nM of mouse PFFs for 14 hours were separated on a 16.5% SDS-PAGE gel. After Coomassie staining, two bands at ~15 (indicated by a double red asterisks) and 12 kDa (indicated by a single red asterisk) were extracted from SDS-PAGE gels (**Figure S6A-B**). Blue arrows indicate the intermediate α-syn-truncated fragments. Isolated bands were selected based on the size of the proteolytic fragments observed by WB and subjected to proteolytic digestion followed by LC-MS/MS analysis. Glu-C was used to analyse N-terminal truncation^68^, which enables detection by LC-MS/MS of intact peptide corresponding to the first 13 amino acids of α-syn. To analyse C-terminal truncations, α-syn was digested using trypsin, which cannot cleave the C-terminal domain, thus allowing detection of an intact peptide corresponding to the 103-140 region by LC-MS/MS^68^. Proteomic analyses revealed that the C-terminal domain was cleaved at residues Asp-135, Ser-129, and Asp-119 with a predominant site of cleavage at residue Glu-114. This resulted in the formation of three fragments (1-135, 1-129, and 1-119) detected in the upper band sliced in and one main fragment (1-114) found in the lower band excised. **B**. Proteomic analysis showing the predominant site of cleavage at residue Glu-114. **C-E**. α-syn KO neurons **(C)** or WT neurons **(D-E)** were treated for 14 hours or 10 days, respectively, with PFFs^WT^, PFFs^E114A^, and PFFs^D115A^. At the indicated time, neurons were fixed for imaging analyses with confocal imaging **(D)** and quantitative HTS **(E)** or lysed and analysed by WB **(C, F). C, F**. Level of total α-syn (SYN-1 antibody) (15 kDa, indicated by a double red asterisks; 12 kDa, indicated by a single red asterisk or HMW) and pS129 were estimated in the soluble and insoluble fractions of KO neurons **(C)** and WT neurons **(F)** by measuring the WB band intensities normalized to the relative protein levels of actin. Blue arrows indicate the intermediate α-syn-truncated fragments. **D-E.** Newly formed inclusions were detected using pS129 antibody (81a) in WT neurons after 10 days of treatment. Neurons were counterstained with MAP2 antibody, and the nucleus was counterstained with DAPI staining. **(D)** Representative confocal images. Scale bar = 5 μm. **(E)** Quantification of images acquired by a high-throughput wide-field cell imaging system. For each independent experiment, duplicated wells were acquired per condition, and nine fields of view were imaged for each well. Each experiment was reproduced at least 3 times independently. Images were then analysed using Cell profile software to identify and quantify the level of LB-like inclusions (stained with pS129 antibody, 81a clone) formed in neurons (MAP2-positive cells). The graphs **(C, E-F)** represent the mean +/− SD of three independent experiments. p<0.01=*, p<0.0001=**, p<0.0001=*** (ANOVA followed by Tukey HSD post-hoc test, Tris vs. PFF-treated neurons). p<0.0005=### (ANOVA followed by Tukey HSD post-hoc test, PFFs^WT^ vs. mutants PFF-treated neurons).

To validate the mass spectrometry data, we further characterized the C-terminal truncated fragments using a set of antibodies raised against the non-amyloid-β component (NAC) or the C-terminal region of α-syn (Figures S6E, G).

### Preventing α-syn cleavage at residue 114 does not impact the seeding capacity of PFFs in primary neurons

Given the rapid proteolytic processing of PFFs upon internalization, we hypothesized that C-terminal cleavage of the PFFs might be a prerequisite for the initiation of α-syn aggregation in neurons. We observed several cleaved fragments (1-114, 1-119, 1-129, and 1-135) and sought to determine whether these cleavages occurred sequentially or independently. Towards this goal, we generated PFFs from α-syn mutants that were designed to block cleavage via the deletion or single substitution of amino acids at the putative cleavage sites (α-syn^1–114^, α-syn^Δ111-115^, α-syn^Δ111-115 Δ133-135^, α-syn^E114A^, α-syn^D115A^) (Figures S1). We first analysed the fate of these different types of PFFs in KO neurons to specifically follow the cleavage of the seeds in the absence of endogenous α-syn and seeding mechanism. As expected, WB analyses of the insoluble fraction of neurons treated with PFFs^1–114^ showed only one band corresponding to the α-syn 1-114 fragment (Figure S7A, single red asterisk). In contrast, PFFs derived from mutants that were designed to prevent cleavage around residue 114, namely PFFs^Δ111-115^ and PFFs^Δ111,115Δ133-135^, did not undergo cleavage to produce the 1-114 fragment (Figure S7A). In neurons treated with PFFs^Δ111-115^, three main fragments were detected: 1-140, 1-135, and 1-129 (Figure S7A-B, blue arrows). PFFS^Δ111-115Δ133-135^ were resistant to C-terminal cleavage at residue 114, which led to an accumulation of the 1-140 full-length protein and the release of a minor product corresponding to the 1-129 fragment (Figure S7A, blue arrows).

Similarly, biochemical analysis of the insoluble fraction of mouse striatum dissected 7 days after injection with PFFs^WT^ revealed an α-syn fragment corresponding to the size of the 1-114 recombinant protein. However, the α-syn 1-114 fragment was not detectable in insoluble fractions from brains injected with PFFs^Δ111-115^ or PFFs^Δ111-115Δ133-135^ (Figure S7C, single red asterisk). These findings demonstrate that the primary C-terminal cleavages occur around residue 114 and between residues 129-135. To pinpoint the exact residue required for the formation of the 1-114 fragment, we next used PFFs carrying a single amino acid substitution at residues 114 (E114A) or 115 (D115A) (Figure S1, mouse E114A and D115A PFFs) and evaluated their capacity to be processed. The data showed that 24 hours after their addition to the KO neurons, α-syn^E114A^ PFFs but not α-syn^D115A^ PFFs underwent cleavage to generate the 1-114 fragment (Figure 3C), thus establishing that the residue D115 is critical for the truncation of α-syn fibrils.

Next, the seeding capacity of PFFs^E114A^ and PFFs^D115A^ was compared to that of PFFs^WT^ after 10 days of treatment in WT neurons. PFFs^WT^, PFFs^E114A^, PFFs^D115A^ all induced a similar level of seeding in WT neurons (Figures 3D-E). These findings were confirmed by the WB analyses of the insoluble fractions extracted from WT neurons treated with PFFs^WT^, PFFs^E114A^, or PFFs^D115A^, in which a similar level of pS129 and HMW signals was detected (Figure 3F). Accordingly, the level of endogenous α-syn remaining in the Triton-soluble fraction of PFF-treated neurons showed that PFFs^E114A^ and PFFs^D115A^ induced similar levels of aggregation compared to neurons treated with PFFs^WT^ (Figure 3F). Altogether, the results suggest that blocking the C-terminal truncation of the seeds does not prevent the seeding and the recruitment of endogenous α-syn or the formation of aggregates in neurons. This is in line with previous studies reporting that the deletion of various α-syn regions other than the NAC domain does not inhibit the formation of the LB-like structures^6,10^.

### α-syn newly formed fibrils are C-terminally cleaved during their growth and maturation in primary neurons

We next assessed whether newly formed fibrils were also altered by these cleavages. Towards this goal, we further investigated the potential interplay in cells between C-terminal truncation and the formation and processing of newly formed α-syn fibrils. We first determined the proportion of the internalized seeds present in the newly formed aggregates. Using fluorescently labelled PFFs^AF488^, ICC and confocal imaging, we showed that the internalized seeds are present in the newly formed aggregates as a minor species (Figure S8A-B).

We next used quantitative proteomic analyses (Figures 4A-B) combined with WB (Figures 4C-E) and ICC (Figures 4F-I) analyses to determine if the newly formed fibrils were also altered by truncation during the aggregation process. The proteomic and WB (epitopes: 1-5 and 1-20) analyses showed that the N-terminal domain of α-syn was intact in both the 12 and 15 kDa bands and in the HMW species. This suggests that that the newly formed aggregates are not subjected to N-terminal proteolytic processing.

**Figure 4.**
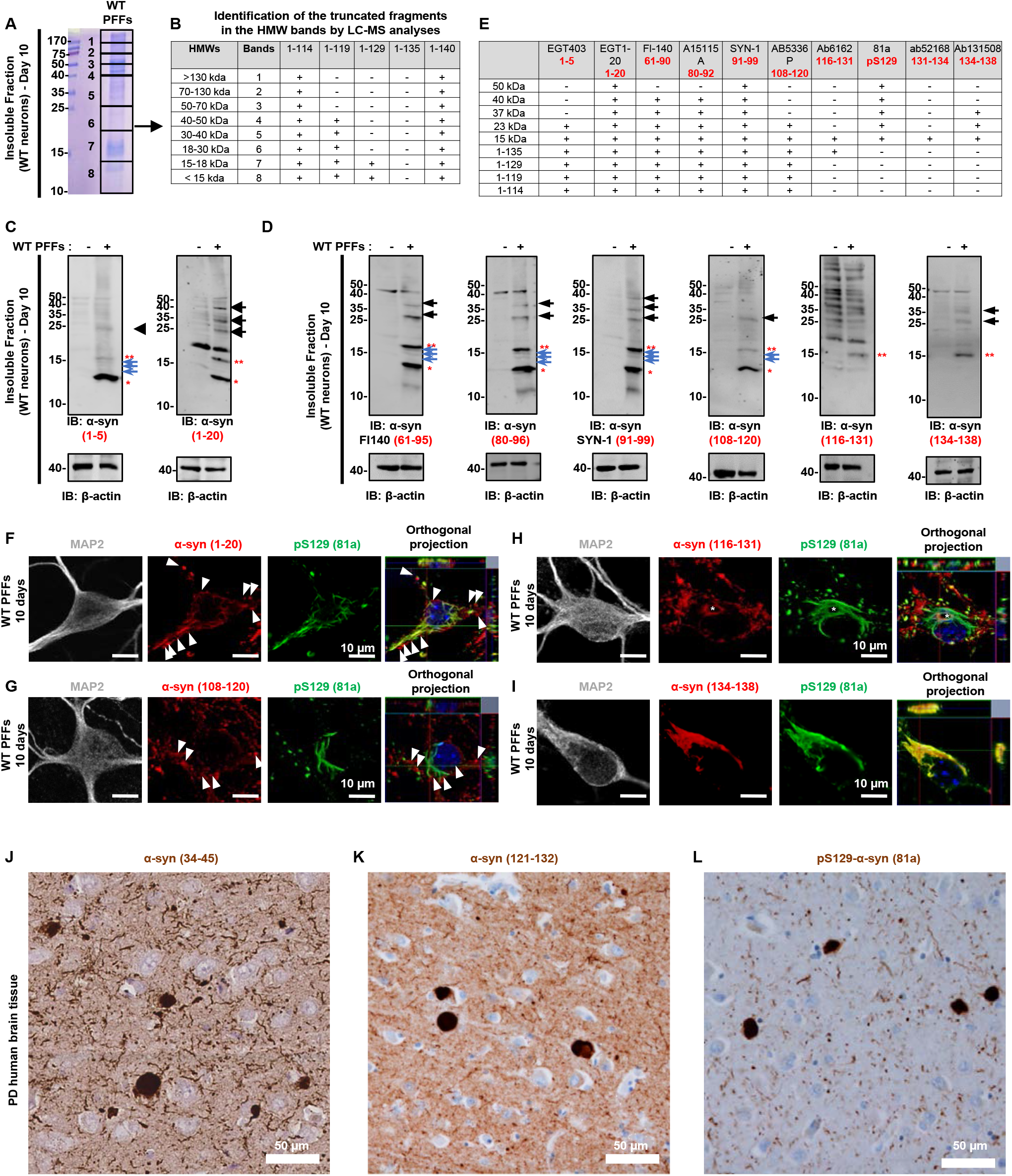
Newly formed fibrils are cleaved in WT neurons treated with PFFs^WT^ for 10 days. WT neurons were treated for 10 days with PFFs^WT^. Neurons were lysed, and the insoluble fractions were analysed by proteomic analysis **(A-B)**, WB **(C-E)**, or fixed for confocal imaging **(F-I)**. **A-B**. Insoluble fractions of α-syn WT primary neurons treated with 70 nM of mouse PFFs^WT^ for 10 days were separated on a 16.5% SDS-PAGE gel. After Coomassie staining, 8 bands were extracted from the SDS-PAGE gel (A). Isolated bands were subjected to proteolytic digestion using trypsin (to analyse C-terminal truncation^68^), followed by LC-MS/MS analysis. **B.** Proteomic analyses showed the presence of the 1-114 and 1-119 C-terminal truncated fragments in the HMW species. **C.** N-terminal antibodies raised against the residue 1-5 or the residue 1-20 (**Figure S6F**) were capable of detecting full-length (15 kDa, indicated by a double red asterisk) or truncated (~12 kDa, indicated by a single red asterisk) α-syn, but also the HMW species (indicated by the black arrows), confirming that the N-terminal region is intact in α-syn relocated in the insoluble fraction of WT neurons treated for 10 days with PFFs. Blue arrows indicate the intermediate α-syn-truncated fragments. **D.** Mapping of the C-terminal cleaved product using antibodies raised against the NAC and the C-terminal domain of α-syn (**Figure S6G**). The 1-114 fragment was well detected by the NAC antibodies and by the C-terminal antibody against residues 108-120 in the insoluble fraction of WT neurons treated with PFFs^WT^ for 10 days. Antibodies with epitopes further than the residue 116 could not detect the cleaved fragments formed in these neurons. **E**. Table summarizing the capacity of NAC, N-, and C-terminal antibodies to detected full-length α-syn (15 kDa), C-terminally cleaved fragment of α-syn (~12 kDa), and the HMW formed in WT neurons after 10 days of treatment with PFFs^WT^. **F-I**. Antibody mapping of the newly formed inclusions using pS129 antibody (81a clone) in combination with N-terminal (**F**, epitope 1-20) or C-terminal (**G-I**, respective epitopes [108-120], [116-131], or [134-138]) antibodies revealed the presence of α-syn-positive aggregates that were not pS129 positive **(G)** or only partially phosphorylated at S129 residue **(H-I)**. Scale bars = 10 μm. **J-L**. Eight-micrometer-thick paraffin-embedded serial sections from the cingulate cortex of PD human brain tissue were stained with the α-syn N-terminal antibody (J, epitope: 34-45), C-terminal antibody (K, epitope: 121-132) or pS129 antibody (L, 81a). Scale bars = 50μm.

We next assessed the presence of the C-terminal truncated fragments in the HMW species observed in WT neurons after 10 days of treatment with mouse PFFs^WT^. Interestingly, mass spectrometry analyses revealed the presence of the 1-114 fragment along with full-length α-syn in the HMW aggregates in WT neurons treated for 10 days with mouse PFFs^WT^ (Figures 4A-B). The HMW species above 130 kDa, however, were mainly composed of the endogenous α-syn (Figure S8C). These data suggest that the newly formed fibrils are prone to C-terminal cleavage. WB analyses using antibody mapping confirmed the presence of C-terminal truncated species in the HMW bands (Figure 4D). The HMW species were mainly detected by antibodies raised against the N-terminal and NAC regions (Figure 4D). Among the C-terminal antibodies tested, the ones against 108-120 still revealed the HMW bands at 23 kDa, and those against 134-138 revealed the HMW bands at 23 and 37 kDa. Antibody specific to the region 116-131 did not allow to detect specifically the HMW aggregates at 23, 37, 40, and 50 kDa. Conversely, pS129 antibody can detect the HMW species at 23, 37, 40, and 50 kDa (Figure 1F). Overall, our data demonstrate that the HMW aggregates are composed of both full-length and truncated fragments of α-syn, which supports the hypothesis that C-terminal truncation also occurs during aggregate formation or post-fibrillization.

To further confirm these findings, we compared the immunoreactivity of α-syn aggregates by ICC using a set of antibodies targeting the N- and C-terminal regions of the proteins (Figure S6E). The use of different antibodies allowed the detection of different sub-populations of α-syn-positive aggregates in WT neurons after 7 or 10 days of treatment with PFFs^WT^ (Figures 4F-I and S8D-E). After 7 days of treatment, N- and C-terminus antibodies allowed the detection of all the pS129-positive aggregates inside the neurons (Figures S8D-E). However, after 10 days of treatment, the N-terminal antibody and C-terminal antibodies raised against the 108-120 or the 116-131 region uncovered the presence of α-syn-positive accumulations that were not pS129 positive (Figures 4F-G) or only partially phosphorylated at the S129 residue (Figure 4H). These sub-populations of aggregates were either localized near the pS129-positive inclusions (Figure 4F-G, white arrows) or inside the pS129-positive filamentous structures (Figure 4H, white asterisk). As expected, the C-terminal antibody (aa 134-138) that recognizes only full-length α-syn by WB (Figure 4D) detected the aggregates that were pS129 positive (Figure 4I).

Antibody mapping demonstrates that antibodies raised against peptides bearing amino acids after residue 116 do not detect a large proportion of the C-terminally-cleaved α-syn species present in the insoluble fraction of the primary neurons or in post-mortem human brain tissue (Figures 4D-I). This occurs because the respective antigens appear to be removed during the cleavage of α-syn fibrils in neurons. Consistent with this hypothesis, the use of antibodies targeting full-length and truncated α-syn species led to the identification of different types of α-syn-positive aggregates inside the PFF-treated neurons.

To validate the pathophysiological relevance of our findings, we next stained individual serial sections of brains from patients with sporadic PD (cingulate cortex, Figures 4J-L; substantia nigra, Figures S9A-B), with *SNCA* G51D mutation (pons and cingulate cortex, Figures S9 C and D respectively) or with MSA (Figure S9E) using antibodies recognizing α-syn phosphorylated at residue S129 or the NAC, the N-or the C-terminal domain of α-syn (Figure S6E). Strikingly, this set of antibodies revealed different types of α-syn inclusions present in the patient’s brain tissues. In the cingulate cortex of PD patients, the N-terminal antibody (epitope: 34-45) allowed the detection of the LBs, the LNs and the Lewy dots next to synaptic α-syn. As expected, the C-terminal antibody (epitope: 121-132) and the pS129 (81a) antibody, commonly used to characterize synuclein pathology, recognized the LBs but were less efficient in staining the LNs and dots (Figures 4J-L). In the substantia nigra of PD patients, the NAC (epitope: 80-96) and the 4B12 (epitope: 103-108) antibodies showed a heavier pathology load with more numerous and larger structures detected than the LB509 (epitope: 115-122) and the pS129 (81a) antibodies (Figure S9A). These results demonstrate that the use of a set of antibodies allowed us to capture the morphological spectrum of α-syn pathology in various brain regions of patients with sporadic PD. In the *SNCA* G51D pons and cingulate cortex, all antibodies showed positivity for neuronal cytoplasmic inclusions, GCI-like inclusions and threads (Figures S9C-D). In the MSA pons and putamen, the antibodies against N-ter (epitopes: 1-20 and 34-45) and C-ter (epitope: 121-132) of α-syn and against α-syn pS129 (81a) recognized frequent GCIs and threads, as well as neuronal nuclear and cytoplasmic inclusions (NNIs and NCIs, respectively) (Figure S9E). Altogether our results highlight the importance of developing new tools in addition to the pS129 or C-terminal antibodies that target residues 116 to 140 to monitor and quantify α-syn aggregation and the spread of pathology.

### Consequence of post-fibrillization C-terminal truncations of newly formed fibrils: disruption of C-terminus-mediated α-syn protein-protein interactions

Increasing evidence suggests that the C-terminal domain and/or PTMs within this domain play an important role in regulating the processing and clearance of α-syn fibrils by the chaperones and the degradation machinery. Therefore, based on our findings, we hypothesized that the C-terminal cleavage of the newly formed aggregates could result in the loss of the C-terminal partners in α-syn inclusions overtime. Toward testing this hypothesis, we first identified the interacting partners of intact recombinant PFFs by incubating cortical brain mouse lysates with PFFs labelled with biotin (Figure S10A-F). Biotinylated fibrils were then specifically pulled down using streptavidin beads, and putative interacting partners were determined by LC-MS/MS analysis (Figures S10G, I). These conditions allow the analysis of α-syn fibril interaction with partners independently of the truncation process and/or the formation of the LB-like inclusions. As expected, ~80% of the previously reported putative α-syn C-terminal interacting partners (Figure S10G-I) were found to bind to full-length PFFs in the pull-down assay (Figure S10G).

We next conducted quantitative proteomic studies on the insoluble fraction of PFF-treated neurons. We monitored the changes in the levels of the 76 previously reported C-terminal α-syn interacting proteins over time (Figure S10H) and determined which C-terminal interacting proteins were lost under conditions where truncations were detected in the neuronal seeding model. After 14 days of treatment, ~25% of the proteins identified as putative C-terminal interactors were significantly enriched in the insoluble fraction of the PFF-treated neurons. After 21 days, only 9% of these C-terminal interactors were still present in the insoluble fraction of the PFF-treated neurons (Figure S10I). These findings are in agreement with our hypothesis that C-terminal truncations of α-syn occur post-fibrillization and lead to the disruption of the α-syn interactome involving the C-terminal region of the protein. This mostly includes proteins involved in the cytoskeleton architecture (e.g. MAP1B, Tubb3, Tubb5, Tubb6, Myh10, Rap1a), suggesting that the remodelling of the newly formed fibrils by C-terminal cleavage leads to a loss of physical interactions with the cytoskeleton and other partners. Our results suggest that C-terminal truncations could reflect a cellular response to protect against aberrant C-terminal interactions in the fibrillar state or an active process related to the remodelling of fibrils and formation or maturation of LBs.

### C-terminal truncation promotes the lateral association of α-syn fibrils *in vitro*

To investigate the impact of C-terminal truncations on the morphology of a-syn fibrils, we initially assessed the structural properties of fibrils formed after the incubation of full-length α-syn monomers (1-140) or C-terminal truncated α-syn monomers (1-135, 1-133, 1-124, 1-120, 1-115, or 1-111) at 37°C under agitation conditions (Figures 5A-B). After 6 days, EM imaging showed that the PFFs^1–140^ and PFFs^1–135^ formed predominantly well-dispersed single fibrils, whereas PFFs formed by monomers derived from shorter C-terminally truncated fragments (PFFs^1–124^, PFFs^1–120^, PFFs^1–115^, and PFFs^1–111^) exhibited a high tendency to associate laterally and pack together (Figures 5B), forming micrometer-large dense aggregate clumps. The PFFs^1–133^ exhibited an intermediate behaviour with less dispersed fibrils compared with PFFs^WT^ (Figure 5B). As expected, single (Δ111-115, Δ120-125 or Δ133-135) or double deletions (Δ111-115Δ133-135) that do not substantially decrease the number of negative charges compared to WT α-syn did not favour the lateral association of these PFFs (Figure 5B). Interestingly, well-dispersed single fibrils were observed when PFFs^1–114^ were formed at pH 10.5, which kept the number of negative charges (−14) comparable to that of the WT protein (−12) (Figure 5C). However, upon the adjustment of the pH to 7.5 (corresponding to the isoelectrical point of α-syn 1-114), the PFFs^1–114^ underwent rapid lateral association and packed together in aggregate clusters. Altogether, our data demonstrate that the level of lateral association, the size, and the density of the resulting fibrils clumps appeared to depend on the charge state of the C-terminal domain, whereby monomers with the shortest C-terminal domains (i.e. the lowest number of C-terminal negative charges) led to the formation of aggregate clusters that are more clumped, larger in size, and more densely packed.

**Figure 5.**
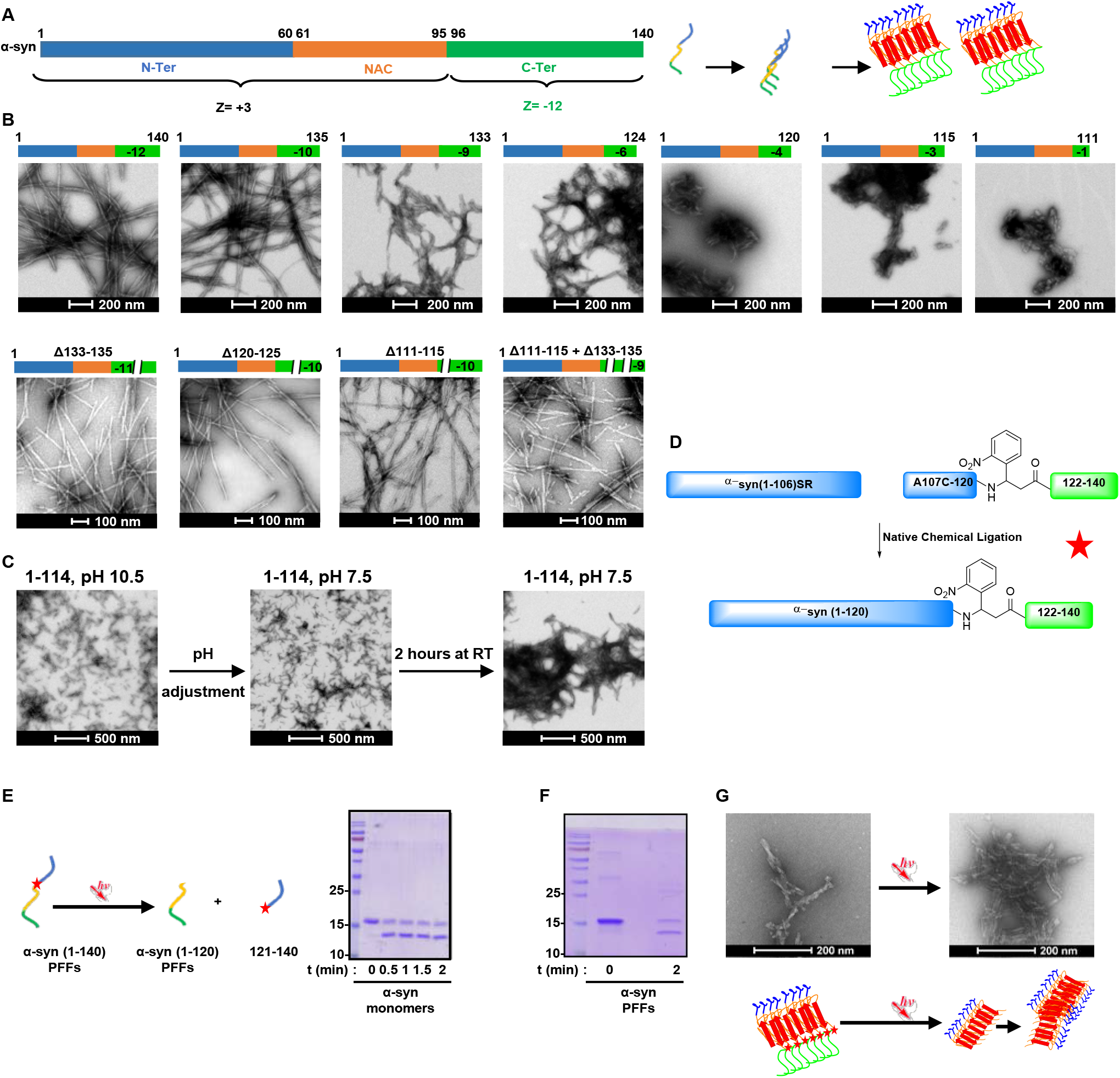
C-terminal truncation promotes the lateral association of α-syn fibrils *in vitro* A-B. Full-length recombinant α-syn, C-terminal truncated α-syn (1-135, 1-133, 1-124, 1-120, 1-115 or 1-111), and α-syn with single (Δ111-115, Δ120-125 or Δ133-135) and double deletions (Δ111-115Δ133-135) were incubated at 37°C for 6 days under shaking conditions and imaged by EM. We observed that the removal of charges in the C-terminal truncated proteins induce lateral aggregation of the PFFs, resulting in highly packed fibrils. The level of lateral association is stronger for the proteins with a lower number of negative charges in their C-termini. The number of C-terminal charges is indicated on the right side of each diagram. Scale bar = 200 nm. Lateral association is not observed for the PFFs^1–135^, for which the number of negative charges remains comparable to that of the WT protein. Likewise, PFFs will single (Δ111-115, Δ120-125 or Δ133-135) and double deletions (Δ111-115Δ133-135) do not laterally associate, as the number of negative charges remains comparable to that of the WT protein. Scale bars = 200 nm. **C.** Fibril formation was induced by incubating C-terminal truncated α-syn 1-114 at 37°C and pH 10.5 in 50 mM Tris and 150 mM NaCl under constant agitation at 1000 rpm on an orbital shaker. After 6 days, fibrils were sonicated, and the pH was re-adjusted to 7.5. After 2 hours at room temperature, fibrils laterally associated and formed large, dense aggregate clumps. Scale bars = 500 nm. **D**. Semisynthesis strategy of photocleavable α-syn at position 120. **E-F**. Photolysis of α-syn monomers **(E)** and fibrils **(F)** results in the rapid generation of C-terminally truncated α-syn. **G**. EM confirmed that photolysis of photocleavable α-syn fibrils enabled the lateral association and clumping of the fibrils. Scale bars = 200 nm.

To assess the effect of site-specific post-fibrillization cleavage on fibril morphology and lateral association with great precision, we developed a novel semisynthetic form of α-syn with a (2-nitrophenyl)propanoic acid between residues 120 and 122 (α-syn-D121Anp) allowing the temporal regulation of α-syn cleavage site-specifically. The incorporation of this unnatural amino acid allows the photocleavage of α-syn specifically at position 120 (Figure 5D). We first performed cleavage of the monomeric α-syn-D121Anp using ultraviolet (UV) light, and the results showed that cleavage occurred within a few seconds (Figure 5E). Next, we prepared α-syn-D121Anp PFFs (Figure 5F) that were successfully cleaved following exposure to UV light. EM imaging confirmed that the site-specific C-terminal photocleavage of full-length PFFs and removal of the last 20 amino acids led to their tight lateral association and the formation of fibrils that resemble those seen with the C-terminal truncated proteins (Figure 5G).

### Newly formed fibrils undergo lateral association and fragmentation over time

Having shown that C-terminal truncation promotes the lateral association of α-syn fibrils *in vitro,* we next sought to determine if this process indeed occurs in neurons. Using CLEM imaging, we followed the structure and arrangement of α-syn newly formed fibrils at the ultrastructural level in neurons treated for 7, 10, 14, or 21 days with WT PFFs (Figures 6 and S11). First, immunogold labelling of the pS129-positive inclusions allowed the detection of long filaments in the cytosol of PFF-treated WT neurons (Figure 6A), as observed previously^6^. We established that the width of the newly formed fibrils was smaller than the average width of the microtubules (Figure S11A), which enabled us to discriminate α-syn filaments from the cytoskeletal proteins.

**Figure 6.**
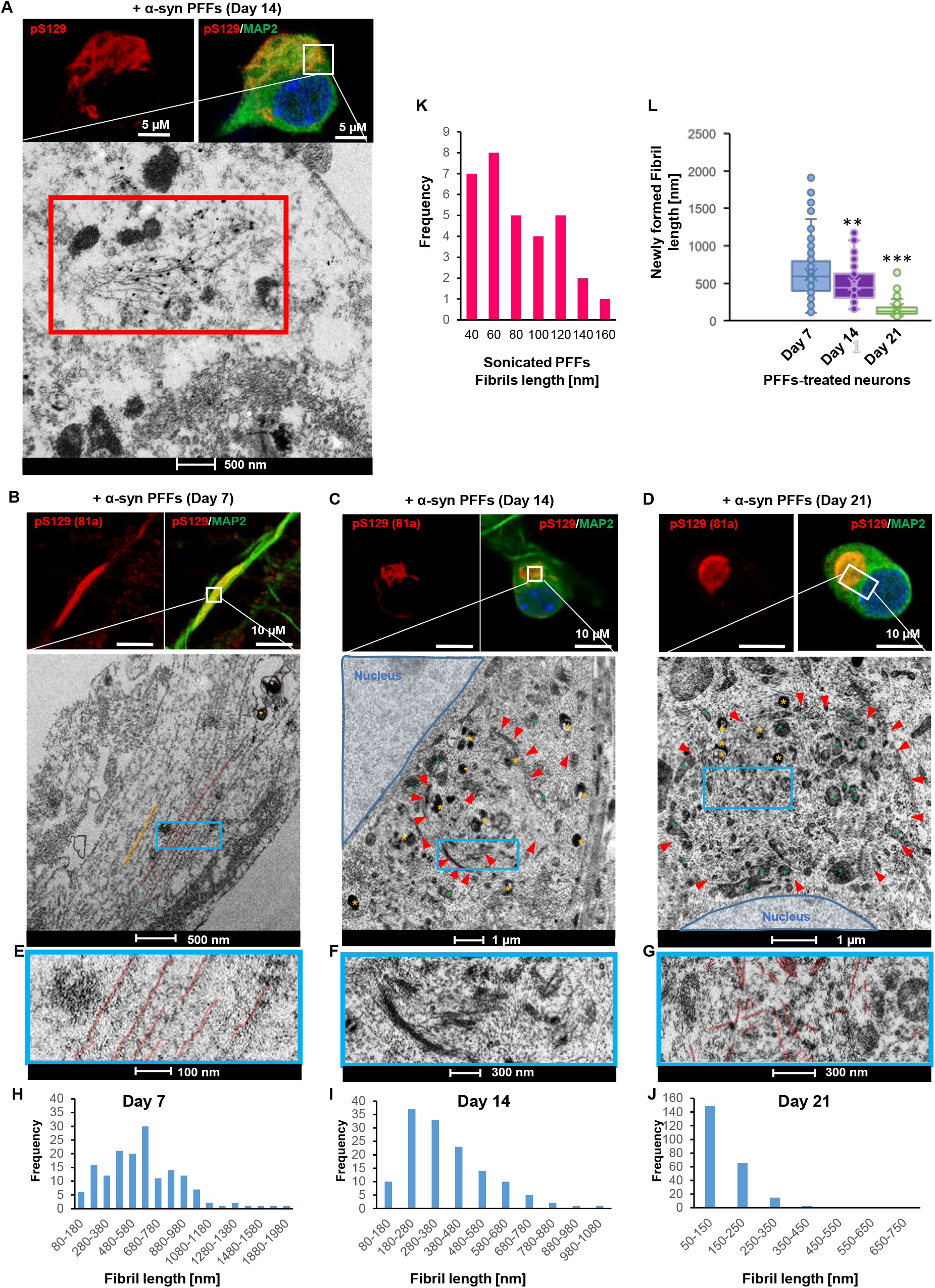
Newly formed fibrils are packed into higher-ordered aggregates. PFFs were added for 7, 14, and 21 days to the extracellular media of hippocampal neurons plated on dishes with alpha-numerical searching grids imprinted on the bottom, which allowed easy localization of the cells. At the indicated time, neurons were fixed and imaged by confocal microscopy (**A-D**, top images), and the selected neurons were embedded and cut by an ultramicrotome. Serial sections were examined by EM **(A-G)**. Immunogold labelling with the pS129 (81a) antibody confirmed that the newly formed fibrils (indicated by the red rectangle) observed in the PFF-treated neurons are positive for pS129-α-syn **(A)**. Representative newly formed α-syn fibrils are highlighted in red (**B, E, G**) or indicated with a red arrow **(C-D)**. Similar inclusion than shown in **D** can be observed in **Figure S11L**. Autophagolysosomal-like vesicles are indicated by a yellow, and mitochondrial compartments are indicated by a green asterisk. Nucleus is highlighted in blue and microtubules in orange. The length of the PFFs seeds and the newly formed fibrils was measured **(H-L)**. A minimum of 160 fibrils were counted for each condition. p<0.0001=**, p<0.0001=*** (ANOVA followed by Tukey HSD post-hoc test, Day 7 vs. Day 14 or Day 21). **A-B.** Scale bar = 500 nm; **C-D**. Scale bar = 1 μm; **E**. Scale bar = 100 nm; **F-G**. Scale bars = 300 nm.

EM imaging revealed that the length of the newly formed fibrils was significantly reduced during the maturation of the inclusions (Figures 6B-J, L, S11F-L). At Day 7, the neurites contained long filaments of α-syn that could reach up to 2 μm in length (Figures 6 B, E, H, L). These data clearly establish that the fibrils detected in the cytosol were not simply internalized PFFs (size: ~40-160 nm, Figure 6K) but truly newly formed fibrils resulting from the seeding process. Strikingly, at day 14, the size of α-syn fibrils was significantly reduced to an average length of 400 nm (Figures 6C, F, I, L and S11 F, H, J), which reinforces the hypothesis that the lateral association of the newly formed fibrils and their packing into higher-organized aggregates requires the cleavage of α-syn into shorter filaments. This was even more obvious in the compact inclusions observed in neurons treated for 21 days, in which only very short filaments were detected with an average size of 300 nm (Figures 6 D, G, J, L and S11 G, I, K-L). Our data fit well with recent findings showing that several chaperones efficiently disaggregate α-syn fibrils *in vitro* and in cells^15^ via fibril fragmentation followed by fibril dissociation into monomers.

CLEM demonstrated that the maturation of the newly formed α-syn fibrils into inclusions is accompanied by the shortening of the filaments and established that newly formed α-syn fibrils undergo a marked reorganization and fragmentation over time. Indeed, at day 7, EM images showed predominantly single fibrils aligned in parallel that stayed distant of each other in the neuronal extensions (Figures 6B, E, highlighted in red), but at day 14, α-syn filaments were completely reorganized into tightly packed bundles of fibrils (Figures 6C, F and S11F-H, red arrows). These clusters were not formed by randomly arranged fibrils but rather by fibrils that were closely aligned in parallel. This suggests that the lateral association of the newly formed fibrils might be required to start the process of inclusion formation.

Finally, we observed a redistribution of the α-syn aggregates from the neuronal extensions (day 7) to the neuronal cell bodies (day 14 and day 21), where they eventually deposited in the vicinity of the nucleus (Figures 6B-D). This suggests that over time, α-syn aggregates are all transported to the same location. Therefore, the local increase in the concentration of the newly formed fibrils might favour physical contact and clustering of the filaments into higher-ordered inclusions. In line with this hypothesis, we observed that after 21 days of treatment, neurons exhibited more compact aggregates (Figures 6D and S11L), in which α-syn newly formed fibrils were rearranged into more rounded structures (indicated by red asterisks) that started to be encircled by mitochondria (indicated by green asterisks) and autophagolysosome-like vesicles (indicated by yellow asterisks) (Figure 6D and S11L). The process of inclusion formation might not be only based on the mechanical assembly of newly formed α-syn fibrils but seems to be accompanied by the sequestration of other proteins, membranous structures, and organelles over time, as previously shown for the cortical LBs extracted from post-mortem PD brains and analysed by proteomic analyses^16^.

Altogether, our findings suggest that the formation of α-syn inclusions, in the context of the neuronal seeding model, requires a sequence of events starting with 1) the internalization and cleavage of PFF seeds, followed by 2) the initiation of the seeding, 3) fibril elongation along with the incorporation of PTMs (e.g. phosphorylation and ubiquitination) and finally 4) the C-terminal truncation of the newly formed fibrils by proteases, which allows the structural reorganization of the fibrils, their lateral assembly, and their packing into higher-order aggregates (Figure 9).

### Calpains 1 and 2 are involved in the truncation of PFFs and regulate α-syn truncation during inclusion formation in primary neurons

Having established that α-syn C-terminal cleavage plays a critical role in the lateral association of the newly formed fibrils and their packing into higher-organized aggregates, we next investigated which proteases could be involved in this process. Previous studies have shown that calpains 1 and 2 cleave α-syn fibrils predominantly at the amino acids 114 and 122^17^ *in vitro.* Furthermore, inhibition of calpain 1 reduces α-syn truncation, aggregation, and toxicity in animal models of synucleinopathie^s18,19^. To evaluate the potential contribution of calpains 1 and 2 in regulating α-syn seeding and inclusion formation, we first investigated whether these enzymes were activated in the WT neuronal primary cultures in response to treatment with PFFs (Figures 7 and S12A-B). No significant levels of calpain 1 and calpain 2 proteins were detected in the extracellular media by WB analyses (Figures S12A-B). Additionally, no differences in calpain activity level were measured in the extracellular media of Tris or PFF-treated neurons (Figure 7A).

**Figure 7.**
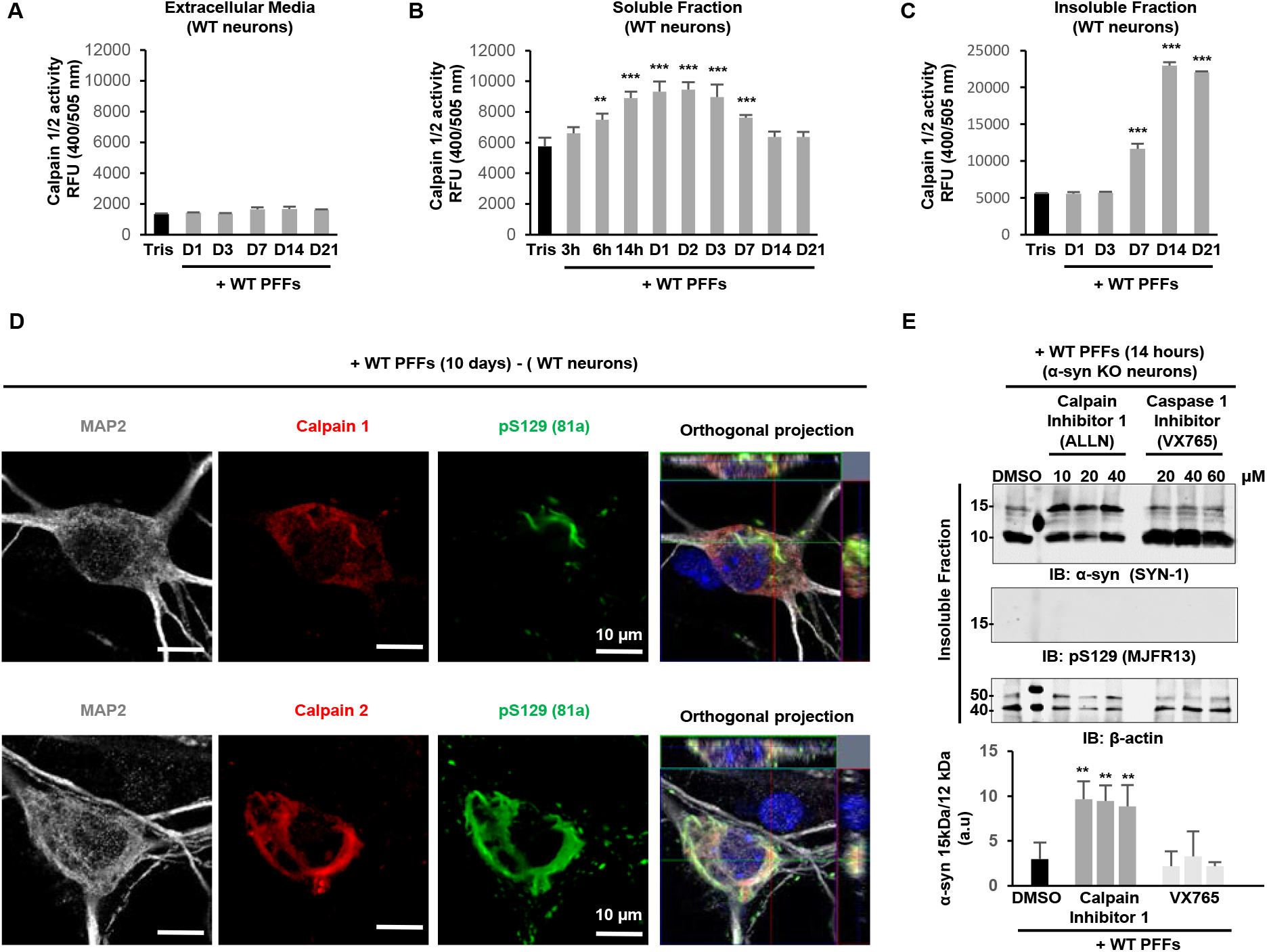
Calpains 1 and 2 are involved in the truncation of α-syn in primary neurons A-C. Calpains 1 and 2 are activated in the soluble **(B)** and insoluble fractions **(C)** but not in the extracellular media **(A)** of PFF-treated WT neurons or control neurons (Tris). Activity levels of calpains 1 and 2 were assessed at the indicated times. The graphs represent the mean +/− SD of 3 independent experiments. p<0.001=**, p<0.0001=*** (ANOVA followed by Tukey HSD post-hoc test, Tris vs. PFF treated neurons). **D.** Confocal imaging confirmed the recruitment of calpains 1 and 2 inside the LB-like inclusions positively stained for pS129-α-syn. Neurons were counterstained with MAP2 antibody, and the nucleus was counterstained with DAPI staining. Scale bar = 10 μm. **E.** α-syn truncation is blocked by calpain 1 inhibitor. α-syn KO neurons were pre-treated for 6 hours with increasing concentrations of calpain 1 inhibitor (10, 20, or 40 μM), caspase 1 inhibitor (20, 40, or 60 μM), or DMSO as a control. 70 nM of PFFs were then added for 14 hours. After sequential extractions of the soluble and insoluble fractions, cells lysates were analysed by immunoblotting. Total α-syn was detected by the SYN-1 antibody. Histogram shows the densitometry analysis from 3 independent experiments. The graph represents the mean +/− SD of 3 independent experiments. p<0.0001=*** (ANOVA followed by Tukey HSD post-hoc test, DMSO vs. enzymatic inhibitors in KO neurons treated with PFFs).

Interestingly, calpains 1 and 2 began to be significantly activated inside the neurons 6 hours after the addition of PFFs (Figure 7B). This increase in calpain activity correlated well with the cleavage of α-syn and the appearance of the ~12 kDa fragment, which starts to accumulate within the first 6 hours after the addition of PFF seeds to the neurons (Figure 1G). Moreover, the enzymatic activity for calpains 1 and 2 was still significantly upregulated in the soluble fraction of PFF-treated neurons up to 7 days post-treatment (Figure 7B).

We next determined whether the cleavage of α-syn mediated by calpains 1 and 2 was only restricted to the cleavage of soluble α-syn or whether it could also be involved in the truncation of α-syn seeds and/or the newly formed fibrils present in the insoluble fraction. To this end, we evaluated the changes in calpain 1 and 2 activities over time in the insoluble fractions and within α-syn inclusion in PFF-treated neurons (Figure 7C). Remarkably, in the insoluble fraction of the PFF-treated neurons, a significant increase in calpains 1 and 2 was first observed at day 7, which coincided with the formation of LB-like inclusions in the neurites and cell bodies, as well as the first appearance of HMW α-syn species according to WB (Figure 1F).

The enzymatic activity of calpains 1 and 2 greatly increased at day 14 and day 21 posttreatment (Figure 7C). These observations suggest that the calpains play a key role in α-syn aggregation and/or the processing of newly formed α-syn aggregates. Consistent with this hypothesis, calpains 1 and 2 were detected inside the LB-like structures, and their signals strongly colocalized with pS129 immunoreactivity (Figure 7D). Interestingly, the majority of calpain 2 was detected within the α-syn aggregates, and only low levels of the protein remained diffused in the cytosol (Figure 7D, bottom panel). LC-MS/MS analysis of the protein content from the PFF-treated neurons also revealed that calpain 2 was significantly recruited into the insoluble fraction of PFF-treated neurons from day 14 (Figure S12C). In contrast, calpain 1 was still predominantly diffused in the cytosol (Figure 7D, top panel). In line with this observation, calpain 1 was not found to be significantly enriched in the insoluble fraction of the PFF-treated neurons according to proteomic analyses (data not shown). Conversely, gene expression profiling showed a tight gene expression regulation of the key actors of PFF processing and α-syn aggregation (Figures S12E-D).

In order to validate our findings in the neuronal seeding model, we assessed the ability of the PFF seeds to be cleaved in the presence of pharmacological inhibitors of calpain 1 or calpain 2 in α-syn KO primary neurons. The data showed that the pharmacological inhibition of the enzymatic activity of calpain 1 (ALLN or PD150606) or calpain 2 (calpain inhibitor IV) in KO neurons significantly reduced the truncation level of α-syn in a concentration-dependent manner (Figure S12G). However, increasing the concentrations of the calpain inhibitors above 10 μM did not result in a significant increase in the inhibition of α-syn cleavage beyond 50% (Figure 7E). This suggests that other enzymes might be implicated in this process. Hence, we tried to block caspase 1 activity, which has been recently implicated in α-syn processing ^20^. However, inhibiting caspase 1 activity (with VX765) failed to prevent α-syn truncation in PFF-treated KO neurons (Figure 7E).

Altogether, our data demonstrate that both calpains 1 and 2 play roles in the C-terminal truncation of α-syn PFF seeds (Figures 7C, E, S12G), the newly formed fibrils (Figure 7C), and the soluble α-syn (Figure 7B). However, confocal imaging (Figure 7D) and proteomic analyses (Figure 12C) strongly suggest that calpain 2 might have a significantly higher contribution to the truncation of PFFs than calpain 1. This is especially interesting as it has been recently reported that calpains 1 and 2 exert opposite effects in the neurons, with calpain 2 activation being neurodegenerative and calpain 1 activation being neuroprotective ^21–23^. Thus, the activation of calpain 2 could contribute to the toxicity observed from day 21 in the seeding model (Figures S2F-H).

### Truncation of α-syn is required for inclusion formation and maturation in cells

To determine if truncation of the endogenous monomeric protein is required for the formation and maturation of inclusions, we compared the capacity of the cells to form aggregates when soluble endogenous α-syn cannot be cleaved. HeLa cells overexpressing similar protein levels (Figure 8A) of either WT α-syn or the D115A α-syn mutants, which cannot be cleaved into the 1-114 fragment, were treated with human WT PFFs for 48 hours. ICC revealed that pS129-positive inclusions were formed in both the HeLa cells overexpressing WT (HeLa^WT^) and D115A (HeLa^D115A^) α-syn. The levels of aggregates were quantified based on their shapes (Figure 8D), which showed that most of the aggregates in HeLa^WT^ appeared as large and filamentous inclusions (red arrows in Figures 8B and S5B). However, small and rounded inclusions were mainly detected in HeLa^D115A^ cells (Figure 8C, green arrows).

**Figure 8.**
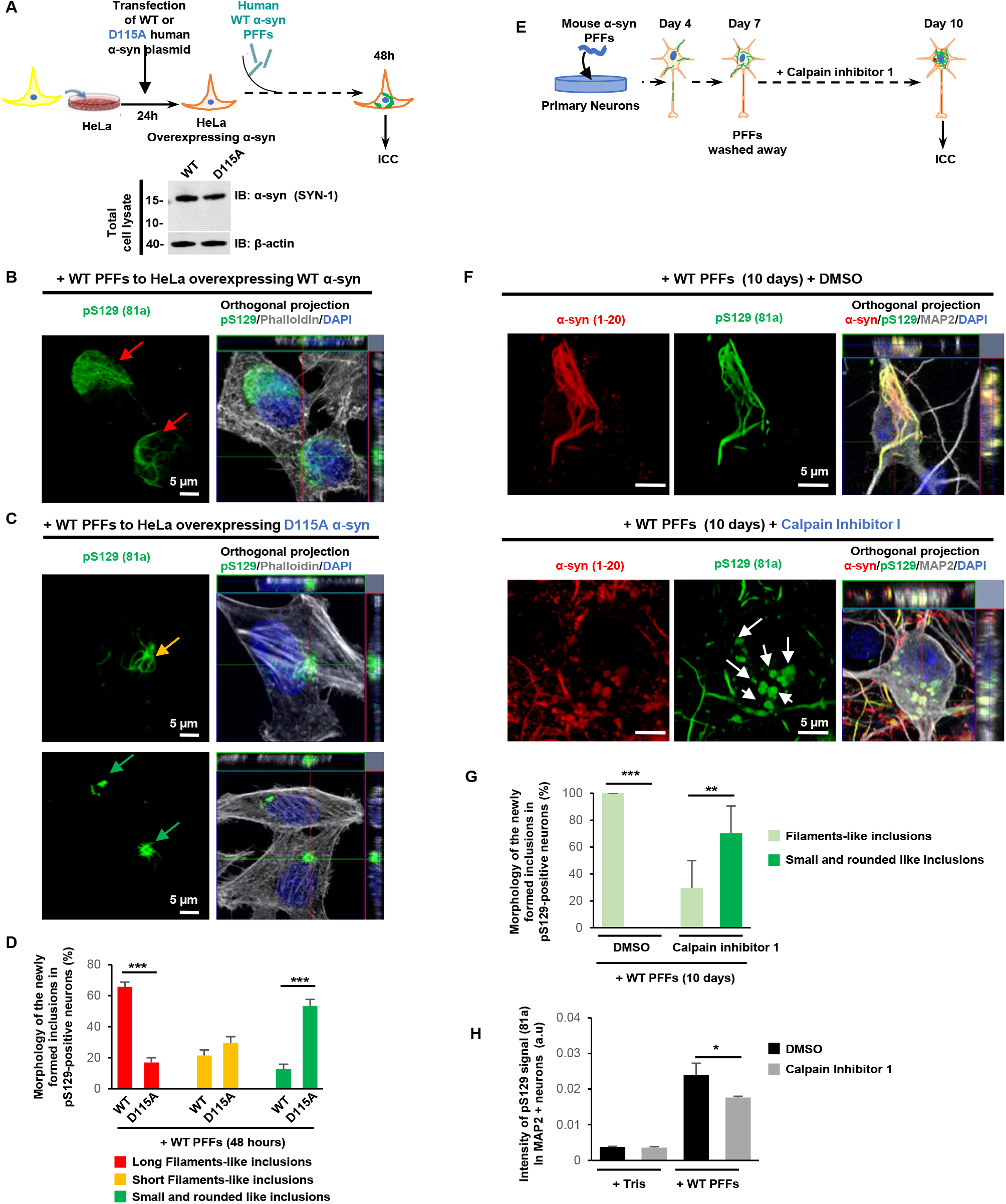
Truncation of α-syn is required for inclusion formation and maturation in cells. **A.** 24 hours after transfection with plasmids coding for WT α-syn (HeLa^WT^) or D115A α-syn mutant (HeLa^D115A^), HeLa cells were treated with 500 nM of human WT PFFs for 48 hours. **B-D**. Level and morphology of pS129-positive inclusions formed in HeLa^WT^ or HeLa^D115A^ were quantified from ICC and confocal imaging. Newly formed aggregates were classified based on their shapes in 3 groups: long filamentous inclusions (**B**, red arrows), short-filamentous inclusions (**C**, yellow arrows), or small and rounded inclusions (**C**, green arrows). Each experiment was independently reproduced at least 3 times. A minimum of 260 cells were counted for each condition. Scale bars = 5 μm. **E.** DMSO or 10 μM of calpain inhibitor I was added to PFF^WT^-treated WT hippocampal neurons from day 7 to day 10. **F-H.** Level and morphology of the pS129-positive inclusions formed in PFF-treated WT neurons in the presence of DMSO or calpain inhibitor I were assessed after immunostaining using pS129 (81a), total α-syn (1-20), and MAP2 (neurons) antibodies. Nucleus were counterstained with DAPI staining. Each experiment was reproduced at least 3 times independently. **F**. Representative images by confocal imaging. Scale bars = 5 μM. **G**. Newly formed aggregates were classified based on their shapes in 2 groups: long filamentous inclusions (**F**, top panel) or small and rounded inclusions (**F**, bottom panel). A minimum of 60 neurons were counted for each condition. **H**. Quantification of images acquired by a high-throughput wide-field cell imaging system. For each independent experiment, duplicated wells were acquired per condition, and nine fields of view were imaged for each well. Each experiment was reproduced at least 3 times independently. Images were then analysed using Cell Profile software to identify and quantify the level of LB-like inclusions (stained with pS129 antibody, 81a clone) formed in neurons (MAP2-positive cells). The graphs **(D, G-H)** represent the mean +/− SD of three independent experiments. p<0.01=*, p<0.0001=**, p<0.0001=*** (ANOVA followed by Tukey HSD post-hoc test, HeLa^WT^ vs. HeLa^D115A^ or DMSO vs. calpain inhibitor I-treated neurons).

Thus, a point mutation that prevents α-syn cleavage at the monomeric level did not prevent the initiation of the seeding, but it had a significant impact on the growth and morphology of α-syn fibrils. This reinforces the contribution of the C-terminal truncation events in the processing of fibrils and inclusion maturation. Next, we investigated the extent to which C-terminal truncation is required in the maturation of α-syn fibrils into inclusions in the neuronal seeding model. Calpain inhibitor I was added to PFF^WT^-treated WT neurons from day 7 (when the newly formed filaments are not yet cleaved, Figures 6B and S8D-E) until day 10 (Figure 8E). The shape and morphology of the aggregates were assessed by confocal imaging. Strikingly, upon treatment with calpain inhibitor I, the morphology of the aggregates detected in PFF^WT^-treated WT neurons was reshaped into small and rounded inclusions (Figure 8F). However, in the control neurons, newly formed fibrils were organized as long and/or compact filamentous structures (Figure 8G). As expected, the C-terminal antibody (epitope: 134-138) detected the small and rounded inclusions, which were pS129 positive (Figure S13A).

The C-terminal antibody raised against the 116-131 region mainly detects α-syn-positive aggregates that are not pS129 positive or only partially phosphorylated at S129 residue (Figure 4H). Interestingly, under calpain inhibitor I treatment, it was able to detect all the small and rounded pS129-positive inclusions (Figure S13B). This confirms that the small and rounded inclusions mainly contain full-length α-syn. These results demonstrate that C-terminal truncation is not only a key regulator of the formation and growth of α-syn fibrils but also plays a key role in regulating the fibril interactome, their efficient packaging, and their sequestration into mature inclusions.

## Discussion

The presence of fibrillar α-syn as the main component of the LBs has led to the hypothesis that α-syn aggregation is the primary event driving LB formation and PD pathogenesis. *In vitro* biophysical studies have provided insight into the mechanisms of α-syn seeding and fibril formation, but very little is known about the mechanisms that underlie the formation and maturation of α-syn LB and pathological inclusions in synucleinopathies^24^. Several α-syn PTMs are consistently associated with LB and pathological inclusions, which suggests that these modifications are either markers of pathology or play important roles in initiating α-syn misfolding, aggregation, and formation of pathological inclusions inside the neurons. However, several studies on the role of PTMs in regulating α-syn aggregation *in vitro* and in different synucleinopathy models by our group and others have shown that the great majority of these modifications either inhibit or do not affect α-syn fibril formation, with the only notable exception being C-terminal truncations^13,25–27^. Combined with the fact that the pathological aggregates in synucleinopathies are enriched in modified forms of α-syn, these observations led us to hypothesize that α-syn PTMs occur post-fibrillization and may play important roles in orchestrating the processing of the fibrils and/or the formation of the LBs. To test this hypothesis, we performed in-depth investigation of the roles of the two most common PTMs found in LBs and reported to play important roles in regulating α-syn aggregation and inclusions formation: phosphorylation at residue S129 and C-terminal truncations. This was accomplished using a well-established neuronal model where α-syn seeding and formation of LB-like aggregates are induced upon the addition of a small amount (70 nM) of preformed fibrils^3^.

### Cleavage of α-syn PFF seeds is a general phenomenon that occurs in different cellular seeding models

Although C-terminal truncated forms of α-syn have been observed in healthy adult human brain tissue^28^, their levels were found to increase significantly in brains from P^D13,25,27^, DLB^29^, and MSA^30–32^ patients, as well as brain tissues from Alzheimer’s disease (AD) patients without LB pathology^33^. In some studies, biochemical characterization of LB and GCI revealed that more than 90% of the α-syn detected in these inclusions is truncated^30^. Truncations of α-syn have also been reported in transgenic mouse models^34^ and their formation has been linked to increased toxicity in anima^18,35–42^ and cellular models of synucleinopathies^43–45^. However, the role of the C-terminal truncations in regulating α-syn aggregation has been studied almost exclusively in the context of its impact on the aggregation of monomeric α-syn, with an emphasis on truncation as potentially the primary trigger of α-syn aggregation, seeding and LB formation^27,29^. This working hypothesis is based on the consistent observation that C-terminal truncations dramatically increase the propensity of monomeric α-syn to aggregate *in vitro*^46–51^ and that fibrils derived from C-terminal truncated α-syn fragments seed the aggregation of full-length α-syn *in vitro*^17,50,52^ and *in vivo*^35,45,50,52,53^. To the best of our knowledge, the present work is the first attempt to dissect the role of post-fibrillization C-terminal cleavage of α-syn in the seeding, fibril formation and the evolution and maturation of LB-like aggregates in neurons.

In this work, we show that α-syn seeds are rapidly and efficiently cleaved at specific C-terminal sites after their internalization in mammalian cell lines, different types of rodent neurons, animal models, and iPSC-derived dopaminergic neurons (Figure 2). This led us to investigate the role of these truncations in regulating the initiation of the α-syn seeding process. Our results clearly demonstrate that preventing α-syn truncation, at the seed level, does not interfere with the seeding capacity of α-syn in neurons (Figure 3D-F). Notably, α-syn C-terminal truncation was not reported in the initial studies using the seeding models in mammalian cell lines and primary neurons^6,9,10,54–57^. This can be explained by the fact that the antibodies used in these studies target epitopes in the region beyond residue 114, which limited the detection to full-length or C-terminally intact α-syn species^7,9,10,54–56^ (Figure S4A). Our results are supported by recent studies which show that the truncated forms of α-syn are easily detected in seeding models when antibodies that recognize the N-terminal or the NAC region of α-syn are used, although the exact cleavage sites and role of these species in α-syn seeding and inclusion formation was not investigated in these studies^12,58^.

### C-terminal cleavage: a master regulator of α-syn aggregation and LB formation

Herein, we used biochemical, confocal, and ultrastructure imaging approaches to show that post-fibrillization C-terminal truncation plays an important role in regulating the processing and structural reorganization of the fibrils from their lateral assembly into higher-order aggregates to the formation and maturation of LBs (Figures 5–6). Our data clearly demonstrate that the newly formed α-syn fibrils in the seeding model undergo three major changes over time: 1) fragmentation evidenced by the significant decrease in fibril length over time (Figure 6); 2) increased lateral association (Figure 5 and 6); and 3) loss of C-terminal interacting proteins (Figure S10).

The fragmentation of the fibrils most likely occurs due to the action of molecular chaperones that attempt to disassemble and clear α-syn fibrils^15^ (Figure 9A). In line with this hypothesis, recent studies have shown that the human Hsp70 chaperone induces the fragmentation and disassembly of α-syn fibrils into non-toxic monomers and reduces the level of aggregates in cells^15^. This disaggregase activity is mediated by direct interactions between Hsp70 and the C-terminal domain of α-syn^15^, as evidenced by the failure of Hsp70 to disassemble fibrils prepared from C-terminally truncated α-syn variants. These observations suggest that the C-terminal domain plays a crucial role in the clearance of α-syn through the recruitment of chaperones to fibrils. Failure of the chaperone machinery or disruption of α-syn interactions with molecular chaperones would result in the accumulation of fibrils, which could lead to aberrant interactions with cytosolic proteins (Figure 9B). Under such circumstances, the sequestration of fibrils within inclusions represents an ideal detoxification mechanism through reducing aberrant interactions with fibrils. For this to happen, fibrils must be recruited and packed in high-density inclusions. When these fibrils are made of full-length α-syn, the high charge density of the C-terminus that coats the surface of the fibrils would disfavour this process unless the charge density is reduced by binding to calcium and/or other proteins or is removed by proteolytic cleavage of the C-terminal domain. These conditions would strongly favour lateral association and sequestration of fibrils into inclusions (Figure 9B). Consistent with this model, we observed an increasing degree of lateral association that correlated well with the increasing accumulation of C-terminal truncated α-syn species in the seeding model (Figure 6). Furthermore, our temporal proteomic studies showed a significant decrease in the number of C-terminal interacting proteins in the α-syn inclusions with time (Figure S10).

**Figure 9.**
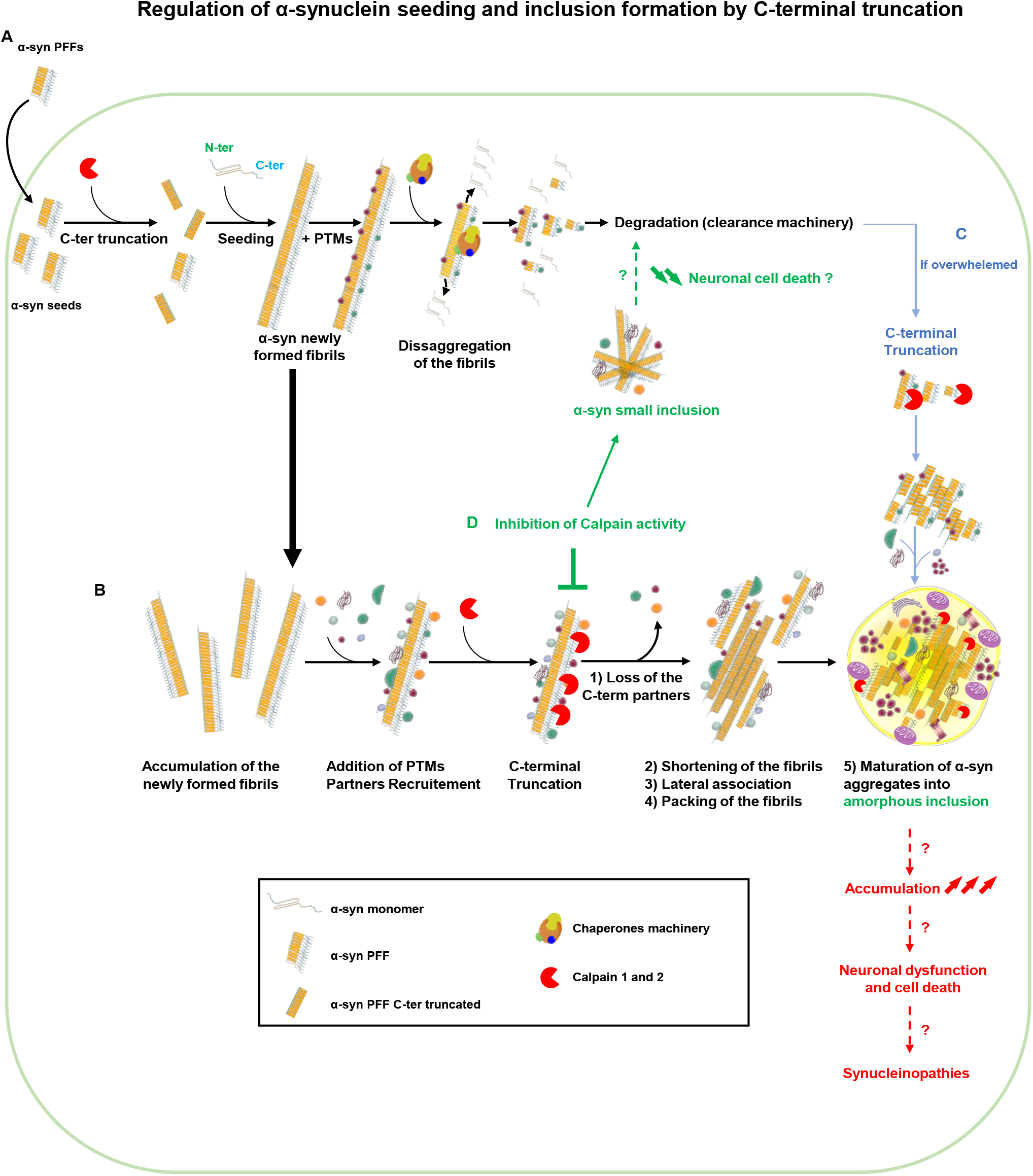
Post-fibrillization C-terminal cleavage is a master regulator of α-synuclein inclusion formation and LB biogenesis. **A**. Our findings suggest that the formation of α-syn inclusions in the context of the neuronal seeding requires a sequence of events starting with 1) the internalization and cleavage of PFF seeds followed by 2) the initiation of the seeding and 3) fibril elongation along with the incorporation of PTMs such as phosphorylation at residue S129. We hypothesized that the Hsp70 disaggregase-associated chaperones complex (Hsc70, DNAJB1 and Hsp110) can disassemble α-syn fibrils into non-toxic monomers or into shorted aggregates that can be degraded by the autophagic and/or the proteasomal pathway, which reduces the level of aggregates in cells^15^. Failure of the degradation machineries and/or disruption of α-syn interactions with molecular chaperones would result in the accumulation of fibrils in the cytosol, which in turn could lead to aberrant interactions with cellular partners. **B**. In the context of the seeding model, our data demonstrate that the C-terminal truncation of the newly formed fibrils by the proteases (e.g. calpains 1 and 2) leads to 1) the loss of the C-terminal interacting proteins, 2) the fragmentation of the filaments over time, 3) increased lateral association facilitating, and 4) the packing of the fibrils into higher-order inclusions. **C**. When the degradation machineries become overwhelmed and unable to disaggregate α-syn fibrils, C-terminal truncations could be an alternative cellular response to protect against aberrant C-terminal interactions and pack toxic fibrils into amorphous inclusions. **D**. Our data show that preventing C-terminal truncation by reducing the calpain activity disfavoured the formation of large filamentous inclusions and led to the accumulation of small inclusions in the cytosol that could be cleared more efficiently via the degradation machineries. Overall counteracting C-terminal truncation could prevent the formation, accumulation, and the cell-to-cell transmission of toxic α-syn species^4^.

Thus, under conditions where the degradation machinery fails to disaggregate and clear α-syn fibrils, the neurons would be left with no other choice but to sequester the fibrils into dense inclusions (Figure 9C). In line with this hypothesis, we have shown that the capacity of the fibrils to laterally associate and pack together is dramatically influenced by the length of the C-terminal domain and therefore the number of C-terminal negative charges carried by α-syn (Figures 5A-B) with C-terminal fragments ending at residues < 120 having the least net negative charge and exhibiting the highest degree of lateral association and packing into dense fibril networks. When these proteins are aggregated at higher pH, they form well dispersed fibrils, which undergo rapid lateral association upon shifting back to physiological pHs. Finally, using a photocleavable form of α-syn, we showed that site-specific post-fibrillization C-terminal cleavage of α-syn induces rapid lateral association of fibrils *in vitro* (Figures 5E-G).

### Calpains 1 and 2 play a key role in the packaging of the fibrils and their sequestration into mature inclusions

Having established that α-syn C-terminal cleavage plays a critical role in the lateral association of the newly formed fibrils and their packing into higher-organized aggregates, we next investigated which proteases could be involved in this process. Several enzymes, including cathepsin D, neurosin, metalloproteinases, caspase 1, calpain 1 and calpain 2 have been identified *in vitro* or *in vivo,* as potential proteases that regulate α-syn C-terminal truncations^59^. Among these, it has been shown that calpains 1 and 2 cleave α-syn fibrils predominantly at the amino acids 114 and 122^17^ *in vitro.* Furthermore, inhibition of calpain 1 reduces α-syn truncation, aggregation and toxicity in animal models of synucleinopathies^18,19^.

To evaluate a potential contribution of calpains 1 and 2 in regulating α-syn seeding and inclusion formation, we first investigated whether these enzymes were activated in the WT neuronal primary cultures in response to treatment with PFFs. The proteomic and immunofluorescence data showed the presence of calpains 1 and 2 within the LB-like inclusions formed in primary neurons (Figure 7), which is consistent with previous studies that investigated post-mortem PD and DLB brain tissues^29^. The cellular activity of calpains 1 and 2 also increased upon the internalization of α-syn PFF seeds and later during the formation and maturation of the LB-like inclusions (Figures 7 and 9B). These data are in line with previous reports showing that calpain activity is increased in dopaminergic neurons in PD^60^ as well as in the putamen and the cerebellum in MSA^61^. Further evidence in support of a central role of calpains 1 and 2 in regulating α-syn processing and PD pathogenesis comes from previous studies showing that 1) calpain-cleaved α-syn fragments are found in the core of the LBs from PD and DLB brain patients^29^; 2) calpains 1 and 2 are detected in the LBs of PD brain tissues^29,62^; 3) calpains 1 and 2 are upregulated in an 1-methyl-4-phenyl-1,2,3,6-tetrahydropyridine (MPTP) mouse model of PD, and their inhibition prevents the behavioural deficit classically observed in this model^60,63,64^; and 4) inhibiting calpain activity reduces the truncation and the aggregation level of α-syn and the motor deficit in an A30P α-syn mouse model of PD^18^.

Interestingly, transient pharmacological inhibition of calpain-mediated cleavage of endogenous α-syn monomers 7 days after addition of PFFs or preventing C-terminal truncation of the endogenous proteins at 114 (D115A mutant) does not inhibit the formation of newly formed aggregates but results in significant changes in the size and shape of these inclusions (Figure 8). Further studies are needed to understand the consequences of these changes on α-syn toxicity and maturation of LBs.

Our data and working model allow for reconciling the effects of C-terminal cleavage at different stages of α-syn fibrillization and LB formation and maturation. Under conditions where the chaperone machinery and quality control systems are operating efficiently, blocking C-terminal cleavage should prevent or reduce α-syn aggregation and promote the clearance of α-syn fibrils (Figure S14). If the chaperone and/or quality control machinery are impaired, then activation of enzymes that cleave the C-terminus provide an alternative mechanism to facilitate the sequestration α-syn aggregates in protective inclusions (Figure S14). Inhibition of post-fibrillization C-terminal truncation using calpain inhibitors or blocking C-terminal cleavage (using truncation resistant mutants) significantly altered the morphology and organization of α-syn inclusions, but did not prevent α-syn aggregation in the seeding model. These findings suggest that the C-terminal domain of α-syn plays important roles in regulating the α-syn molecular and cellular interactome during the transition from fibrillar aggregates to LBs. It is plausible that the extent of post-fibrillization cleavage and other PTMs are important determinants of α-syn fibril strains and the pathological diversity in PD, which could potentially explain the clinical heterogeneity of PD and related synucleinopathies.

### Characterization of the seeding model at multiple levels suggests that it is a good model to investigate the mechanism of Lewy body formation and maturation

An increasing number of research groups in academia and industry have used the seeding-based models to investigate the mechanisms of LB formation and pathological diversity in PD and synucleinopathies. However, systematic studies have not been performed to compare the biochemical and structural properties of the aggregates formed in these models to the pathology seen in human disease. Towards addressing this gap of knowledge, we employed an integrated approach using advanced techniques in proteomics, EM, and biochemistry to investigate the biochemical and structural properties of α-syn aggregates over time. Our extensive and in-depth characterization of the neuronal seeding model provides unique insights into the mechanism of α-syn inclusion formation and an opportunity to determine the extent to which this model reproduces the different stages of LB formation. Collectively, our results demonstrate that α-syn seeding and fibril formation can be faithfully reproduced in this seeding-based neuronal model, but not the very late stages of LB formation.

Brainstem and cortical LBs are typically described as circular inclusions with a dense core surrounded by radiating fibers^65^. In the neuronal seeding model, however, the newly formed aggregates appear as tightly packed bundles of fibrils. Interestingly, in the case of MSA, characterization of the neuronal cytoplasmic inclusions (NCIs) by EM showed that these structures are composed of long α-syn filaments that are densely packed into bundles of fibrils surrounded by mitochondria and lipofuscin vesicles^30,65^. The close resemblance between the fibrils of the NCIs and the structural organization of the newly formed fibrils in the neuronal seeding model suggest that this model might recreate more facets of MSA disease rather than PD. This is especially relevant since C-terminal truncations have also been linked to MSA, and preventing α-syn cleavage significantly reduced neurodegeneration in a rodent transgenic model of MSA^38^. However, it is important to emphasize that the rearrangement of newly formed α-syn fibrils into inclusions that are morphologically similar to *bona fide* LBs might require time and the active participation of other cellular components and machinery. Consistent with this hypothesis, our CLEM imaging clearly showed that α-syn fibrils undergo rearrangement over time to form more rounded structures that become encircled by mitochondria and autophagolysosome-like vesicles (Figures 6D and S11L). These results indicate that the proteins constituting the LBs might be recruited by different mechanisms during the maturation of the inclusions through either direct interaction with α-syn fibrils or indirectly when the organelles start to be in close proximity to α-syn. Altogether our data are in agreement with a recent model proposed by Shahmoradian et al., (preprint paper, BioRxiv)^66^, who reported that LBs from the substantia nigra and the hippocampal CA2 region from PD patients were organized structures that mainly consist of deformed mitochondria, cytoskeletal components, and lipidic structures including vesicles and fragmented membranes of organelles. This points toward impaired trafficking of organelles as a potential key driver of PD pathogenesis. Interestingly, Shahmoradian et al. reported that no radiating α-syn fibrils were found in these LBs. We propose that their findings could be explained by our results showing that the fibrils become increasingly fragmented into small fibrils over time (Figure 6L, ~50-350 nm in length at day 21 in PFFs-treated neurons) which are abundant in the final LB-like inclusions.

Such small fibrils, which are also C-terminally truncated, may not be recognized and/or could be missed, especially when C-terminal antibodies are used to localize them. It is also plausible that LBs with radiating fibrils represent only one type of LB pathology and may reflect only a subset of α-syn pathologies in PD and synucleinopathies. Indeed, several reports have confirmed the existence of a large morphological spectrum of LBs^5^. Altogether our results suggest that the neuronal seeding model recapitulates many of the key events and processes that govern α-syn seeding, aggregation and LB formation. The use of a model system that allows investigating these processes over a longer period of time (e.g: iPSC derived neurons) could enable further insights into the later stages of LB formation and maturation.

### Implications for investigating α-syn pathology formation and pathological diversity in synucleinopathies

Several antibodies are available to assess phosphorylation at S129, and an increase in pS129 signal correlates with the appearance of pathology. This has led to the use of pS129 immunoreactivity as a primary tool to assess the level of LBs in human PD brain tissue ^13^, to stage α-syn pathology in PD patients, and to quantify seeding capacity and propagation of α-syn in various animal^2^ and cellular models^3^. We report three observations that highlight the importance of new tools in addition to pS129 antibodies to assess and quantify α-syn aggregates and pathology formation. First, our findings demonstrate that C-terminal truncations play a critical role in the processes of α-syn inclusion formation and maturation. This is supported by a rich literature demonstrating that C-terminally truncated α-syn species are enriched in α-syn inclusions in PD and related synucleinopathies^13,25,27,30–33^. Together, these observations suggest that pathologies composed of truncated α-syn species would not be detected by α-syn C-terminus antibodies, including pS129 antibodies.

Second, we established that the presence of other PTMs in neighbouring residues interfere with the ability of some commercially available antibodies to detect α-syn phosphorylated at pS129 (data not shown). Third, we showed that the antibodies targeting pS129 α-syn species or the N- and C-terminal regions reveal different types of α-syn aggregates in the same PD brain tissues (Figures 4J-L, S9). These findings demonstrate that pS129 antibodies are not reliable tools for quantifying or capturing the morphological spectrum of α-syn pathology and underscore the critical importance of using multiple antibodies or unbiased approaches to accurately investigate the diversity of α-syn pathology in post-mortem tissues and in animal models of PD and other synucleinopathies.

The majority of studies on correlating the spreading of α-syn pathology and disease progression were heavily based on the use of antibodies that target the C-terminus of α-syn (Figure S4B). Thus, we argue that similar studies should be carried out again using an expanded tool box and/or integrative approaches that enable characterization of the pathology at the sequence, biochemical, and structural levels. The in-depth understanding of the human pathology would open new opportunities to investigate the relationship between the pathological diversity of α-syn and the clinical heterogeneity of PD. Such knowledge would also have significant implications in the development of imaging agents for assessing α-syn pathology, disease progression, and the efficacy of novel therapies (Figure S14).

### Implications of therapeutic strategies targeting α-syn aggregation

Although several studies have shown that the C-terminal cleavage of α-syn occurs under both physiological^28,67^ and pathogenic^25,27,32,43^ conditions, studies in cell-free systems^40,45,48–52^ and in cellular and animal models^34,36,37,39,41–43,45,50–53^ of α-synucleinopathies have consistently shown that C-terminal truncations accelerate α-syn aggregation and pathology formation. Furthermore, α-syn aggregates derived from C-terminally truncated variants of α-syn retain the ability to seed the aggregation of full-length α-syn. These observations led us to the hypothesis that C-terminal cleavage is one of the primary triggers for initiating α-syn aggregation *in vivo.* Consistent with this hypothesis, blocking or reducing α-syn C-terminal cleavage using monoclonal antibodies directed against the C-terminal part of α-syn^19^ or using inhibitors of proteases that cleave within the C-terminus of α-syn [caspase-1^20,38^ or calpain 1^18^] protected against α-syn-induced neurodegeneration and attenuated α-syn pathology spreading in animal models of PD^19^ and MSA^38^, respectively (Figure 9D). Together, these findings suggest that inhibiting C-terminal cleavage represents a viable therapeutic strategy for the treatment of PD and synucleinopathies. Since our work focused on the processing of internalized PFF seeds and processing of intracellularly α-syn fibrils and inclusions, it does not provide a clear steer on the selection of therapeutic α-syn antibodies: if α-syn antibodies actually exert intracellular effects, then C-terminally directed antibodies capable of blocking the C-terminal cleavage may be beneficial^19^ (Figure S14). However, if their main mechanism of action is neutralization of extracellular spreading species, then the key question is, which regions of α-syn are exposed to potential antibody binding in these seeds in human biofluids. It is noteworthy that systematic investigation of α-syn species in the CSF revealed that the great majority represents full-length α-syn. No significant amounts of post-translationally modified forms of the proteins could be detected (unpublished data).

The majority of the studies mentioned above and the interpretation of their findings were based on the premise that C-terminal truncations occur primarily at the monomer level and did not address the role of post-fibrillization C-terminal cleavage in LB formation and the pathogenesis of PD. Our findings, however, clearly demonstrate that significant C-terminal cleavage also occurs post-fibrillization and plays a critical role in the processing of α-syn fibrils and LB formation. Whether post-fibrillization cleavage enhances or protects against α-syn toxicity remains unclear. The answer to this question depends on whether LB formation is toxic or protective (Figure S14). As the neuronal seeding model used in this study allows only for observing the early stages of LB formation, further studies are required to optimize this model and extend the lifetime of the neurons to allow for LB formation and maturation. This would provide unique opportunities to finally understand the relationship between α-syn LB formation and neurodegeneration, and elucidate the role of PTMs in regulating the different stages of this process from α-syn misfolding to fibrillization to LB formation and clearance.

In conclusion, our work provides the most comprehensive characterization of a highly reproducible neuronal model of α-synucleinopathies and reveals novel molecular mechanisms that regulate α-syn seeding and LB formation. We demonstrate that post-fibrillization C-terminal cleavage act as master regulators of fibril processing and the formation and maturation of the LBs. We also show that the neuronal seeding model could be used as a powerful platform to identify molecular determinants and cellular pathways that regulate the different stages of LB formation, as well as to screen for therapeutic agents based on the modulation of these pathways.

## Material and Methods

### Antibodies and compounds

Information and RRID of the primary antibodies used in this study are listed in Figure S3. Tables include their clone name, catalog numbers, vendors and respective epitopes. The inhibitors of calpain 1 (Calpain inhibitor I, ALNN, #A6185; calpastatin peptide, #SCP0063) and calpain 2 (Calpain inhibitor IV, #208724) were purchased from Sigma-Aldrich (Switzerland). The caspase 1 inhibitor (VX-765, #inh-vx765i-1) was obtained from Invivogen (USA).

### Expression and Purification of human and mouse α-syn

pT7-7 plasmids were used for the expression of recombinant human and mouse α-syn in *E.coli.* Human and mouse wild type (WT) α-syn or mouse α-syn mutants (1-135, 1-133, 1-124, 1-120, 1-115, 1-114, 1-111 or 1-101, Δ111-115, Δ120-125, Δ133-135, Δ111-115Δ133-135 and S129A) were expressed and purified using anion exchange chromatography (AEC) followed by reverse-phase High-Performance Liquid Chromatography (HPLC) and fully characterized as described in detail by Fauvet et al^69^. For α-syn 1-114, the AEC purification was replaced by a cation exchange chromatography, due to the lack of the negatively charged C-terminal, then the protein was further purified and characterized similarly to the other proteins. pY125 was prepared using a semisynthesis approach as previously described^70^.

### Preparation of WT, deletion mutants and C-terminally truncated PFFs for in vitro studies

Lyophilized full length, C-terminal truncated (1-135, 1-133, 1-124, 1-120, 1-115 or 1-111) single (Δ111-115, Δ120-125 or Δ133-135) and double deletions (Δ111-115Δ133-135) α-syn proteins were dissolved in 50 mM Tris, 150 mM NaCl. The pH was re-adjusted to 7.5 using a 1M sodium hydroxyde solution and the proteins were subsequently filtered through 100 kDa MW-cut-off filters. The concentration of α-syn in solution was determined using its UV absorption at 280 nm, while, due to the low number of remaining tyrosine residues, the concentration of the C-terminal truncated α-syn mutants and α-syn deleted mutants were determined using the Pierce™ BCA Protein Assay kit (Thermo Scientific). Fibril formation was induced by incubating the proteins at a final concentration of 20 μM at 37°C and pH 7.5 (unless stated otherwise) in 50 mM Tris and 150 mM NaCl under constant agitation at 1000 rpm on an orbital shaker for 6 days.

### Preparation of WT, S129A, deletion mutants and biotinylated PFFs for primary neurons, iPSC and in vivo studies

Monomeric human or mouse WT, S129A, single (Δ111-115, Δ120-125 or Δ133-135), double deletions (Δ111-115Δ133-135), point mutations (E115A and D115A), fluorescently labelled or biotinylated α-syn were dissolved in 500 μL of Tris buffer (50 mM Tris, 150 mM NaCl, pH 7.5) filtered using a 100 kDa filter (Millipore, Switzerland) at a final concentration of 1-2 mg/mL. Due to the low number of remaining tyrosine residues, the concentrations of the C-terminal truncated and deletion mutant α-syn proteins were determined using the Pierce™ BCA Protein Assay kit (Thermo Scientific). For constituency, concentration of WT α-syn was assessed by BCA Protein Assay. Fibril formation was induced by incubating the proteins under constant orbital agitation (1000 rpm) (Peqlab, Thriller, Germany) at 37°C for 5 days^69^. After sonication with a fine probe [(4 times, 5 sec at amplitude of 20%, (Sonic Vibra Cell, Blanc Labo, Switzerland)], the biophysical and structural properties of the fibrils were assessed by EM, ThT fluorescence, SDS-PAGE and Coomassie blue staining^71^. All types of PFFs were snap-frozen in liquid nitrogen and kept at −80°C for long term storage. Of note, in the case of the α-syn 1-114 fragment, we had to adjust the fibrils preparation method to avoid clumping of the C-terminally truncated fibrils. In this particular case, the 1-114 fragment was dissolved in 50 mM Tris, 150 mM NaCl <u>at pH 10</u>. After sonication, the pH was re-adjusted to 7.5 using a solution of diluted hydrochloric acid.

### Preparation of fluorescently labelled mouse PFFs

α-syn mouse WT PFFs were diluted at a concentration of 250 μM in a final volume of 500μl of PBS. pH was adjusted to 8.5. One equivalent of Atto 488 maleimide (Atto-Tec, Switzerland) was added and incubated at 4°C overnight. The labelled PFFs were then ultracentrifuged at 100,0 *g* for 1 hour at 4°C. The supernatant was collected and the pellet re-suspended in PBS. This wash step was repeated until the dye in excess was removed. PFFs were loaded onto a SDS-PAGE gel and labelling was confirmed by scanning the gel using Typhoon FLA 7000 (GE Healthcare Life Sciences, Switzerland) with respective excitation and emission wavelength of 400 and 505 nm. Labelled fibrils were then fragmented via sonication [(4 times, 5 sec at amplitude of 20%, (Sonic Vibra Cell, Blanc Labo, Switzerland)]. PFF were then snap-frozen in liquid nitrogen and kept at −80 °C for long term storage. The biophysical and structural properties of the fibrils were then assessed by EM, SDS-PAGE/Coomassie staining and ThT assays.

### Semisynthesis, aggregation and photolysis of α-syn photocleavable at 120

The semisynthesis of α-syn photocleavable at 120 (α-syn D121Anp) was prepared as previously described^70^, we first dissolved 1-106-SR into 6M guanidine and 200 mM Na2PO4 supplemented with 30 mM TCEP. Then 1.5 equivalent of A107C-140 D121Anp peptides was added to the reaction mixture. Next the native chemical ligation was performed with shaking (1000 rpm at 37°C) for 3 hours. When the ligation was complete, as monitored my mass spectroscopy and SDS-PAGE, the protein was purified as previously described^70^. To prepare monomeric α-syn D121Anp, 100 ug of the protein was dissolved in PBS. For the preparation of fibrils, 500 ug of the protein was dissolved in 100 uL of PBS and the fibrils were formed by incubating the protein under constant orbital agitation (1000 rpm) (Peqlab, Thriller, Germany) at 37°C for 5 days^69^. The photocleavage of both the fibrils and monomer was performed using a panasonic UP50, 200 W mercury Xenon lamp, band pass filter 300-410 lamp with 100% power.

### EM

Samples (3.5 μl) were applied onto glow-discharged Formvar/carbon-coated 200-mesh copper grids (Electron Microscopy Sciences) for 1 min. The grids were blotted off, washed twice with ultrapure water and once with staining solution (uranyl formate 0.7% (w/V)), and then stained with uranyl formate for 30 sec. The grids were blotted and dried. Specimens were inspected using a Tecnai Spirit BioTWIN operated at 80 kV and equipped with an LaB6 filament and a 4K x 4K FEI Eagle CCD camera.

### HeLa cell culture, plasmid transfection and treatment with human α-syn fibrils

HeLa cells were cultured in Dulbecco’s modified Eagle’s medium (DMEM), high glucose, pyruvate and GlutaMAX™ supplemented with 10% FBS and 10 μg/mL penicillin and streptomycin in a humidified incubator, 5% CO2, 37°C. HeLa cells were transfected with a pAAV vector coding for human α-syn using lipofectamine 2000 transfectant reagent (Life Technologies, Switzerland) according to the manufacturer’s protocol. 24 hours posttransfection, human α-syn fibrils were added to the cells at a final concentration of 500 nM.

### Primary culture of hippocampal neurons and treatment with mouse α-syn fibrils

Primary hippocampal neurons were prepared from P0 pups from WT mice (C57BL/6JRccHsd, Harlan) or α-syn KO mice (C57BL/6J OlaHsd, Harlan) and cultured as previously described^72^. The neurons were seeded in black clear bottom 96 well plates (Falcon, Switzerland), 6 wells plates or onto coverslips (CS) (VWR, Switzerland) previously coated with poly-L-lysine 0.1% w/v in water (Brunschwig, Switzerland) at a density of 300,000 cells/mL. After 5 days in culture, the WT or α-syn KO neurons were treated with extracellular mouse α-syn fibrils to a final concentration of 70 nM and up to 14 days as previously describe^d6,7^.

### Intra-striatal stereotaxic injection of mouse α-syn fibrils in mice brains

3-month old C57BL/6J Rj male mice were anesthetized by intra-peritoneal injection of 100 mg/kg ketamine and 10 mg/kg xylazine. The animals were then mounted on a stereotaxic frame (model 963, Kopf, California, USA) and a lubricant eye ointment was applied. A hole was created in the right parietal bone (0.4 mm anterior and 2 mm lateral to the bregma) and 5 μg of WT PFFs or PFFs^Δ111-115^ or PFFs^1–114^ or PFFs^Δ111-115Δ133-135^ diluted in 2 μl of PBS or 2 μl of PBS as control were injected with a 34G cannula at the flow rate of 0.1 μl/min, 2.6 mm beneath the dura. The cannula remained in place for 5 min after injection. The animals were sacrificed 1, 3 and 6 months after injection by intracardiac perfusion with heparinized 0.9% sodium chloride (NaCl) followed by 4% paraformaldehyde (PFA) in PBS. The brains were post-fixed overnight in 4% PFA in PBS and paraffin-embedded. All procedures were approved by the Swiss Federal Veterinary Office (authorization number VD2067.2).

### Intra-striatal stereotaxic injection of mouse and human α-syn fibrils in mice brain

Animal experiments were performed according to the guidelines of the European Directive 2010/63/EU and Belgian legislation. The ethical committee for animal experimentation from UCB Biopharma SPRL (LA1220040 and LA2220363) approved the experimental protocol. All surgical procedures were done with wild-type male mice from C57Bl/6J and male α-syn KO mice (The Jackson Laboratory; #023837). Surgeries were performed under general anesthesia using a mixture of 50 mg/kg of ketamine (Nimatek, Eurovet Animal Health B.V.) and 0.5 mg/kg of medetomidine (Domitor, Orion Corporation) injected intraperitoneally. In addition, 2.5 mg/kg atipamezole (Antisedan, Orion Corporation) was given to support awakening. The recombinant purified full-length human PFFs were thawed and sonicated at RT by probe sonication (Q500 sonicator from Qsonica; 20 kHz; 65% power, for 30 pulses of 1 sec ON, 1 sec OFF). C57BL/6 wild-type and α-synuclein KO mice were anesthetized and stereotactically injected into the right striatum (coordinates: anterior-posterior, 0.2 mm; mediolateral, −2 mm; dorsoventral, −3.2 mm relative to bregma and dural surface) using a glass capillary attached to a 25 μl Hamilton microsyringe. 5 μg of human PFF (2.5 μg/μl in sterile PBS) were administered at a constant rate of 0.2 μl per minute and the needle was left in place for an additional 2.5-min period before its slow retraction. Mice were sacrificed at 0, 1, 3 and 7 dpi. After anesthesia, the animals were perfused through the ascending aorta with a mixed solution (30 ml/mice) made of PBS and heparin (10 U/ml, 4°C). Brains were removed from the skull and the right striatum were dissected out. Brain tissues were snap-frozen in nitrogen and stored at −80°C.

### Cell lysis and WB analyses of HeLa cells, primary hippocampal neurons and mouse-PFFs-injected mice brains

#### Primary hippocampal neurons and mouse-PFFs-injected mice brains

After α-syn fibrils treatment, primary hippocampal neurons and injected brains were lysed as described in Volpicelli-Daley *et al*.^6,7^. Briefly, treated neurons were lysed in 1% Triton X-100/ Tris buffered saline (TBS) (50 mM Tris, 150 mM NaCl, pH 7.5) supplemented with protease inhibitor cocktail, 1 mM phenylmethane sulfonyl fluoride (PMSF) and phosphatase inhibitor cocktail 2 and 3 (Sigma-Aldrich, Switzerland). After sonication using a fine probe [(0.5 sec pulse at amplitude of 20%, 10 times (Sonic Vibra Cell, Blanc Labo, Switzerland], cell lysates were incubated on ice for 30 mins and centrifuge at 100,000 g for 30 min at 4°C. The supernatant (soluble fraction) was collected while the pellet was washed in 1% Triton X-100/TBS, sonicated as described above and centrifuged for another 30 min at 100,000 g. Supernatant was discarded whereas the pellet (insoluble fraction) was resuspended in 2% sodium dodecyl sulphate (SDS)/TBS supplemented with protease inhibitor cocktail, 1 mM PMSF and phosphatase inhibitor cocktail 2 and 3 (Sigma-Aldrich, Switzerland) and sonicated using a fine probe (0.5 sec pulse at amplitude of 20%, 15 times). Mouse striatum homogenates were prepared in TBS, 1% Triton-X100 with protease inhibitor cocktail, 1 mM PMSF and phosphatase inhibitor cocktail 2 and 3 (Sigma-Aldrich, Switzerland) using a Dounce grinder. They were sonicated (20 x 1 sec at 20% amplitude, Sonic Vibra Cell, Blanc Labo, Switzerland; same condition for all following sonication steps) and an aliquot was saved as a crude homogenate. The homogenate was centrifuged (100,000 g for 60 min at 4°C) to obtain triton-soluble fractions. Pellets were vortexed in TBS, 1% Triton-X100, sonicated, and centrifuged at 100,000 g for 30 min at 4°C. New pellets were resuspended by sonication in TBS, 2% SDS (plus protease and phosphatase inhibitor cocktails) to obtain triton-insoluble fractions. Bicinchoninic acid (BCA) protein assay was performed to quantify the protein concentration in the soluble and insoluble fractions before addition of Laemmli buffer 4x (4% SDS, 40% glycerol, 0.05% bromophenol blue, 0.252 M Tris-HCl pH 6.8 and 5% β-mercaptoethanol).

#### HeLa Cells

HeLa cells were lysed in radioimmunoprecipitation assay (RIPA) buffer (150 mM sodium chloride, 50 mM Tris pH 8.0, 1% NP-40, 0.5% deoxycholate, 0.1% SDS) supplemented with protease inhibitor cocktail, 1 mM PMSF and phosphatase inhibitor cocktail 2 and 3 (Sigma-Aldrich, Switzerland). Cell lysates were cleared by centrifugation at 4°C for 15 min at 13 000 rpm. The pellet (insoluble fraction) was resuspended in SDS/TBS supplemented with protease inhibitor cocktail, 1 mM PMSF and phosphatase inhibitor cocktail 2 and 3 (Sigma-Aldrich, Switzerland) and sonicated using a fine probe (0.5 sec pulse at amplitude of 20%, 15 times). BCA protein assay was performed to quantify the proteins concentration in the soluble and insoluble fractions before addition of Laemmli buffer 4x. Proteins from the soluble and the insoluble fractions were then separated on a 16.5% or a 18% SDS-PAGE gel, transferred onto a nitrocellulose membrane (Fisher Scientific, Switzerland) with a semi-dry system (Bio-Rad, Switzerland) and immunostained as previously described^71^.

#### Cell lysis and WB analyses of human PFFs-injected mice brains and human brain tissues

The soluble and insoluble fractions of mouse and human brain tissues, for biochemistry analyses, were prepared according to previously described protocols^6,7^. Briefly, tissues were homogenized in TBS-T lysis buffer containing 50 mM Tris-HCl, 1% Triton X-100, 150 mM NaCl, and a cocktail of protease and phosphatase inhibitors (Roche), and were sonicated at 20 % power for 20 s using Q500 sonicator (Qsonica). Homogenates were centrifuged at 100,000 g for 30 min at 4 °C. The supernatant corresponded to the soluble fraction. The pellet was resuspended in TBS-T lysis buffer, sonicated at 20% power for 20 s, and then centrifuged at 100,000 g for 30 min at 4°C. The supernatant was discarded. The insoluble fraction was prepared by resuspending the pellet in TBS-T lysis buffer containing 2% SDS, and then sonicated at 20% power for 40 s. Protein concentration was determined using BCA assay. 20 μg of total protein was loaded in Tris-glycine 16% gels (Novex Thermo) and migrated at 100 V. Gels were transferred onto polyvinylidene difluoride membranes (Immobilion) at 25 V for 30 min using Trans-Blot Turbo (Bio-Rad). Membranes were blocked for 1 hour with Odyssey blocking buffer (LiCor), and then incubated overnight at 4°C with different primary antibodies diluted in the same blocking buffer. After rinsing with TBS-Tween 0.1%, membranes were incubated with IRDye^®^ conjugated secondary antibodies (1:5,000; LiCor) for 1 hour at RT, and visualization was performed by fluorescence using Odyssey CLx from LiCor. Signal intensity was quantified using Image Studio 3.1 from LiCor (RRID:SCR_013715). Primary antibodies were directed to human α-syn epitope 103-108 (4B12, 1:1,000, Thermofisher), mouse α-syn (D37A6, 1:1,000, Cell Signaling), α-syn epitope 1-20 (1:750, homemade), α-syn epitope 91-99 (clone 42, SYN-1, 1:1,000, Becton Dickinson), α-synuclein epitope 134-138 (1:1,500, Abcam), phospho-serine 129 α-syn (1:1,500, Abcam), actin (1:3,000, Cell Signaling).

#### Immunocytochemistry (ICC)

After α-syn fibril treatment, HeLa cells or hippocampal primary neurons were washed 2 times with PBS, fixed in 4% PFA for 20 min at RT and then immunostained as previously described^71^. The antibodies used are indicated in the corresponding legend section of each figure. Source and dilution of each antibody can be found in Figure S4. The cells plated on CS were then examined with confocal laser-scanning microscope (LSM 700, Carl Zeiss Microscopy, Germany) with a 40x objective and analyzed using Zen software (RRID:SCR_013672). The cells plated in black clear bottom 96 well plates were imaged using the IN Cell Analyzer 2200 (with an x10 objective). For each independent experiment, two wells were acquired per tested condition and in each well, nine fields of view were imaged. Each experiment was reproduced at least 3 times independently.

#### Human brain samples

Frozen human brain samples from MSA patients were obtained from the Netherlands Brain Bank, Netherlands Institute for Neuroscience (Amsterdam, the Netherlands; open access: www.brainbank.nl) after approval of the project by the Netherlands Brain Bank ethical committee. All material was collected from donors for or from whom a written informed consent for brain autopsy and the use of material and clinical information for research purposes had been obtained by the Netherlands Brain Bank (Figure S7F).

Human brain samples from PD, MSA and *SNCA* G51D mutation patients, collected using a London Multicentre Research Ethics Committee-approved protocol and stored under a licence issued by the Human Tissue Authority, were selected from the Queen Square Brain Bank for Neurological Disorders (QSBB) at the UCL Queen Square Institute of Neurology, University College London. This research project was approved by the procedures of the QSBB within the research tissue bank approval of the London Multicentre Research Ethics Committee.

#### Immunohistochemistry on post-mortem human tissue

Brain tissue was acquired from the Queen Square Brain Bank for Neurological Disorders, UCL Institute of Neurology, where tissue has been donated for research. The brain donation programme and protocols have received ethical approval for research by the NRES Committee London Central and tissue is stored for research under a license issued by the Human Tissue Authority (No. 12198). 3 cases of PD, 2 cases of MSA and 2 cases of *SNCA* G51D mutation was used for immunohistochemistry. Eight-micrometer-thick paraffin-embedded sections were cut sequentially. For selected primary antibodies (Figure S3A) immunohistochemical staining was performed using a Menarini Intellipath automated staining machine. Sections were de-paraffinized and treated in formic 98% acid for 10 min before pre-treatment with super access buffer using a pre-treatment unit. Sections were washed in water before staining according to the manufacturer’s instructions (A. Menarini Diagnostics, Wokingham, UK). For the remaining primary antibodies (Figure S4) sections were de-paraffinized and the endogenous peroxidase de-activated via incubation in methanol/ hydrogen peroxide (H_2_O_2_). Following 10 min pre-treatments in 98% formic acid and in citrate buffer under pressure-cooking, 10% non-fat milk in 1X TBS-Tween (Thermo Scientific, UK) was used to block non-specific binding. Tissues were incubated for 1 hour at RT with primary antibody (See Figure S4). The secondary antibody (polyclonal rabbit anti-mouse immunoglobulins, Dako #E0354, or polyclonal swine anti-rabbit immunoglobulins, Dako #E0353; 1:200) was added for 30 minutes at room temperature, followed by the application of the avidin-biotin complex (Vectastain ABC kit, Vector Laboratories, UK). Colour development was carried out using di-aminobenzidine/ H_2_O_2_, and the nuclei were counterstained using Mayer’s haematoxylin (TCS Biosciences, UK).

#### Quantitative high-throughput wide-field cell imaging screening (HTS)

After α-syn fibrils treatment, hippocampal primary neurons plated in black clear bottom 96 well plates (BD, Switzerland) were washed twice with PBS, fixed in 4% PFA for 20 min at RT and then immunostained as described above. Images were acquired using the Nikon 10X/ 0.45, Plan Apo, CFI/60 of the IN Cell Analyzer 2200 (GE Healthcare, Switzerland), a high-throughput imaging system equipped with a high resolution 16-bits sCMOS camera (2048×2048 pixels), using a binning of 2×2. For each independent experiment, duplicated wells were acquired per condition, and nine fields of view were imaged for each well. Each experiment was reproduced at least 3 times independently. Images were then analyzed using Cell profiler 3.0.0 software (RRID:SCR_007358) for identifying and quantifying the level of LB-like inclusions (stained with pS129 antibody) formed in neurons MAP2-positive cells.

#### Correlative Light Electron Microscopy (CLEM)

Mouse primary hippocampal neurons were grown on 35 mm dishes with alphanumeric searching grids etched to the bottom glass (MatTek Corporation, Ashland, MA, USA) and treated with WT PFFs. At the indicated time point, cells were fixed for 2 hours with 1% glutaraldehyde and 2.0% PFA in 0.1 M phosphate buffer (PB) at pH 7.4. After washing with PBS, ICC was performed (for more details, see the corresponding section in the Materials and Methods). Neurons with LB-like inclusions (positively stained for pS129) were selected with fluorescence confocal microscope (LSM700, Carl Zeiss Microscopy) for ultrastructural analysis. The precise position of the selected neuron was recorded using the alpha-numeric grid etched on the dish bottom. The cells were then fixed further with 2.5% glutaraldehyde and 2.0% paraformaldehyde in 0.1 M PB at pH 7.4 for another 2 hours. After washing (5 times for 5 min) with 0.1 M cacodylate buffer at pH 7.4, cells were post fixed with 1% osmium tetroxide in the same buffer for 1 hour and washed with double-distilled water before contrasting with 1% uranyl acetate water for 1 hour. The cells were then dehydrated in increasing concentrations of alcohol (2 × 50%, 1 × 70%, 1 × 90%, 1 × 95% and 2 × 100%) for 3 min each wash. Dehydrated cells were infiltrated with Durcupan resin diluted with absolute ethanol at 1: 2 for 30 min, at 1: 1 for 30 min and 2: 1 for 30 min and twice with pure Durcupan (Electron Microscopy Sciences, Hatfield, PA, USA) for 30 min each. After 2 hours of incubation in fresh Durcupan resin, the dishes were transferred into a 65°C oven for the resin to polymerize overnight. Once the resin had hardened, the glass CS on the bottom of the dish was removed by repeated immersion in hot (60°C) water, followed by liquid nitrogen. The cell of interest was then located using the alphanumeric coordinates previously recorded and a razor blade used to cut this region away from the rest of the resin. This piece was then glued to a resin block with acrylic glue, trimmed with a glass knife using an ultramicrotome (Leica Ultracut UCT, Leica Microsystems), and then ultrathin sections (50-60 nm) were cut serially from the face with a diamond knife (Diatome, Biel, Switzerland) and collected onto 2 mm single-slot copper grids coated with formvar plastic support film. Sections were contrasted with uranyl acetate and lead citrate and imaged with a transmission electron microscope (Tecnai Spirit EM, FEI, The Netherlands) operating at 80 kV acceleration voltage and equipped with a digital camera FEI Eagle, FEI).

#### Immunogold Staining

Mouse primary hippocampal neurons were grown on 35 mm dishes with alphanumeric searching grids etched to the bottom glass (MatTek Corporation, Ashland, MA, USA) and treated with WT PFFs. At the indicated time point, cultured cells were fixed for 2 hours with buffered mix of paraformaldehyde (2%) and glutaraldehyde (0.2%) in PB (0.1M), then washed in PBS (0.01M). After washing with PBS, ICC was performed as described^71^. Briefly, cells were incubated with pS129 antibody (81a, Figure S4B) for 2 hours at RT diluted in the blocking solution [(3% BSA and 0.1% Triton X-100 in PBS), (PBS-BSA-T)]. After 5 times washes with PBS-BSA-T, neurons were incubated with the Alexa Fluor^647^ Fluoronanogold secondary antibody (Nanoprobes, USA) for 1 hour at RT. Neurons were washed 5 times with PBS-BSA-T and mounted in polyvinyl alcohol mounting medium supplemented with anti-fading DABCO reagent. Neurons with LB-like inclusions (positively stained for pS129) were selected with fluorescence confocal microscope (LSM700, Carl Zeiss Microscopy) for ultrastructural analysis. The precise position of the selected neuron was recorded using the alpha numeric grid etched on the dish bottom. The cells were next washed in phosphate buffer (0.1M) and then fixed again in 2.5% glutaraldehyde before being rinsed in distilled water. A few drops of a silver enhancement solution (Aurion, Netherlands) were then added to each CS to cover all the cells and left in the dark for 1 hour at RT. The solution was then removed and the CS washed in distilled water followed by 0.5% osmium tetroxide for 30 min and 1% uranyl acetate for 40 min. Following this, the CS were dehydrated through increasing concentrations of ethanol and then transferred to 100% epoxy resin (Durcupan, Sigma Aldrich) for 4 hours and then placed up-side down on a glass CS in a 60°C oven overnight. Once the resin had cured, the CS was removed and regions of interest were cut away with a razor blade and mounted onto a blank resin block for thin sectioning. Serial thin sections were cut at 50 nm thickness with a diamond knife (Diatome, Switzerland) in an ultramicrotome (UC7, Leica Microsystems) and collected onto a pioloform support firm on copper slot grids. These were further stained with lead citrate and uranyl acetate. Sections were imaged with a digital camera (Eagle, FEI Company) inside a transmission electron microscope (Tecnai Spirit, FEI Company) operating at 80 kV.

#### Calpain 1 and calpain 2 activity in primary neurons

Using calpain activity assay kit (Abcam, UK), calpain activity was measured in primary culture treated with Tris buffer (negative control) or with α-syn WT PFFs for 3 hours and up to 21 days. At each indicated time-points, 100 μl of the extracellular media was collected before harvesting the neurons. Collected extracellular media and neurons were then lysed in the extraction buffer provided in the kit and centrifuged at 4°C at 13 000 rpm for 5 min. Soluble fractions were collected and handled following manufacturer’s instructions. Pellets from the lysed neurons were resuspended in the extraction buffer provided in the kit and mild sonication [(10 sec, 20% amplitude) (Sonic Vibra Cell, Blanc Labo, Switzerland)] was performed to ensure the complete dispersion of the pellet. Calpain activity assay was then performed in accordance with the supplier’s instructions. Fluorescein emission was quantified using Tecan infinite M200 Pro plate reader (Tecan, Männedorf, Switzerland) with respective excitation and emission wavelength of 400 and 505 nm.

#### Relative Quantification of WB and statistical Analysis

The level of total α-syn (15 kDa, 12 kDa or HMW) or pS129-α-syn were estimated by measuring the WB band intensity using Image J software (U.S. National Institutes of Health, Maryland, USA; RRID:SCR_001935) and normalized to the relative protein levels of actin. All the experiments were independently repeated 3 times. The statistical analyses were performed using Student’s t-test or ANOVA test followed by Tukey-Kramer *post-hoc* test using KaleidaGraph (RRID:SCR_014980). The data were regarded as statistically significant at p<0.05.

## Supporting information

Graphical_Abstract

## Acknowledgements

This work was supported by funding from EPFL and UCB (H.A.L), (A.L.M), (F.A), (J.B), (N.M), (N.A.B) and (A.C). R.W.-M and S.V were supported by the Joint Programme for Neurodegenerative Disease (JPND) from the UK Medical Research Council (R.W.-M.). J.H and C.S were supported by the Multiple System Atrophy Trust; the Multiple System Atrophy Coalition; Fund Sophia, managed by the King Baudouin Foundation; and Karin & Sten Mortstedt CBD Solutions. Queen Square Brain Bank for Neurological Disorders is supported by the Reta Lila Weston Institute for Neurological Studies and the Medical Research Council UK. This research was supported in part by the National Institute for Health Research University College London Hospitals Biomedical Research Centre. J.Y.L and C.H were supported by the NNSF-81430025, Swedish Research Council, EU-JPND (aSynProtec, REfrAME) and EU-ITN (Syndegen).

We acknowledge Elena Gasparotto, Jonathan Ricci and Jérémy Campos for their valuable technical assistance respectively: Elena for her continuous support with the preparation of the primary culture and cloning of mutated α-syn, Jonathan and Jérémy for the expression and purification of WT or mutants α-syn proteins and Jonathan for his help with the *in vitro* assays. We are grateful to Dr. Arne Seitz and his staff at the Bio-imaging Core Facility (EPFL) for their technical support. We thank the Protemics Core Facility (EPFL): Dr Marc Moniatte for the discussions and Dr Diego Chiappe for handling the samples. We also thank Dr Gerardo Turcatti and his staff, Dr. Marc Chambon and Dr. Fabien Kuttler, at the Biomolecular Screening Core Facility (EPFL) for the discussions and their technical support. We thank Dr Graham Knott for the discussions and Mary Croisier-Coeytaux for her technical support at the Bio-EM Core facility (EPFL).

## Author contributions

H.A.L conceived and supervised the study. H.A.L and A.L.M.M designed all the experiments and wrote the paper. A.L.M.M performed and analyzed the experiments shown in Figures 1–4, 6-9 and Figures S2-S4, S6, S7A-B, S8, S10 H-I, S11, S12C-G, S13-S14. F.A designed, performed and analyzed the experiments shown in Figures 4J-L, S5A-C and 8A-D. F.A also produced and characterized α-syn monomers and fibrils depicted in Figure S1. J.B designed, performed and analyzed the experiments shown in Figure S7C. N.M designed, performed and analyzed the experiments shown in Figures S10 G, I. N.A.B and A.C designed, performed and analyzed the experiments shown in Figure 5. A.C produced and characterized α-syn-biotin PFFs shown in Figure S10A-F. S.V and R.W.M designed Figures 2D and S5D-E. S.V performed and analyzed the experiment shown in Figure 2D and S5D-E. R.H performed and analyzed the experiments shown in Figures 3A-B and 4B-E and S6A-C. J.H and C.S performed and analyzed Figures 4 J-L and S9B-E. C.H and J.Y.L performed and analyzed Figure S9A. M.C prepared the samples for CLEM analysis and acquired EM images in Figures 6 and S11. G.K supervised the experiments shown in Figures 6 and S11 and contributed to the interpretation of the data. G.M.C designed and analyzed the experiments shown in Figure 2C, E. L.W performed and analyzed the experiments shown in Figure 2C, E. A.M designed and performed the experiments shown in Figure 2C. P.D and M.C supervised the study performed in Figure 2C, E. All authors reviewed and contributed to the writing.

