## Supplemental_Info_Figures for "The making of a Lewy body: the role of α-synuclein post-fibrillization modifications in regulating the formation and the maturation of pathological inclusions"

[1]Laboratory of Molecular and Chemical Biology of Neurodegeneration, Brain Mind Institute, Ecole Polytechnique Fédérale de Lausanne (EPFL), 1015 Lausanne, Switzerland. [2]Department of Physiology, Anatomy and Genetics, University of Oxford, Oxford, OX1 3QX, UK. [3]Queen Square Brain Bank for Neurological Disorders, Department of Molecular Neuroscience, UCL Institute of Neurology, London, UK. [4]Neural Plasticity and Repair Unit, Wallenberg Neuroscience Center, Department of Experimental Medical Science, Lund University, 22184 Lund, Sweden. [5]Institute of Health Sciences, China Medical University,110122 Shenyang, China. [6]Proteomic Core Facility and Technology Platform, EPFL, Lausanne 1015, Switzerland. [7]BioEM Core Facility and Technology Platform, EPFL, Lausanne 1015, Switzerland. [8]UCB Biopharma SPRL, Chemin du Foriest B-1420 Braine-l'Alleud, Belgium.

*To whom correspondence should be addressed at: Laboratory of Molecular and Chemical Biology of Neurodegeneration, Brain Mind Institute, Ecole Polytechnique Fédérale de Lausanne, 1015 Lausanne. Tel: +41216939691, Fax: +41216939665,

**Running title:**

C-terminal truncation: a master regulator of α-syn inclusion formation and LB biogenesis.

**Keywords:** alpha-synuclein (α-synuclein), Parkinson’s disease, aggregation, post-translational modification (PTM), truncation, phosphorylation

### Material and method

#### ***Quantification of cell death in hippocampal primary neurons***

Hippocampal primary neurons were plated in 96-well plate and treated with increasing concentrations of α-syn PFFs (70 nM to 500 nM). After 7, 14 or 21 days of treatment, cell death was quantified using complementary cell death assays.

*Quantification of cell death by dye exclusion method*

Cell death was quantified by the vital dye exclusion method using Propidium Iodide (PI, Sigma-Aldrich, Switzerland), a membrane impermeant dye that enters only in cells with damaged plasma membranes. Briefly, PI was added to the neuronal culture media at a final concentration of 2 µg/ml. As positive control, one well was fixed with 4% PFA for 15 min at RT and washed twice with PBS before the addition of PI. After 15 min of incubation, cells were washed once with PBS and fluorescence was quantified using Tecan Infinite M200 Pro plate reader (Tecan, Maennedorf, Switzerland) with respective excitation and emission wavelength of 530 nm and 620 nm. Cell toxicity was calculated for each well as follows: % cytotoxicity = 100 × fluorescence measured in live cells/ fluorescence measured in PFA-fixed well (maximum of PI fluorescence).

*Quantification of active caspase 3*

CaspaTag fluorescein caspase 3 activity kit (ImmunoChemistry Technologies, MN, USA) allows the detection of active effector caspase (caspase 3) in living cells. These kits used the specific fluorochrome peptide inhibitor (FAM-DEVD-fmk) of the caspase 3 (FLICA). This probe passively enters the cells and binds irreversibly to the active caspases. Neurons were washed twice with PBS and incubated for 30 min at 37°C with FAM-DEVD-FMK in accordance with the supplier's instructions. Fluorescein emission was quantified using Tecan infinite M200 Pro plate reader (Tecan, Maennedorf, Switzerland) with respective excitation and emission wavelength of 487 nm and 519 nm.

*<u>Quantification of LDH release</u>*

Using CytoTox 96® Non-Radioactive Cytotoxicity Assay (Promega, Switzerland), released lactic acid dehydrogenase (LDH) in culture supernatants was measured following manufacturer's instructions. After a 30 min coupled enzymatic reaction, which results in the conversion of a tetrazolium salt (INT) into a red formazan product, the amount of color formed, that is proportional to the number of damaged cells, was measured using Tecan infinite M200 Pro plate reader (Tecan, Maennedorf, Switzerland) at a wavelength of 490 nm.

***iPSC-derived neurons and treatment with human α-syn fibrils***

Induced pluripotent stem cells were derived from dermal fibroblasts. The line used in this study stems from a healthy control female of 78 years old and was reprogrammed using CytoTune-iPS Sendai Reprogramming kit (Invitrogen). The iPSC line has been characterized in previous studies as SFC856-03-04 (control 4)[1,2]. The iPSCs were kept at 37ºC, 5% $CO_2$ in feeder-free culture conditions with daily changes of mTeSR™1 (StemCell Technologies), on hESC-qualified Matrigel-coated plates (BD), and passaged as single cells using TrypLE Express incubation (Life Technologies) and ROCK inhibitor (10 µM Y-27632) (Bio-Techne, Tocris). IPSCs were differentiated into dopaminergic neurons using a floor-plate based culture established by Kriks *et al.*[3] and modified as previously described[4,5]. Cells were seeded in 6-well plates coated with Geltrex (Life Technologies) and grown to confluency. Patterning was achieved using medium with differentiation and neurotrophic factors as listed in supplemental data (Material and Method section). Medium was fully changed every two days with half change every other day until day *in vitro* 20 (DIV20). Cells were then dissociated with StemPro Accutase (Life Technologies) and re-plated onto Geltrex-coated 12 well plates (Corning) in an even monolayer of $3x10^5$ cells/cm$^2$ in medium containing ROCK inhibitor (10 µM Y-27632). Cultures were treated with 1 µg/ml mitomycin C (Bio-Techne, Tocris) in NB medium for 1 hour to remove proliferating cells and washed with NB medium before addition of fresh NB medium. After a full medium change 3 days later to remove dead cells, medium was half changed every 2-3 days until DIV50 when 70 nM of PFFs were added to neurons 14, 10, 7, 3 and 1 days before fractionation analysis at DIV64. For western blot analysis, cultures were washed in phosphate buffered saline (PBS), detached by scraping into fresh 1% Triton X-100/ Tris buffered saline (TBS) (50 mM Tris, 150 mM NaCl, pH 7.5) with protease (cOmplete™, Roche) and phosphatase inhibitors (PhosSTOP™, Roche) and subjected to a previously described fractionation protocol[6]. After sonication using a fine probe, 0.5 sec pulse at amplitude of 20%, 10 times (UP50H, Hielscher, Germany), cell lysates were incubated on ice for 30 min and centrifuged (Optima TLX ultracentrifuge, TLA-110 rotor, Beckman Coulter, USA) at 100,000 g for 30 min at 4°C and the supernatant (soluble fraction) collected. The pellet (insoluble fraction) was resuspended in 2% sodium dodecyl sulphate (SDS)/TBS supplemented with protease (cOmplete™, Roche) and phosphatase inhibitors (PhosSTOP™, Roche) and sonicated using a fine probe (0.5 sec pulse at amplitude of 20%, 15 times). 10 ug of protein from each fraction was loaded on Criterion™ TGX™ precast gel 4-15% (BioRad) and transferred onto Trans-Blot® Turbo™ polyvinylidene difluoride (PVDF) membranes (BioRad). Membranes were blocked in 5% milk in PBS containing 0.1% Tween for 30 min and incubated overnight in blocking solution with a total α-syn antibody (SYN-1, BD biosciences; 1:500). The secondary anti-mouse HRP (BioRad; 1:5000) and loading control β-actin-HRP (Abcam, 1:50 000) were diluted in blocking solution and applied for 1 hour at room temperature (RT). The membrane was exposed to Immobilon Western Chemiluminescent HRP Substrate and developed with the ChemiDoc™ System (BioRad). Protein levels were measured with the Image Lab software (BioRad) and analyzed with GraphPad Prism.

### *Medium and reagents used during differentiation of iPSC cells into dopaminergic neurons*

| Medium | Product name | Supplier | Catalogue no | Working | Time added |
|---|---|---|---|---|---|
| **KO DMEM KSR** | Knockout DMEM | Life Technologies | 10829018 | n/a | |
| | KnockOut Serum Replacement | Life Technologies | 10828010 | n/a | 100% DIV 0-4<br>75% DIV 5-6<br>50% DIV 7-8<br>25% DIV 9-10 |
| | L-Glutamine | Life Technologies | 25030-024 | 2mM | |
| | MEM NEAA | Life Technologies | 11140-050 | 1X | |
| | β-mercaptoethanol | Gibco | 21985023 | 10µM | |
| **NNB** | Neurobasal Medium | Life Technologies | 21103049 | n/a | |
| | N2 Supplement | Life Technologies | 17502048 | 0.5X | 25% DIV 5-6<br>50% DIV 7-8<br>75% DIV 9-10 |
| | B-27® Supplement w/o Vit A | Life Technologies | 12587010 | 0.5X | |
| | L-Glutamine | Life Technologies | 25030-024 | 2mM | |
| **NB** | Neurobasal Medium | Life Technologies | 21103049 | n/a | |
| | B-27® Supplement w/o Vit A | Life Technologies | 12587010 | 1X | DIV11- |
| | L-Glutamine | Life Technologies | 25030-024 | 2mM | |
| **Growth factors** | SB431542 | Bio-techne (Tocris) | 1614 | 10µM | DIV0-4 |
| | LDN193189 | Sigma | SML0559-5MG | 100nM | DIV0-10 |
| | Sonic Hedgehog C24II high activity | Bio-techne (Tocris) | 1845-SH-500 | 100ng/mL | DIV1-6 |
| | Purmorphamine | Bio-techne (Tocris) | 4551/10 | 2µM | DIV1-6 |
| | FGF8a | Stratech | 16124-HNAE-SIB | 100ng/µL | DIV1-6 |
| | CHIR99021 | Bio-techne (Tocris) | 4423 | 3µM | DIV3-12 |
| | TGFβ3 | Peprotech | 100-36E | 1ng/mL | DIV13- |
| | DAPT | Abcam | ab120633 | 10µM | DIV13- |
| | db-cAMP | Sigma | D0627-1g | 0.5mM | DIV13- |
| | GDNF | Peprotech | 450-10 | 20ng/µL | DIV13- |
| | BDNF | Peprotech | 450-02 | 20ng/µL | DIV13- |
| | Ascorbic acid | Sigma | A4544-25G | 0.2mM | DIV13- |

### *Identification of N- and C-terminal truncations sites by quantitative proteomic analyses*

After α-syn WT PFF treatment, primary hippocampal neurons (treated for 14 or 21 days) were lysed as described in Volpicelli-Daley *et al.*[6,7]. and separated by SDS-PAGE on a 16.5 % polyacrylamide gel, which was then stained with Coomassie Safestain (Life Technologies). Each gel lane was entirely sliced and proteins were In-gel digested as previously described[8] skipping the reduction and alkylation procedure. Peptides were desalted on stageTips[9] and dried under a vacuum concentrator. For LC-MS/MS analysis, resuspended peptides were separated by reversed phase chromatography on a Dionex Ultimate 3000 RSLC nano UPLC system connected in-line with an Orbitrap Lumos (Thermo Fisher Scientific, Waltham, USA). Raw data was processed using Mascot (Matrix Science, Boston, USA), MS-Amanda[10] and

SEQUEST in Proteome Discoverer v.1.4 (RRID:SCR_014477) against the Uniprot Mouse protein database. Data was further processed and inspected in Scaffold4 and spectra of interest were manually validated.

#### *Preparation of biotinylated α-synuclein monomers*

Recombinant FL M1C α-syn (10 mg) was placed in 1.0 mL of degassed buffer (200 mM Tris, 6.0 M Gdn at pH 6.5). To make sure that cysteins were in the reduced form, the protein was treated with 1.0 eq of TCEP and incubated at 37°C for 20 min. The reaction was monitored by LC-MS to confirm that dimers were reduced to monomers. Then 3.0 eq of biotin maleimide (10 uL of 200 mM solution in DMF) was added and stirred at 37°C for 60 min at pH 6.5. The progression of the labelling was monitored by ESI-LC/MS and after complete consumption of the starting M1C α-syn protein, the reaction was quenched by lowering the pH from 6.5 to 5. The reaction mixture was cooled down to 5°C and purified by C8 semi-preparative column chromatography using 20%-80% MeCN gradient over 60 min using 3.0 mL/min flow rate. The purity of the fractions was verified by UPLC and the biotin-labelled α-syn was lyophilized. The purity and identity final product was confirmed by UPLC, SDS-PAGE and ESI-LC/MS analyses.

#### *Immunohistochemistry of the substantia nigra of a PD patient*

The substantia nigra of a 69 year-old, female PD patient with Braak stage 5-6 was obtained from Department of Neuropathology, Lund University Hospital, under the ethical permit of Lund University, Sweden. 3 to 5 µm thick paraffin-embedded human brain sections from the substantia nigra of a PD patient were dried at 60°C for 1 hour and then deparaffinized in a series of xylene and ethanol washes. The sections were then treated in 80% formic acid for 10 min followed by several rinses in $dH_2O$. Antigen retrieval was performed in 0.01M citrate buffer (pH 6.0) at 80°C for 40 min. The samples were allowed to cool, rinsed in $dH_2O$ and washed 3 times in PBS. Endogenous peroxidase was quenched with 3% $H_2O_2$ in PBS for 15 min. After washing 3 times in PBS, the samples were blocked in preincubation solution (10% normal horse or goat serum, 0.25%- Triton X-100, 1% BSA in Tris-HCl pH6) for 1 hour at RT. The samples were then incubated with primary antibody (1:100 for EGT 403 and EGT-BL-LASH-N-terminal; 1:500 for BL-LSH-4B12, LB509 and pS129 ab51253; 1:2000 for BL-LASH-34-45, BL-LASH-80-96 and ab131508) diluted in preincubation solution at 4°C in a humid chamber overnight. The samples were washed 3 times in PBS then incubated with secondary antibody (Biotinylated-horse anti-mouse, Vector BA2001 1:200 or in biotinylated-goat anti-rabbit, Vector BA1000 1:200) diluted in preincubation solution for 2 hours at room temperature. After washing, samples were incubated in ABC (Vector, Vectastain Elite ABC HRP kit) solution for 1 hour at RT then washed and developed with DAB (Vector SK-4100) for 10 min. The samples were then counterstained with Meyer's Hematoxylin before dehydration and clearing followed by cover slipping with DPX. Images were taken with an Olympus BX53 microscope with a 40x objective. Three regions were imaged per section, and care was taken to image the same regions in all samples.

***In vitro calpain 1 cleavage***

Cleavage of α-syn PFFs was performed as described previously[11]. Briefly, 0.15 U of active Calpain 1 recombinant protein (Abcam, Switzerland) was added to α-syn PFFs diluted at a final concentration of 50 μM in the reaction buffer [40mM 4-(2-hydroxyethyl)-1-piperazineethanesulfonic acid (HEPES) (pH 7.5) and 5 mM dithiothreitol DTT] at 37 °C. The reaction was initiated by the addition of $CaCl_2$ (1 mM final). Samples were collected before addition of $CaCl_2$ (time 0) or 2, 5, 15 or 30 mins after the activation of Calpain 1. Cleavage of α-syn by Calpain 1 was assessed by Commassie staining after separation of the samples onto a 16.5% SDS-PAGE gels.

***Quantitative real-time RT-PCR***

WT primary neurons were treated with WT PFFs or Tris buffer (negative control) for 7, 14 and 21 days. Total RNA was isolated using RNeasy Mini Kit (Qiagen, Switzerland) according to the manufacturer's protocol. The concentration of each sample was measured with NanoDrop (NanoDrop Technologies, Wilmington, DE, USA), and the purity was confirmed using the ratios at (260/280) nm and (260/230) nm. 2 µg of RNA was used to synthesize cDNA using the High-Capacity RNA-to-cDNA Kit (Life Technologies) following *manufacturer's instructions*. To quantify α-syn, calpain 1, calpain 2 mRNAs levels, we used the SYBR green PCR master mix (Pack Power SYBR Green PCR mix, Life Technologies). Q-PCR assay was performed using the following primers synthesized by Microsynth (Balgach, Switzerland):

| Gene | Forward sequence (5'-3') | Reverse sequence (5'-3') |
|---|---|---|
| **SNCA** | AATGTTGGAGGAGCAGTGGT | GGCATGTCTTCCAGGATTCC |
| **calpain 1** | GAAAGGACCCTGGAGTGACA | TCCGGTGTAAGGTTGCAGAT |
| **calpain 2** | GTTGGTGAAAGGACATGCGT | TCAGGTTGCAGATCTCCAGG |
| **β-actin** | TTGTGATGGACTCCGGAGAC | TGATGTCACGCACGATTTCC |
| **GAPDH** | AACGACCCCTTCATTGACCT | TGGAAGATGGTGATGGGCTT |

40 cycles of amplification were then performed in an ABI Prism 7900 (Applied Biosystem, Foster City, USA) using 384-well plate which allowed simultaneously the analysis of the genes of interest and the housekeeping genes (β-actin and GAPDH) that serve as references for the normalisation step. For each independent experiment, triplicated wells were acquired per condition and each experiment was reproduced at least 3 times independently. To quantify the expression level of the genes of interest in the different conditions tested, the comparative $2^{-\Delta\Delta CT}$ method was used where ΔΔCT = ΔCT(target gene)−ΔCT(reference gene) and ΔCT = CT(target gene)−CT(reference gene). Results were expressed as the fold change relative to control neurons ($2^{-\Delta\Delta CT}$). geNorm method (RRID:SCR_006763, https://genorm.cmgg.be/) was performed to assess the most stable reference gene that should be used for the normalization of the gene expression[12].

### Supplemental information – Titles of the Figures and Legends

**Figure S1. Preparation and characterization of recombinant monomeric and PFFs α-syn species (related to Figures 1 to 8)**

**A. Purity and characterization of α-syn monomers.**

Recombinant WT mouse α-syn, human α-syn, and mouse α-syn mutants including E114A, D115A, 1-114, Δ111-115, Δ120-125, Δ133-135, Δ111-115-Δ133-135, and S129A were produced in *E. coli* and purified by anion exchange chromatography and size-exclusion chromatography, followed by a final chromatographic step using reverse-phase HPLC, as previously described[13]. The purity of recombinant monomeric α-syn after purification was assessed by ESI-LC/MS, which showed the expected mass.

**B-E. Purity and characterization of α-syn fibrils.**

α-syn fibrils were formed by incubation of monomeric α-syn for 5 days at 37°C under constant agitation at 1000 rpm. (**B**). After sonication, fibril formation was assessed by ThT fluorometry. All data represent the average ± SD (n=3). (**C**). Purity of α-syn fibrils was verified by SDS-PAGE gel and Coomassie blue staining. After sonication, fibril preparations were centrifuged, and the presence of the fibrils was verified in the pellet fraction, while the absence of monomer release after the sonication step was assessed in the supernatant fraction or after filtration through a 100 kDa filter (filtration). (**D-E**). α-syn fibrils were characterized by transmission electron microscopy (TEM) imaging. (**D**) Representative images of negatively stained α-syn fibrils before and after sonication. All α-syn fibrils showed the characteristic rigid non-branched fibrillar morphology. Scale bars = 100 nm. (**E**) Average length of the fibrils after sonication.

**Figure S2. Newly formed α-syn fibrils appear to differ in shape and subcellular localization within the same population of PFF-treated WT neurons (related to Figure 1)**

**A-C**. Morphology and subcellular localization of newly formed α-syn fibrils was assessed by ICC after 7, 14, and 21 days in PFF-treated WT neurons. After fixation, newly formed inclusions were detected using pS129 antibodies (Wako or MJFR13 or 81a). Neurons were counterstained with MAP2 antibody, and the nucleus was counterstained with DAPI staining. Confocal imaging of the newly formed inclusions located in MAP2-positive neurons. Scale bars = 10 µm.

**D-E.** Aggregates were detected using pS129 (MJFR13) in combination with total α-syn (SYN-1), p62, or ubiquitin antibodies. Neurons were counterstained with the microtubule-associated protein (MAP2) antibody, and the nucleus was counterstained with DAPI staining. Scale bars = 5 µm.

**F-H.** Addition of α-syn PFFs to primary neurons induces cell death in a time and concentration-dependent manner. Cell death levels were assessed in WT neurons treated with increasing concentrations of PFFs (70 nm to 500 nm) for up to 21 days using complementary assays. Caspase 3 activation (**F**) was measured as an early event of apoptotic cell death, while late cell death event was assessed based on the loss of plasma membrane integrity and quantified using propidium iodide as vital dye (**G**) and lactate dehydrogenase (LDH) release (**H**). As shown previously [14], increasing concentrations of α-syn PFFs in the extracellular media of the neurons enhanced cell death over time (Figure S3A-C). However, we previously showed that membrane disruption induced by adding high concentrations (2-20µM) of exogenous PFFs is a major contributor to α-syn toxicity. Therefore, to allow selective monitoring of changes in α-syn species during seeding, aggregation, and inclusion formation in neurons, we used the recommended PFF concentration of 70 nM [6,7] in all our seeding studies with this model.

**I.** Mouse cortical astrocyte cultures were treated for 24 hours with increasing concentrations of PFFs (70 to 1000 nm). After extraction, the soluble and insoluble fractions and cells lysates were analysed by immunoblotting using SYN-1 and pS129 antibodies.

**J-K.** α-syn PFF seeds are internalized via the endolysosomal pathway and truncated over time**.** KO neurons were treated with WT PFFs$^{488}$ fluorescently labelled for up to 72 hours, and the internalization and the truncation of the seeds were evaluated by confocal imaging. Neurons were counterstained with MAP2 antibody. Scale bars = 10 µm.

**J.** Internalization of the seeds via the endolysosomal pathway was confirmed by detection of the PFFs$^{488}$ fluorescently labelled seeds in LAMP1-positive (late endosome) compartments over time.

**K.** C-terminal truncation of the seeds over time was confirmed by the loss of detection of the seeds by the C-terminal α-syn antibody (134-138) (green arrows). Yellow arrows indicate intact PFF$^{488}$ fluorescently labelled seeds detected by the C-terminal α-syn antibody (134-138).

**Figure S3. List of the antibodies used in this study (related to Figures 1 to 8)**

**A**. Antibodies used for the detection of total α-syn.

For selected primary antibodies (Figure S3A) immunohistochemical staining was performed using a Menarini Intellipath automated staining machine. as previously described [15,16].

**B**. Antibodies used for the detection of α-syn phosphorylated on S129 residue.

**C**. Other antibodies used in the study.

**D**. Secondary antibodies used for immunoblotting or confocal imaging.

**Figure S4. Detection of total α-syn in studies based on the neuronal seeding model or in human PD brain tissues (related to Figure 2 and discussion)**

**A**. Table showing the antibodies used to detect total α-syn by WB approaches in studies using the seeding model in primary neuronal cultures.

**B**. Table showing the antibodies used to detect total α-syn in LB inclusions from human PD brain tissues by WB or IHC approaches.

**Figure S5. PFF cleavage and generation of α-syn truncation is a general phenomenon that occurs in human mammalian cell lines and in human induced pluripotent stem cell (iPSC)-derived dopaminergic neurons (related to Figure 2)**

**A-C.** Truncation of α-syn in HeLa cells overexpressing human α-syn.

**A.** α-syn seeding model based on addition of human PFFs to HeLa cells overexpressing human α-syn.

**B.** ICC analysis of LB-like inclusions that formed at 2 days after adding human PFFs (500 nM) to HeLa cells overexpressing human α-syn. After 48 hours of treatment, LB-like α-syn inclusions positively stained for pS129-α-syn were detected in the cytosol of HeLa cells by ICC. The panel shows examples of the aggregate heterogeneity detected using pS129 (MJF-R13 or WAKO) and total α-syn (SYN-1 or FL-140) antibodies. HeLa cells were counterstained with labelled Phalloidin[565] (actin probe) and the nucleus was counterstained with DAPI staining. Scale bars = 5 μm.

**C.** WB analysis of the truncations pattern of internalized human PFFs into the HeLa cells over 48 hours. After extractions of the soluble and insoluble fractions, cells lysates were analysed by immunoblotting. Total α-syn was detected by SYN-1. Immunoblotting with SYN-1 antibody revealed that α-syn PFFs were also rapidly processed in PFF-treated HeLa cells. Histogram showing densitometry analysis from three independent experiments. The graphs represent the mean +/- SD of 3 independent experiments. $p<0.001$=** (ANOVA followed by Tukey HSD post-hoc test, level of α-syn 15 kDa at 1 hour vs. other time-points or level of α-syn 12 kDa at 1 hour vs. other time-points).

**D-E.** Truncation of α-syn in human iPSC-derived dopaminergic neurons. **D.** Expression of neuronal midbrain markers in iPSC-derived neuronal cultures using an established methodology [3-5]. iPSC-derived neuronal cultures from a healthy control were fixed and stained for proteins expressed in human midbrain dopaminergic neurons. Tyrosine hydroxylase (TH) and FOXA2 are expressed at DIV47 of neuronal differentiation. Scale bars = 100 μm. **E.** Phosphorylation of aggregated α-syn is a rare event in iPSC-derived neurons and the levels were not measurable by WB. ICC analysis of the inclusions formed at 16 days after adding human PFFs (70 nM) to iPSC-derived neurons. The panel shows an example of a phosphorylated aggregate detected using a pS129 (ab51253) antibody and counterstained with MAP2 and DAPI. Scale bars = 50 μm.

**F.** Details of the patient's homogenates used in Figure 3E.

**Figure S6 (related to Figure 3)**

Insoluble fractions of α-syn KO primary neurons treated with 70 nM of mouse PFFs for 4 or 14 hours were separated on a 16.5% SDS-PAGE gel.

**A.** Proteomic analysis shows that the N-terminal domain of α-syn was intact in the insoluble fraction of the KO neurons treated for 14 hours with WT PFFs.

**A-D.** After Coomassie staining, two bands at ~15 (indicated by a double red asterisks) and 12 kDa (indicated by a single red asterisk) were extracted from SDS-PAGE gels (**A**, **B**). Isolated bands were selected based on the size of the proteolytic fragments observed by WB (**C**) and subjected to proteolytic digestion followed by LC-MS/MS analysis. Glu-C was used to analyse N-terminal truncation [17], which enables detection by LC-MS/MS of intact peptide corresponding to the first 13 amino acids of α-syn (**A**). To analyse C-terminal truncations, α-syn was digested using trypsin, which cannot cleave the C-terminal domain, thus allowing detection of an intact peptide corresponding to the 103-140 region by LC-MS/MS [17] (**C**). Proteomic analyses showed intact N-terminal domain of PFFs transduced into the KO neurons (**A**) but revealed that the C-terminal domain was cleaved at residues Asp-135, Ser-129, and Asp-119 with a predominant site of cleavage at residue Glu-114 (**D**). This resulted in the formation of three fragments (1-135, 1-129, and 1-119) detected in the upper band sliced in (**B-D**) and one main fragment (1-114) found in the lower band excised in (**B**). Strikingly, the cleavage sites identified in mouse α-syn were close to those detected in the LBs from human brain tissue (Asp-115, Asp-119, Asn-122, Tyr-133, and Asp-135) [18,19] or in an α-syn neuroblastoma cell line seeding model (Asp-115, Asp-119, Asn-122, Ala-124, Tyr-125, Ser-129, Tyr-133, and Tyr-135) [20]. This suggests the involvement of specific proteases in cellular responses to α-syn PFFs. Notably, human and mouse α-syn proteins differ by only 7 amino acids, of which 5 are in the C-terminal region (L100M, N103G, A107Y, G121D, S122N) and are in close proximity to the cleavage sites that we identified. This could explain the differences observed in the cleavage sites between PD models.

**E.** Epitope mapping of antibodies raised against the NAC, N-terminal, or C-terminal domains of α-syn.

**F.** N-terminal antibodies raised against the residue 1-5 or the residue 1-20 could detect full length (15 kDa, indicated by a double red asterisks) or truncated (~12 kDa indicated by a single red asterisk) α-syn in the insoluble fraction of KO neurons treated for 14 hours, confirming that the N-terminal region of α-syn PFF seeds is intact after internalization into the neurons.

**G.** Mapping of the C-terminal cleaved product using antibodies raised against the NAC and the C-terminal domains of α-syn. Immunoblots of insoluble fractions of KO neurons treated with α-syn PFFs showed that the Fragment 1-114 generated in these neurons was well recognized by the NAC antibodies [(FL-140; 61-95) and (SYN-1; 91-99)] and a C-terminal antibody raised against the residues 108-120. However, it was not recognized by antibodies raised against peptides bearing residues after 116 in the C-terminal domain [(ab6162; 116-131); (ab131508; 134-138) and (ab52168; 131-135)]. Altogether, our data demonstrate that after internalization α-syn PFF seeds are efficiently C-terminally truncated before the initiation of the intracellular seeding mechanisms.

**Figure S7. Preventing α-syn cleavage at 114 does not impact the seeding capacity of PFFs in primary neurons (related to Figure 3)**

**A.** α-syn KO neurons were treated for 14 hours with $PFFs^{WT}$, $PFFs^{1\text{-}114}$, $PFFs^{\Delta 111\text{-}115}$, $PFFs^{\Delta 133\text{-}135}$, or $PFFs^{\Delta 111\text{-}115 \Delta 133\text{-}135}$. Neurons were lysed at the indicated time, and the insoluble fractions were analysed by WB. The levels of total α-syn (SYN-1 antibody) (15 kDa or 12 kDa) were estimated by measuring the WB band intensities normalized to the relative protein levels of actin. The graphs represent the mean +/- SD of 3 independent experiments. $p<0.05$=#, (ANOVA followed by Tukey HSD post-hoc test, $PFFs^{WT}$ vs. mutants PFF-treated neurons).

**B.** α-syn fragments produced in KO neurons treated for 14 hours with $PFFs^{\Delta 111\text{-}115}$ were estimated by WB using recombinant α-syn of different sizes.

**C.** Striatum of WT mice were dissected 7 days after injection with $PFFs^{WT}$, $PFFs^{\Delta 111\text{-}115}$, or $PFFs^{\Delta 111\text{-}115 \Delta 133\text{-}135}$. The level of total α-syn (SYN-1 antibody) (15 kDa or 12 kDa) in the soluble and insoluble fractions was assessed by WB.

**Figure S8. α-syn fragments produced in neurons transduced with PFFs result from C-terminal truncation but not from N-terminal cleavage (related to 4)**

**A.** WT neurons were treated for 10 days with fluorescently labelled WT PFFs$^{488}$. Newly formed inclusions were detected using pS129 antibody (81a clone) and imaged by confocal imaging. Neurons were counterstained with MAP2 antibody, and the nucleus was counterstained with DAPI staining. Scale bars = 20 μm.

**B.** WB analyses of the insoluble fraction of WT neurons treated for 10 days with human or mouse WT PFFs. To discriminate the PFF seeds from the newly formed fibrils by WB analyses, mouse WT neurons were treated with human PFFs for 10 days, and the insoluble fraction was analysed by WB using a combination of a human-specific antibody (4B12) and a mouse-specific antibody (D37A6 XP). Although the PFFs derived from human α-syn seed less efficiently than mouse α-syn PFFs in mouse neurons[21], we still observed seeding and the formation of inclusions. As expected, the human-specific α-syn antibody (4B12) showed that the truncated fragment at ~12 kDa detected after 10 days of treatment was composed of the initial PFF seeds added to the neurons. In contrast, the newly formed aggregates (HMW bands) were mainly detected by the mouse antibody (D37A6 XP) which reflects the incorporation and aggregation of the endogenous mouse α-syn protein. Additionally, it was interesting to note that endogenous mouse α-syn was cleaved upon human PFF-seeding condition.

**C.** Proteomic analysis shows that the N-terminal domain of α-syn was intact in the insoluble fraction of the WT neurons treated for 7 and 21 days with WT PFFs.

**D-E.** α-syn WT neurons were treated for 7 days with PFFs$^{WT}$. Newly formed inclusions were detected using pS129 antibody (81a clone) in combination with N-terminal (**D**, epitope 1-20) or NAC (**E**, epitope 80-92) antibodies together with C-terminal epitopes [116-131] (**D**) or [134-138] (**E**) antibodies. The newly formed aggregates were similarly detected by the pS129, N-ter, NAC, or C-terminal antibodies. Scale bars = 20 μm.

**Figure S9. Antibodies targeting different epitopes reveal different forms of α-syn pathology in PD and MSA patients (related to Figure 4 J-L)**

**A.** Consecutive sections (5 µm thick) of paraffin-embedded substantia nigra from PD patients were stained with antibodies targeting N-terminal, NAC-region and C-terminal α-syn epitopes (Figure S3). Three regions were imaged and compared for each of the staining. NAC-region antibodies revealed the heaviest load of Lewy bodies and Lewy neurites. C-terminal antibodies revealed nuclear structures not seen with other antibodies. N-terminal antibodies detect mild pathological α-syn. Scale bars = 20 µm.

**B-E.** Paraffin-embedded serial sections from the substantia nigra (**B**), *SNCA* G51D pons (**C**) or cingulate gyrus (**D**) of PD patients or pons from MSA patients (**E**) were stained with the α-syn N-terminal antibodies LASH-EGT-1-20 (epitope: 1-20) and LASH-BL-A15110B (epitope: 34-45), the C-terminal antibody LASH-EGT408 (epitope: 121-132) and the pS129 antibody (Biolegend, clone 81a) (See also, Figure S3). Primary antibodies were omitted for the negative control tissue staining. Scale bars = 20 µm.

Our work underscores the critical importance of using multiple antibodies to reveal the pathological heterogeneity of α-syn inclusions in human post-mortem tissue, animal models of PD and other synucleinopathies.

**Figure S10. C-terminal truncations alter newly formed α-syn fibrils and consequently disrupt interactions with the C-terminal interacting partners**

**A**. Strategy to add biotin to monomeric α-syn.

**B**. ESI-LC/MS analysis of biotinylated M1C α-syn (observed mass: 14958 Da, expected mass: 14958 Da).

**C**. Water acuity UPLC spectra of biotinylated M1C α-syn using a BEH 300 C4 column 2.1 mm × 100 mm, 1.7 µm with a linear gradient of 10−90% B over 4 min (solvent A: water/0.1% TFA, solvent B: acetonitrile/0.1% TFA).

**D**. SDS-PAGE analysis of biotinylated M1C α-syn.

**E**. WB of biotinylated M1C α-syn fibrils with and without boiling in sample buffer.

**F**. TEM of biotinylated M1C α-syn fibrils.

**G.** PFFs labelled with biotin were incubated with mouse cortical brain lysates, and biotinylated fibrils were specifically pulled-down using streptavidin beads.

**H.** Primary neurons were treated for 14 days or 21 days with PFFs$^{WT}$, and the insoluble fraction was isolated. The proteome of the insoluble fractions from the pull-down fraction (**G**) and the PFF-treated neurons (**H**) were identified using LC-MS/MS analysis. Identified α-syn interactors were compared to the previously reported putative C-terminal interactors of α-syn[22]. **I.** Table depicts the presence of the putative C-terminal interactors of α-syn[22-27] in the pull-down assay using full-length biotinylated PFFs$^{WT}$ (**G**) or in the insoluble fractions of PFF$^{WT}$-treated neurons over time (**H**).

**Figure S11. Temporal profiling of the LB-like inclusions formation by correlative light electron microscopy (related to Figure 6)**

**A.** PFFs were added for 7, 14, and 21 days to the extracellular media of hippocampal neurons plated on dishes with alphanumerical searching grids imprinted on the bottom, allowing an easy localization of the cells. Neurons were fixed at the indicated time and imaged by confocal microscopy (**B-D, F-I**, top images, scale bars = 10 µm). The selected neurons were embedded and cut by an ultramicrotome. Serial sections were examined by TEM.

**B-D**. Representative confocal images (top images) and EM micrographs (bottom images) of the control neurons treated with Tris buffer for 7 days (**B**) or 21 days (**C**) or neurons treated with PFFs for 14 days (**D, F-H**) or 21 days (**G-I**). Microtubule is highlighted in orange (**B**), and nucleus is highlighted in blue. Autophagolysosomal-like vesicles are indicated by a yellow asterisk, and mitochondrial compartments are indicated by a green asterisk. **D.** CLEM imaging confirmed that PFFs did not accumulate on the outer side of the plasma membrane when used at a nanomolar concentration, as previously shown when higher concentrations were applied to the neurons[14].

**E.** Graph representing the mean +/- SD of the width of the microtubules compared to the newly formed fibrils at day 7 (a minimum of 120 microtubules or newly formed fibrils were counted). Measurements confirmed that the width of newly formed α-syn fibrils of 11.94±3.97 nm (SD) is significantly smaller than the average width of the microtubules of 18.67±2.94 nm (SD). $p<0.001$=*** (student t-test for unpaired data with equal variance), indicating that this parameter can be used to discriminate the newly formed fibrils from the cytoskeletal proteins.

**J-K.** The length of newly formed fibrils was measured over time. A minimum of 160 fibrils were counted for each condition.

**L.** Enlarged image of α-syn inclusion at Day 21. Inclusions is highlighted in red and newly formed α-syn fibrils in black. Autophagolysosomal-like vesicles are highlighted in yellow, lysosomes are highlighted in purple, membranes are highlighted in pink, mitochondrial compartments are highlighted in green and nucleus is highlighted in blue.

**B-C.** Scale bars = 500 nm. (**D**) at low magnification**,** scale bar = 5 µm, and (**F-H**) at higher magnifications: (**F**) scale bar = 500 nm; (**H**) scale bar = 300 nm. **G.** Scale bar = 1 µm. **I.** Scale bar = 300 nm. **L.** Scale bar = 1 µm.

**Figure S12. Calpains 1 and 2 are involved in the truncation of α-syn *in vitro* and primary neurons (related to Figure 7)**

**A-B.** Immunoblotting analyses showing the protein levels over time of calpain 1 (**A,** top panel), calpain 2 (**A**, bottom panel), and α-syn (**B**) in the extracellular media of PFF-treated WT neurons or in control neurons (Tris).

**C.** Temporal proteomic analyses showing the enrichment of calpain 2 protein in the insoluble fraction of PFF-treated WT neurons.

**D-F.** Transcriptional regulation of α-syn (**D**), calpain 1 (**E**) and calpain 2 (**F**) in WT neurons treated with PFFs for up to 21 days. Quantitative RT-qPCR was performed with primers specific for α-syn (**D**), calpain 1 (**E**) and calpain 2 (**F**) at the indicated time-points after adding PFFs to WT neurons. Results are presented as fold increases in comparison to the respective level in control neurons treated with Tris buffer. mRNA levels were normalized to the relative transcriptional levels of GAPDH and actin housekeeping genes. The graphs represent the mean +/- SD of three independent experiments. $p<0.05$=* (ANOVA followed by Tukey HSD post-hoc test, Tris vs. PFF-treated neurons). Interestingly, α-syn mRNA levels increased significantly at day 3 when the intact α-syn protein level was lowest, suggesting that α-syn gene expression was enhanced to compensate for α-syn protein depletion from the cytosolic fraction during the aggregation in PFF-treated neurons (**D**). Conversely, PFF-treated neurons showed a significant reduction of α-syn mRNA levels at 21 days after treatment (**D**). This could possibly reflect cellular responses aimed at downregulating α-syn protein levels to prevent further aggregation and inclusion formation. Additionally, calpain 1 mRNA level was markedly upregulated at day 21 (**E**), which was concomitant with the decrease in α-syn mRNA level (**D**). However, no significant increase was observed for calpain 2 mRNA levels (**F**).

**G.** α-syn KO neurons were pre-treated for 6 hours with increasing concentrations of calpain 2 inhibitor (10, 20, or 40 µM), calpain 1 inhibitor PD105606 (10, 20, or 40 µM), or DMSO as a control. 70 nM of PFFs were then added for 14 hours. After sequential extractions of the soluble and insoluble fractions, cell lysates were analysed by immunoblotting. Total α-syn was detected by SYN-1 antibody. The histogram shows the densitometry analysis from 3 independent experiments. The graphs represent the mean +/- SD of 3 independent experiments. $p<0.0001$=*** (ANOVA followed by Tukey HSD post-hoc test, DMSO vs. enzymatic inhibitors in KO neurons treated with PFFs).

**Figure S13. Truncation of α-syn is required for inclusion formation and maturation in cells (related to Figure 8)**

Level and morphology of the pS129-positive inclusions formed in PFF-treated WT neurons in presence of Calpain inhibitor 1 were assessed after immunostaining using pS129 (81a), total α-syn (epitope: 134-138, top panel), or total α-syn (epitope: 116-131, bottom panel) and MAP2 (neurons) antibodies. Nuclei were counterstained with DAPI staining. Scale bars = 5 µm.

**Figure S14. Blocking C-terminal truncation: implications for understanding the diversity of α-synuclein pathologies and spreading**

Our study has demonstrated that the C-terminal cleavage of α-syn is one of the primary triggers for initiating α-syn aggregation *in vivo* and packing the fibrils into higher-order inclusions. Consistent with our working hypothesis, blocking or reducing α-syn C-terminal cleavage at the monomer level (**A**) or at the fibrils level (**B**) using monoclonal antibodies directed against the C-terminal part of α-syn or using inhibitors of proteases that cleave within the C-terminus of α-syn could protect against α-syn-induced neurodegeneration and could attenuate α-syn pathology spreading in human and animal models of PD and related synucleinopathies. Together, these findings suggest that inhibiting C-terminal represents a viable therapeutic strategy for the treatment of PD and synucleinopathies.

**Figure S1. Related to Figures 1-8**

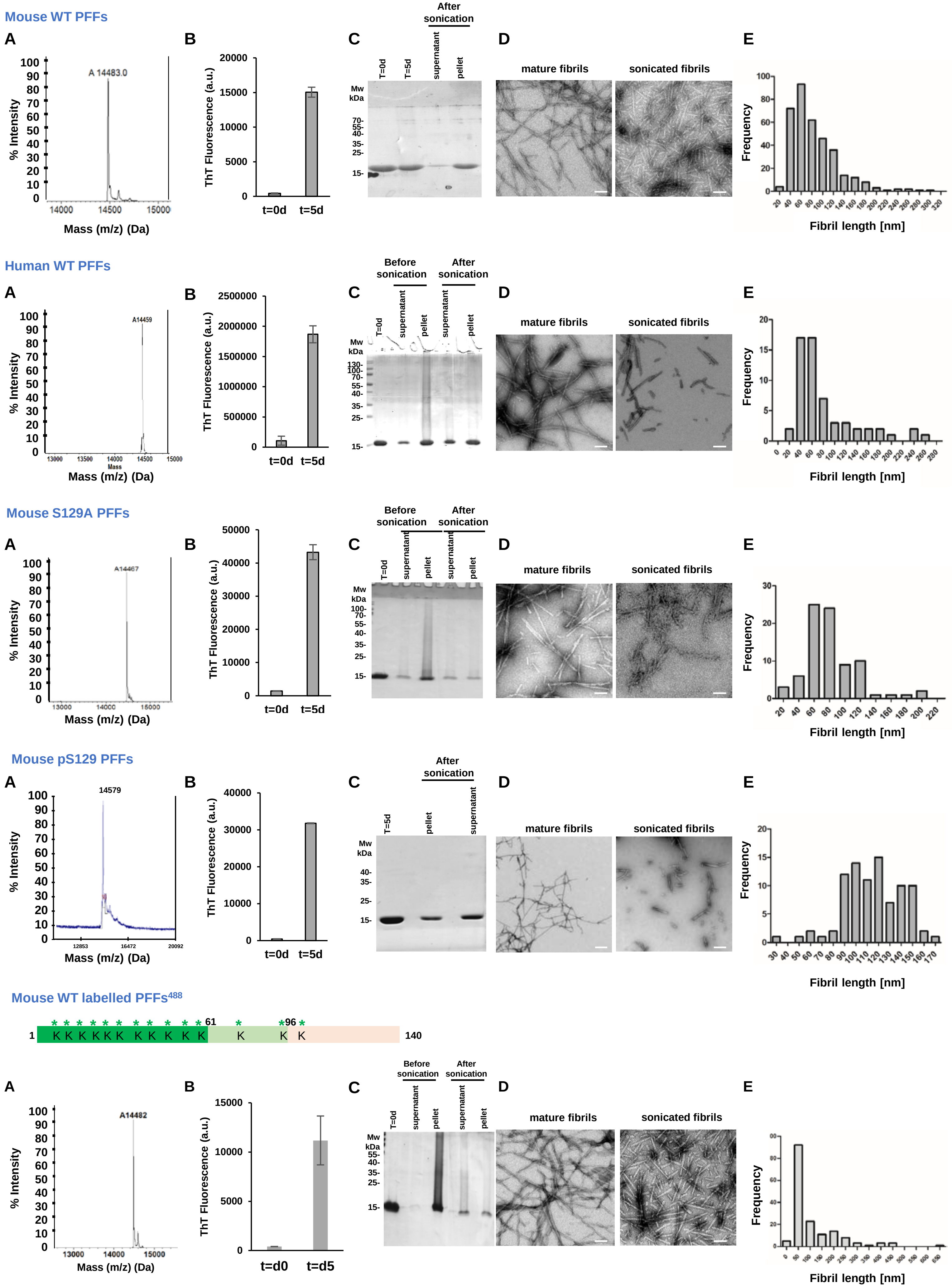

**Figure S1. Related to Figures 1-8**

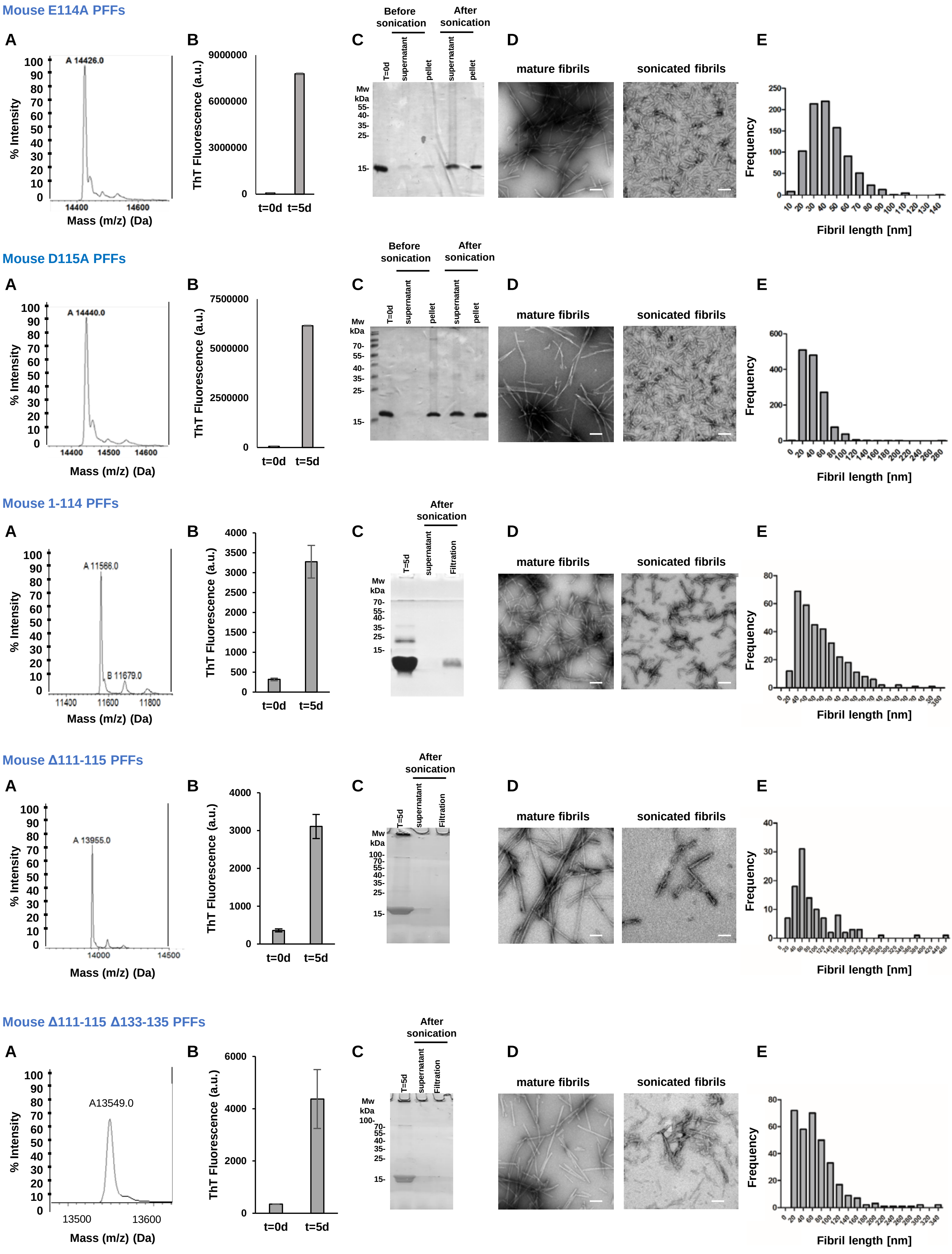

**Figure S1. Related to Figures 1-8**

### Mouse Δ120-125 PFFs

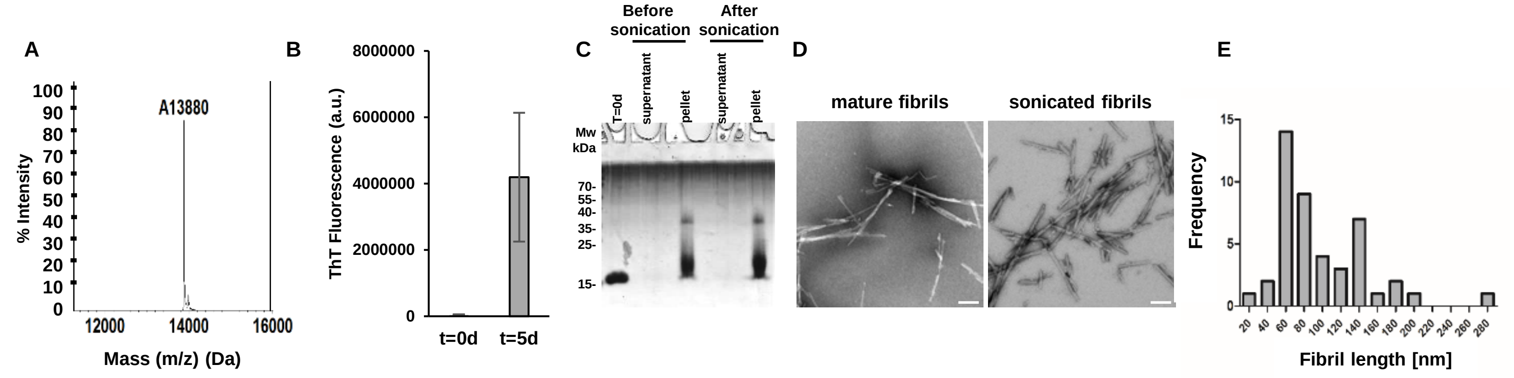


### Mouse Δ133-135 PFFs

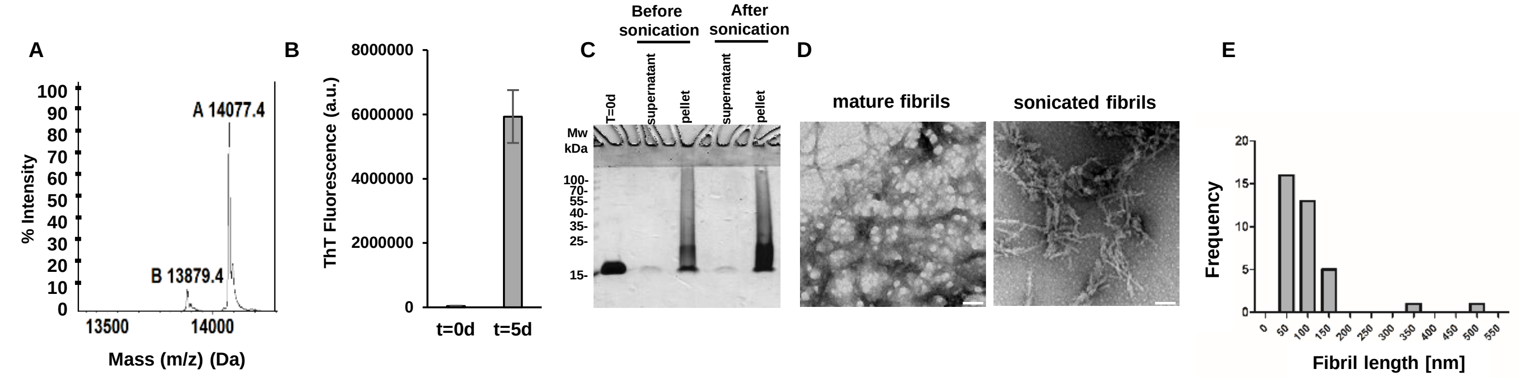

**Figure S2. Related to Figure 1**

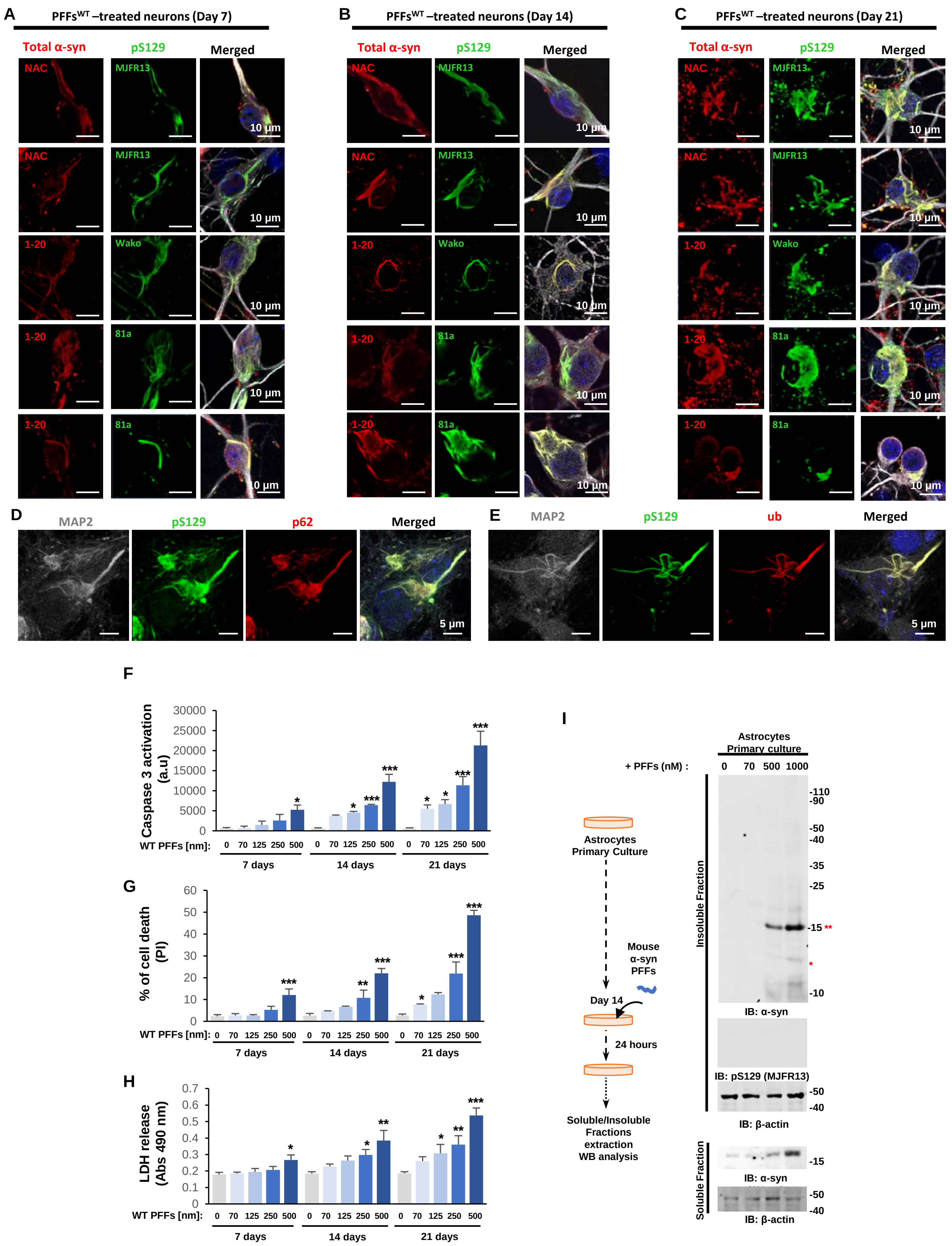

Figure S2. Related to Figure 1

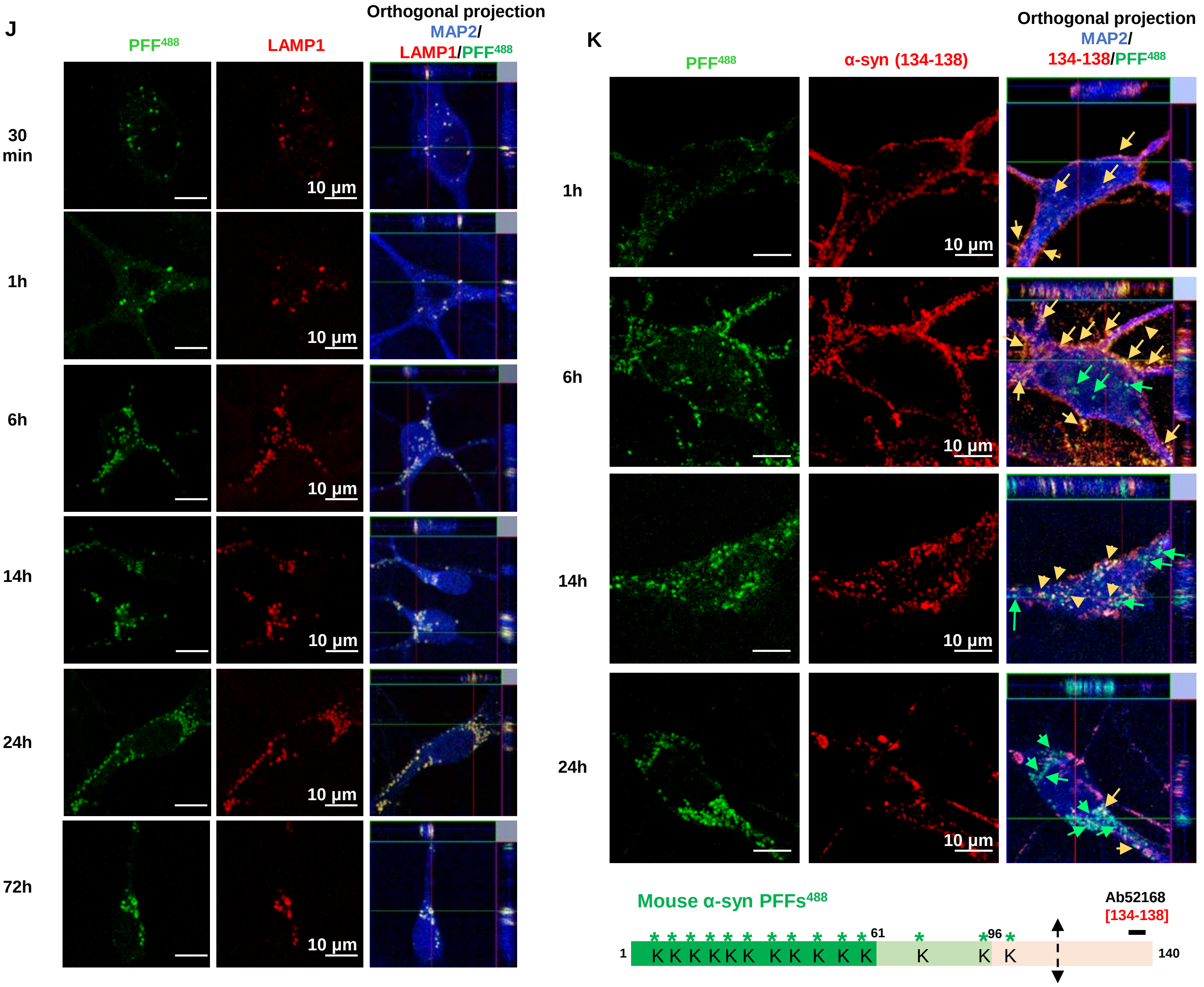

**Figure S3. Related to Figures 1 to 8**

**A**

| Primary Antibody | Catalog # | Company | Clone | RRID | Host | Concentration | WB dilution | ICC dilution | IHC dilution | Epitope |
|---|---|---|---|---|---|---|---|---|---|---|
| anti-α-syn **total** | LASHUEL | - | LASH-EGT403 | - | Mouse | 1.12 mg/ml | 1:500 | 1:250 | | **1-5** |
| anti-α-syn **total** | LASHUEL | - | LASH-EGT1-20 | - | Rabbit | Not provided | 1:1000 | 1:1000 | 1:6'000 (Menarini auto-staining) | **1-20** |
| anti-α-syn **total** | LASHUEL | - | LASH-BL-A15110B | - | Mouse | | 1:1000 | 1:500 | 1:10'000 (Hand staining) | **34-45** |
| anti-α-syn **total** | Sc-10717 | Santa-Cruz | Fl-140 | RRID:AB_2302268 | Rabbit | 0.2 mg/ml | 1:1000 | 1:500 | | **61-90** |
| anti-α-syn **total** | LASHUEL | - | LASH-BL-A15115A | - | Mouse | 0.8 mg/ml | 1:1000 | 1:500 | | **80-96** |
| anti-α-syn **total** | 610787 | BD | SYN-1 | RRID:AB_398108 | Mouse | 0.25 mg/ml | 1:1000 | 1:1000 | | **91-99** |
| anti-α-syn **total** | 807801 | Biolegend | 4B12 | RRID:AB_2564730 | Mouse | 1mg/ml | 1:1000 | - | | **103-108** |
| anti-α-syn **total** | AB5336P | Millipore | - | RRID:AB_2192954 | Sheep | 1 mg/ml | 1:500 | 1:500 | | **108-120** |
| anti-α-syn **total** | ab6162 | Abcam | - | RRID:AB_2192805 | Sheep | 1 mg/ml | 1:500 | 1:500 | | **116-131** |
| anti-α-syn **total** | LASHUEL | - | LASH-EGT408 | - | Mouse | 1.184 mg/ml | - | 1:500 | 1:500 (Menarini auto-staining) | **121-132** |
| anti-α-syn **total** | ab52168 | Abcam | - | RRID:AB_869970 | Rabbit | 1 mg/ml | 1:500 | 1:500 | | **131-135** |
| anti-α-syn **total** | ab131508 | Abcam | - | RRID:AB_11155736 | Rabbit | 1 mg/ml | 1:500 | 1:200 | | **134-138** |
| anti-α-syn **total** | 4179 | Cell signalling | D37A6 | RRID:AB_1904156 | Rabbit | Not provided | 1:1000 | - | | **near Glu105 residue** |

**B**

| Primary Antibody | Catalog # | Company | Clone | RRID | Host | Concentration | WB dilution | ICC dilution | IHC dilution | Epitope |
|---|---|---|---|---|---|---|---|---|---|---|
| anti-**pS129**-α-syn | Ab168381 | Abcam | MJF-R13 | RRID:AB_2728613 | Rabbit | 4.229 mg/ml | 1:500 | 1:500 | | Not provided |
| anti-**pS129**-α-syn | 825701 | BioLegend | P-syn/81A | RRID:AB_2564891 | Mouse | 1.0 mg/ml | 1:1000 | 1:500 | 1:500 (Menarini auto-staining) | AYEMPpSEEGYQ |
| anti-**pS129**-α-syn | 015-25191 | Wako | pSyn#64 | RRID:AB_2537218 | Mouse | 3.0 mg/ml | 1:500 | 1:500 | | AYEMPpSEEGYQ |
| anti-**pS129**-α-syn | 010-26481 | Wako | pSyn#64, biotin conjugated | - | Mouse | 3.0 mg/ml | 1:500 | 1:500 | | AYEMPpSEEGYQ |
| anti-**pS129**-α-syn | Ab51253 | Abcam | EP1536Y | RRID:AB_869973 | Rabbit | 1 mg/ml | 1:500 | 1:500 | | |

**C**

| Primary Antibody | Catalog # | Company | Clone | RRID | Host | Concentration | WB dilution | ICC dilution | Epitope |
|---|---|---|---|---|---|---|---|---|---|
| anti-**actin** | ab6276 | Abcam | AC-15 | RRID:AB_2223210 | Mouse | 2.2 mg/ml | 1:5000 | Not tested | DDDIAALVIDNGSGK |
| anti-**MAP2** | ab92434 | Abcam | - | RRID:AB_2138147 | Chicken | Not provided | Not tested | 1:2000 (ICC) | Recombinant full length protein |
| anti-**p62** | H00008878 | Abnova | 2C11 | RRID:AB_437085 | Mouse | 1 mg/ml | 1:1000 | 1:500 | raised against a full length recombinant SQSTM1 |
| anti-**ubiquitin** | Sc-8017 | Santa-Cruz | P4D1 | RRID:AB_628423 | Mouse | 0.2 mg/ml | 1:500 | 1:500 | 1-76 |
| anti-**calpain 1** | LS-B4768 | LsBio | - | RRID:AB_10801434 | Mouse | 1 mg/ml | 1:500 | 1:500 | See data sheet |
| anti-**calpain 2** | ab39165 | Abcam | - | RRID:AB_725844 | Rabbit | 1 mg/ml | 1:500 | 1:500 | See data sheet |
| anti-**TH** | AB152 | Millipore | - | RRID:AB_390204 | Rabbit | 0.1 mg/ml | - | 1:500 | See data sheet |
| anti-**FOXA2** | AF2400 | R&D | - | RRID:AB_2294104 | Goat | 0.2 mg/ml | - | 1:250 | See data sheet |
| anti-**Actin-HRP** | ab49900 | Abcam | AC-15 | RRID:AB_867494 | - | 3.8 mg/ml | 1:50000 | - | DDDIAALVIDNGSGK |

**D**

| Secondary Antibody | Catalog # | Company | RRID | Concentration | WB dilution | ICC dilution |
|---|---|---|---|---|---|---|
| Goat anti-**mouse AF680** | A21058 | Invitrogen | RRID:AB_2535724 | 2 mg/ml | 1:5000 | - |
| Goat anti-**rabbit AF680** | A21109 | Invitrogen | RRID:AB_2535758 | 2 mg/ml | 1:5000 | - |
| Goat anti-**mouse AF800** | 926-32210 | Li-Cor | RRID:AB_621842 | 1 mg/ml | 1:5000 | - |
| Goat anti-**rabbit AF800** | 926-32211 | Li-Cor | RRID:AB_621843 | 1 mg/ml | 1:5000 | - |
| Goat anti-**mouse AF647 + nanogold particles** | 7502 | Nanoprobes | - | | - | 1:800 |
| Donkey anti-**mouse AF568** | A10037 | Invitrogen | RRID:AB_2534013 | 2 mg/ml | - | 1:800 |
| Donkey anti-**rabbit AF568** | A10042 | Invitrogen | RRID:AB_2534017 | 2 mg/ml | - | 1:800 |
| Donkey anti-**rabbit AF647** | A31573 | Invitrogen | RRID:AB_2536183 | 2 mg/ml | - | 1:800 |
| Donkey anti-**mouse AF647** | A31571 | Invitrogen | RRID:AB_162542 | 2 mg/ml | - | 1:800 |
| Donkey anti-**chicken AF405** | 703-475-155 | Jackson Immunoresearch | RRID:AB_2340373 | 1 mg/ml | - | 1:400 |
| Donkey anti-**chicken AF488** | 703-545-155 | Jackson Immunoresearch | RRID:AB_2340375 | 1 mg/ml | - | 1:500 |
| **Phalloidin Atto594** | 94072 | Sigma | - | 10nM | - | 1:100 |

**Figure S4. Related to Figure 2 and discussion section**

## A

| Previous studies – Seeding model | Primary Antibody | Clone | Epitope | Applications | Detection of C-ter fragments |
|---|---|---|---|---|---|
| **Liu et al., 2005** | anti-α-syn **total** | **SYN-1** | **91-99** | WB | **Yes** |
| **Luk et al., 2009** | anti-α-syn **total** | SNL-4 | 2-12 | WB | - |
| **Volpicelli-Daley et al., 2011** | anti-α-syn **total** | Mouse-specific | 115-125 | WB | - |
| | | SNL-1 | 104-119 | WB | - |
| **Luk et al., 2012** | anti-α-syn **total** | SNL-1 | 104-119 | WB | NO |
| | | SNL-4 | 2-12 | | |
| | | LB509 | 115-122 | WB | NO |
| | | SYN211 | 120-125 | WB | NO |
| | | SYN514 | Nter-oxidized α-syn | | |
| **Tanik et al., 2013** | anti-α-syn **total** | SYN211 | 120-125 | WB | NO |
| | | Mouse α-syn | | | |
| **Tran et al., 2014** | anti-α-syn **total** | SYN202 | 130-140 | WB | NO |
| **Henderson et al., 2017** | anti-α-syn **total** | SNL-4 | 2-12 | | |
| **Karampetsou et al., 2017** | anti-α-syn **total** | C20 | C-ter region | WB | WB did not allow detection of the bands < 15 kda |
| | | **4B12** | **103-108** | WB | **Yes** |
| | | SYN-1 | 91-99 | WB | WB did not allow detection of the bands < 15 kda |
| **Tapias et al., 2017** | anti-α-syn **total** | **SYN-1** | **91-99** | WB | **Yes** |
| **Lee et al., 2018** | anti-α-syn **total** | **SYN-1** | **91-99** | WB | **Yes** |
| **Grassi et al., 2018** | anti-α-syn **total** | **EP1646Y** | **1-100** | WB | **Yes** |
| | | **D37A6** | **Glu105** | WB | **Yes** |
| | | **PAS-17239** | **C-ter region** | WB | **Yes** |

## B

| Detection of α-syn inclusions in human brain tissues | Primary Antibody | Clone | Epitope | Applications | Detection of C-ter fragments |
|---|---|---|---|---|---|
| **Gai et al., 1998** | anti-α-syn **total** | AB5038 | 111-131 | IHC | Not applicable |
| **Braak et al., 2003** | anti-α-syn **total** | | 116-131 | IHC | Not applicable |
| **Anderson et al., 2006** | anti-α-syn **total** | **SYN-1** | **91-99** | WB | **Yes** |
| **Tsuboi et al., 2007** | anti-α-syn **total** | | 116-131 | IHC | Not applicable |
| **Alafuzoff et al., 2009** | anti-α-syn **total** | Nova-Castra | Full length | IHC | Not applicable |
| **Dickson et al., 2010** | anti-α-syn **total** | | 116-131 | IHC | Not applicable |
| **Schneider et al., 2012** | anti-α-syn **total** | Zymed | 117-124 | IHC | Not applicable |
| **Henstridge et al., 2012** | anti-α-syn **total** | SYN211 | 120-125 | IHC | Not applicable |

### Figure S5. Related to Figure 2

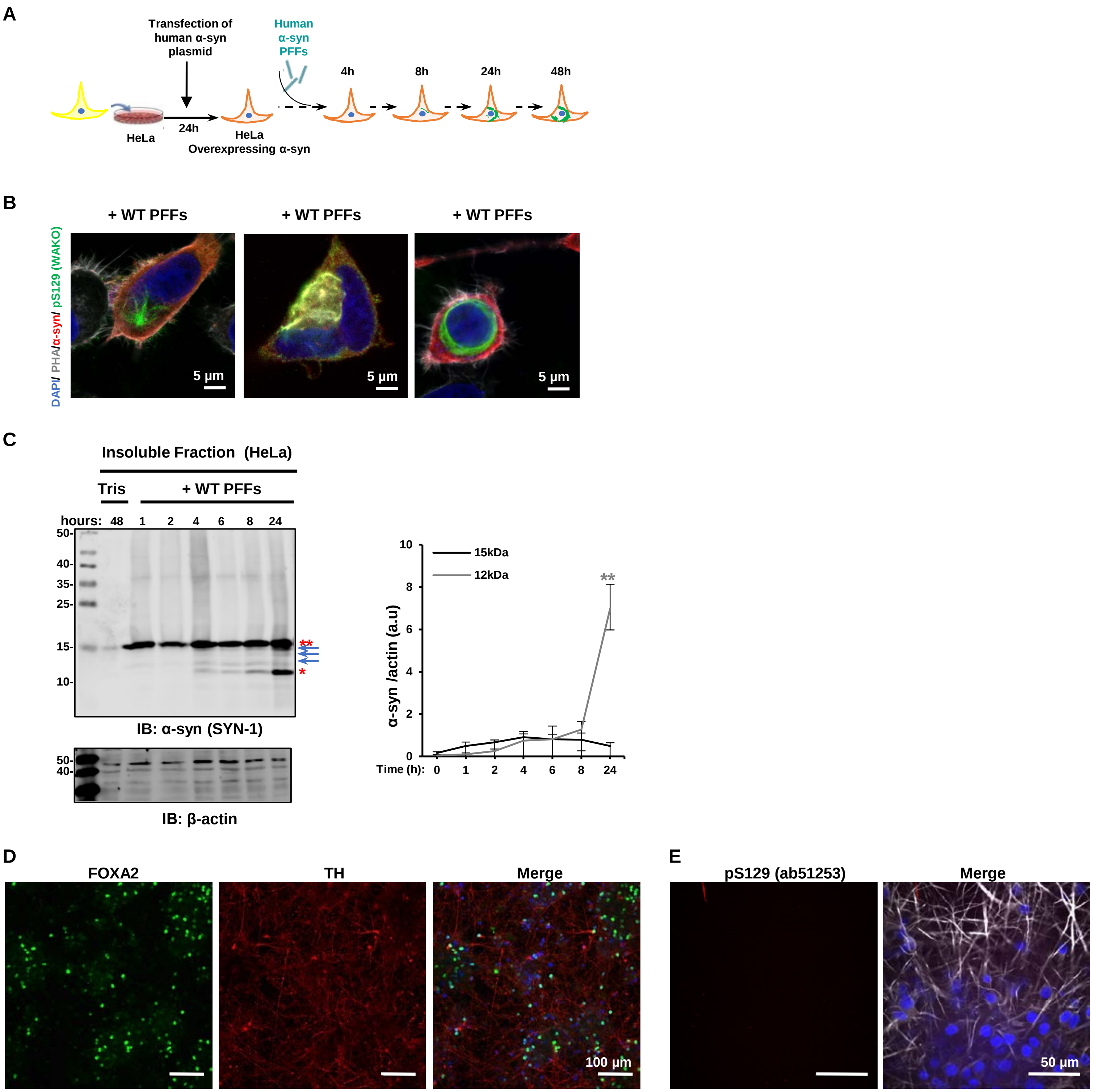


**F**

**Details of the Patient's homogenates used in Figure 3E**

| Number | NBB sample ID | Autopsy ID | Sex | Age | Diagnosis | region |
|---|---|---|---|---|---|---|
| 1 | 2012-010 | S12/010 | F | 59 | Muliple system atrophy | Cerebellum/dentatus |
| 2 | 2012-039 | S12/039 | F | 70 | Muliple system atrophy | Cerebellum/dentatus |
| 3 | 2012-047 | S12/047 | F | 61 | Muliple system atrophy | Cerebellum/dentatus |
| 4 | 2011-081 | S11/081 | M | 55 | Non-demented control | Medial temporalis gyrus |
| 5 | 2011-082 | S11/082 | F | 84 | Non-demented control | Medial temporalis gyrus |
| 6 | 2014-043 | S14/043 | F | 60 | Non-demented control | Locus coeruleus + pons |

**Figure S6. related to Figure 3**

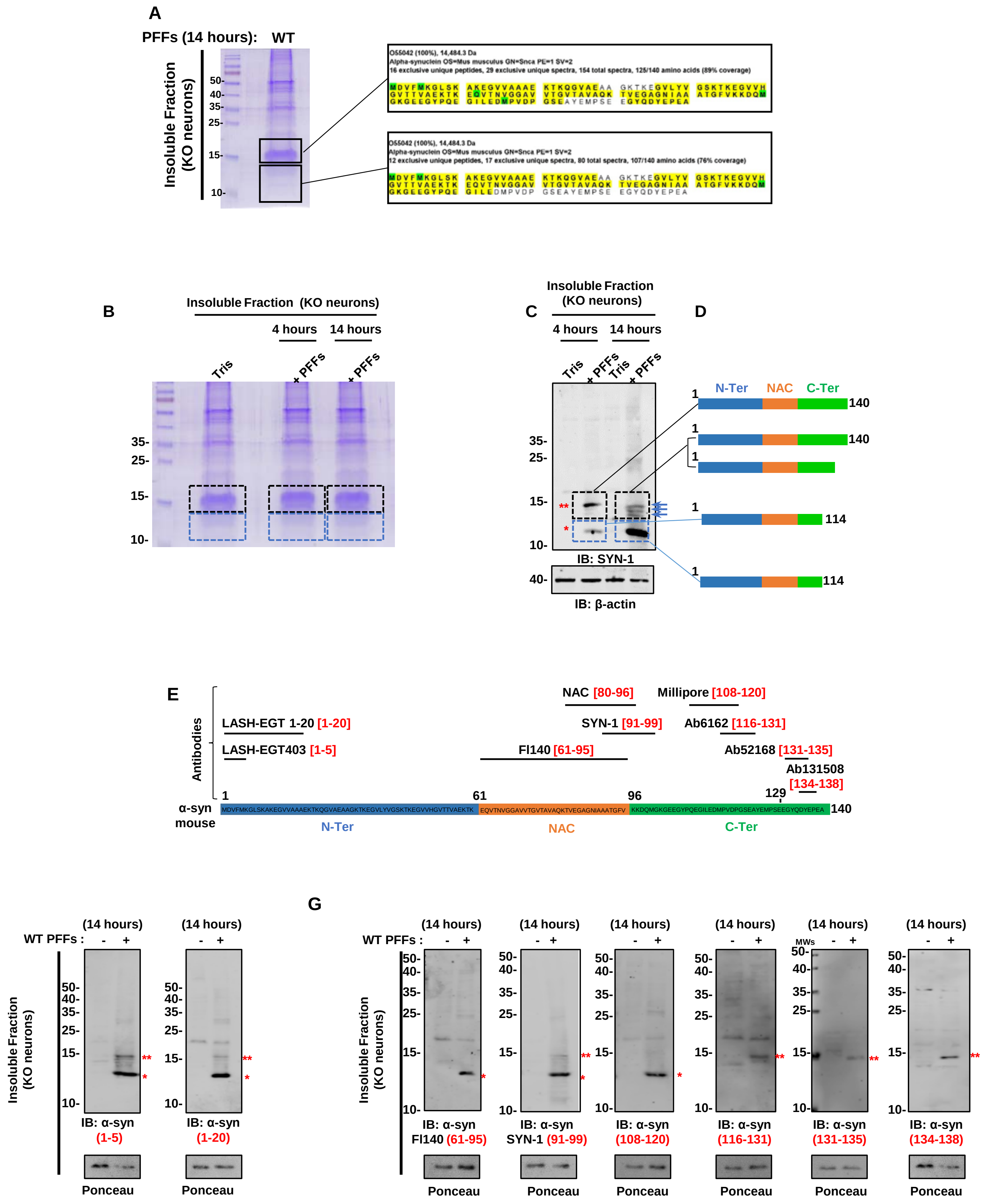

Figure S7. Related to Figure 3

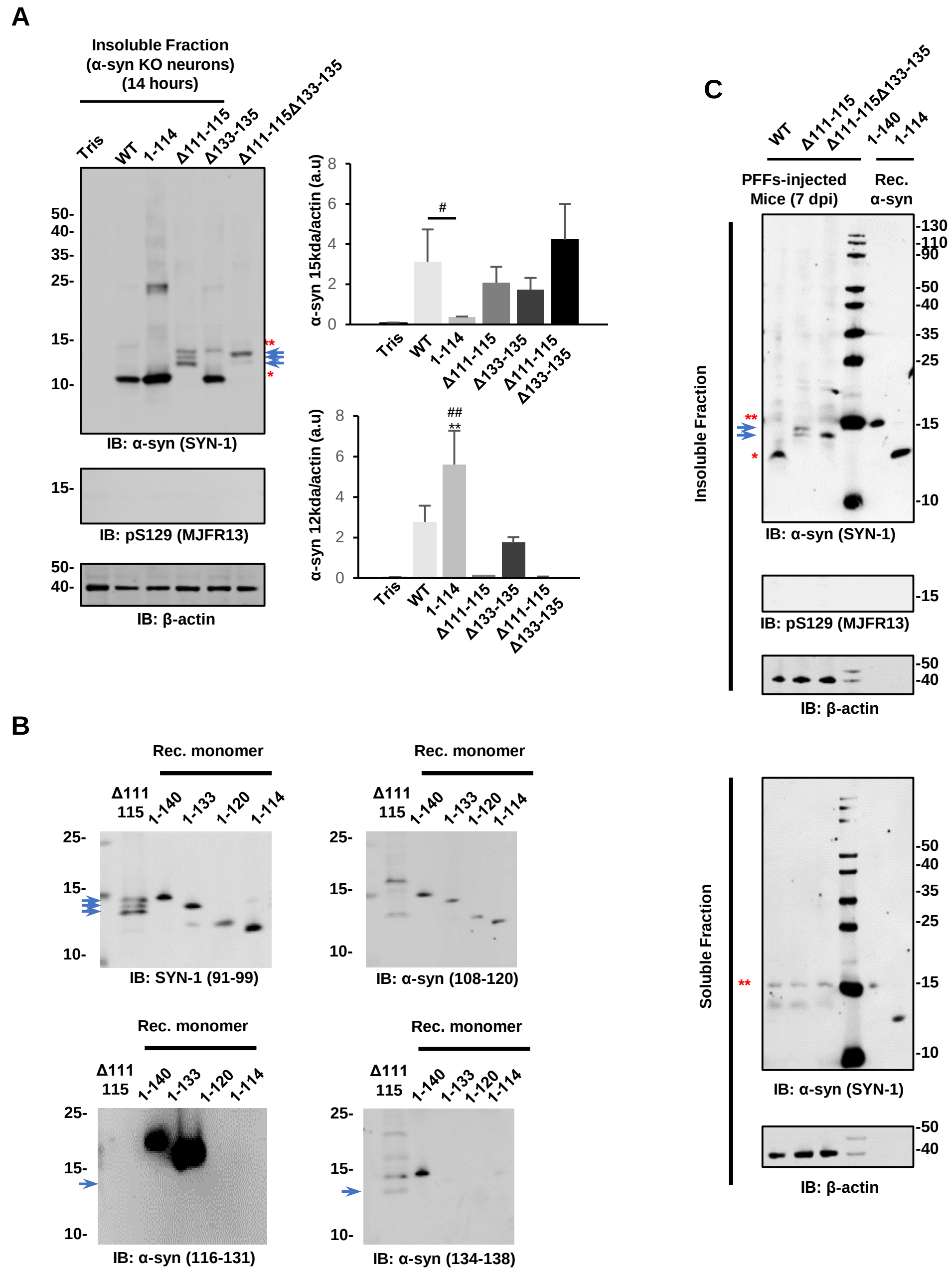

**Figure S8. Related to Figure 4**

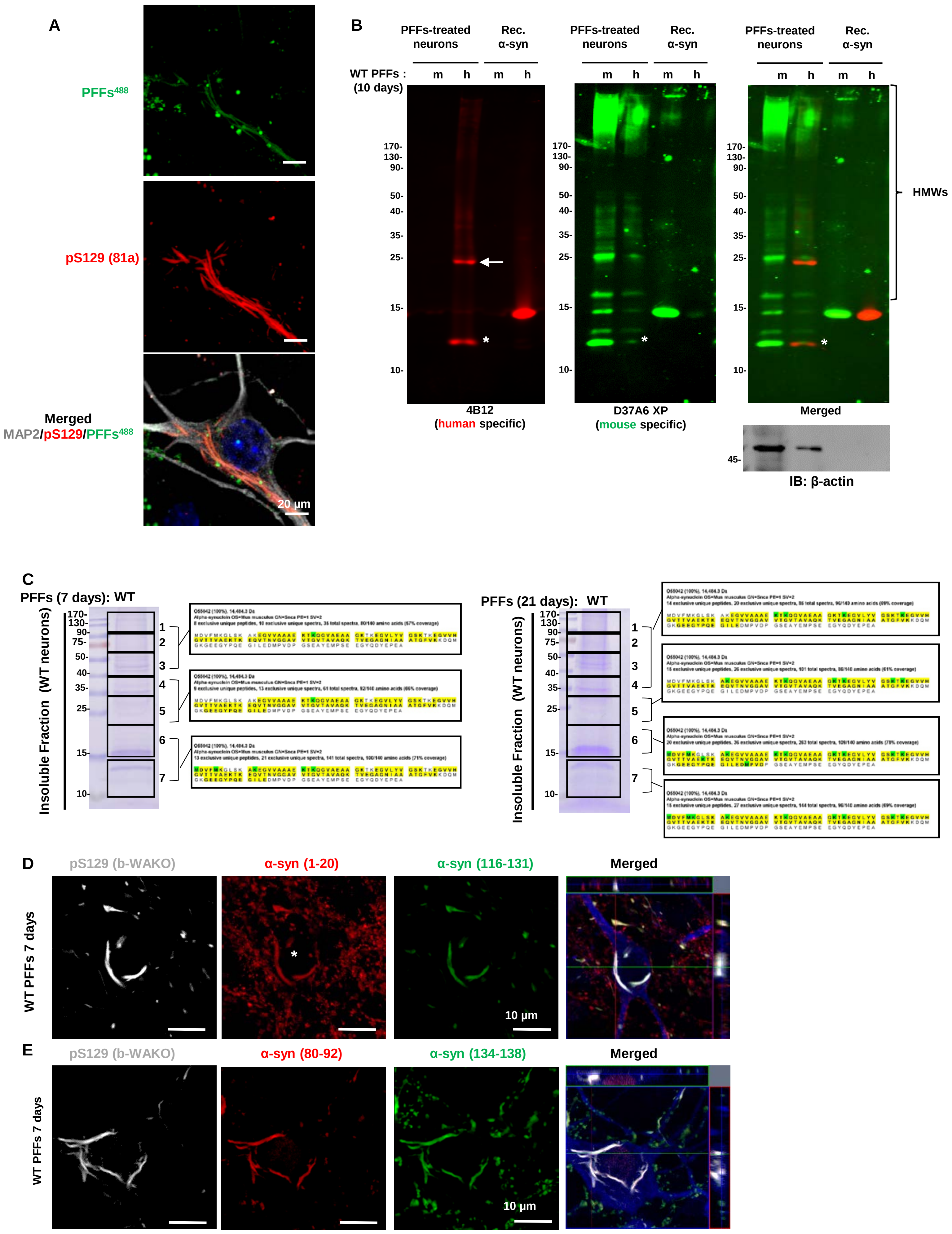

Figure S9. Related to Figure 4 J-L

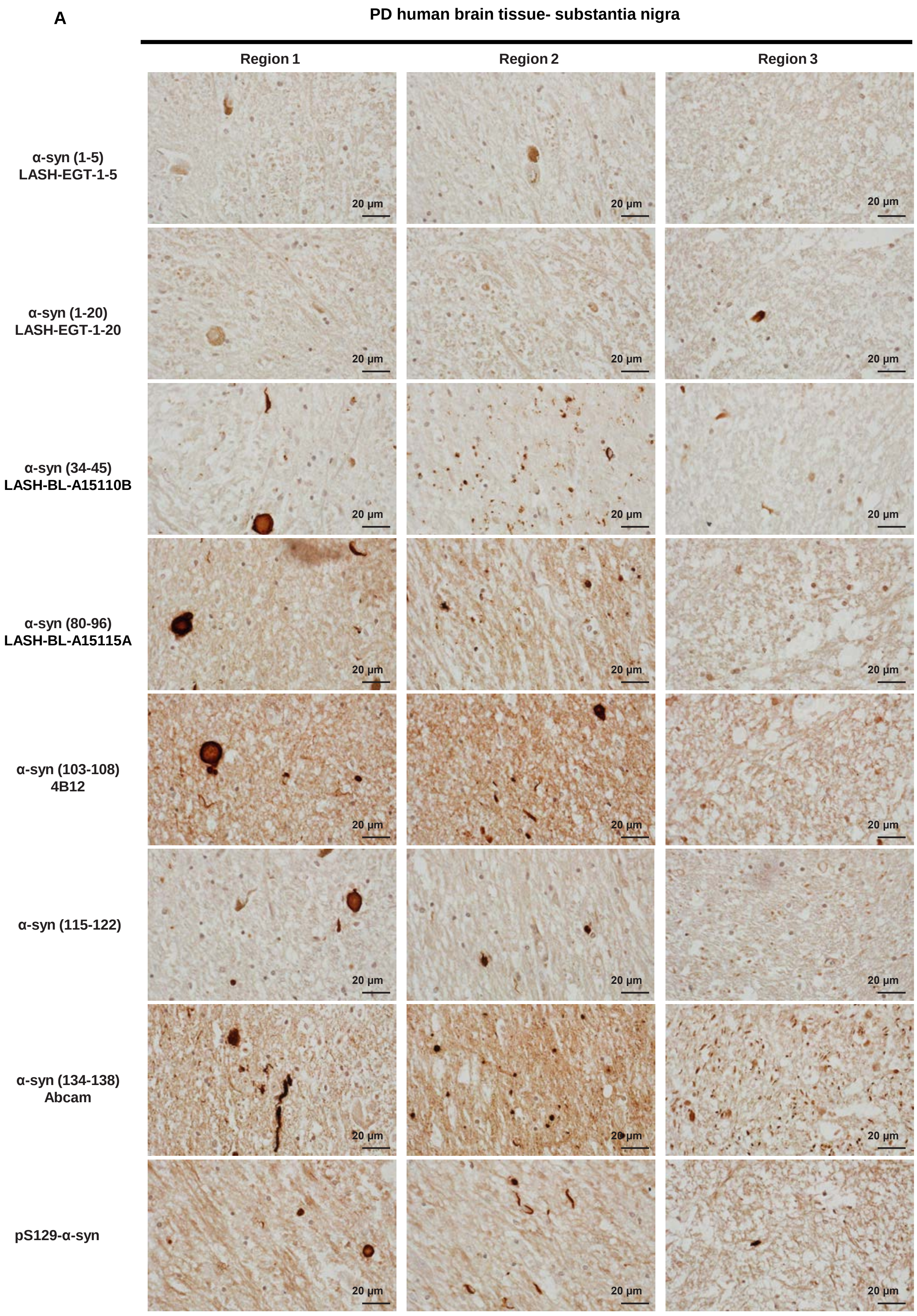

**Figure S9. Related to Figure 4 J-L**

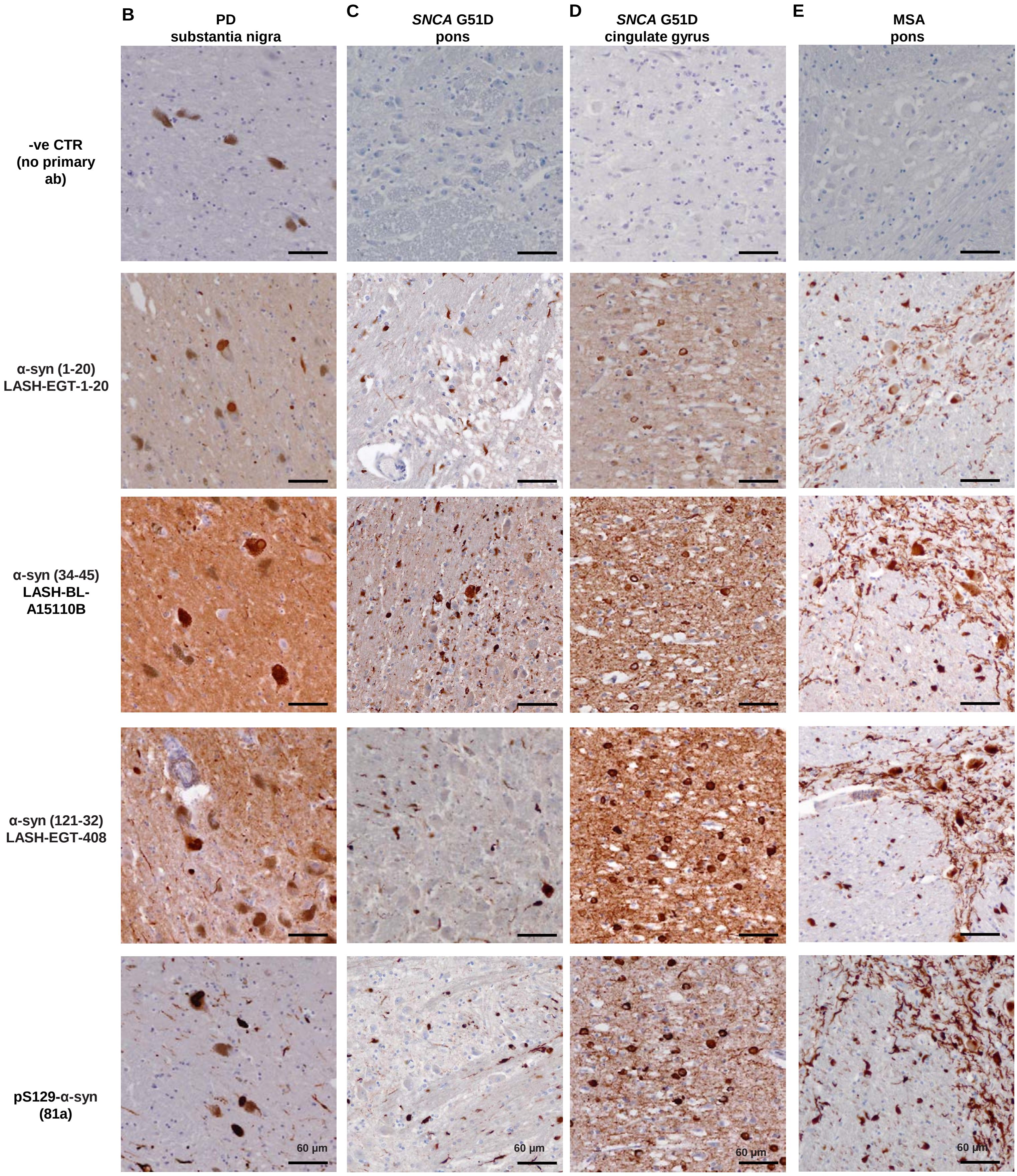

**Figure S10.**

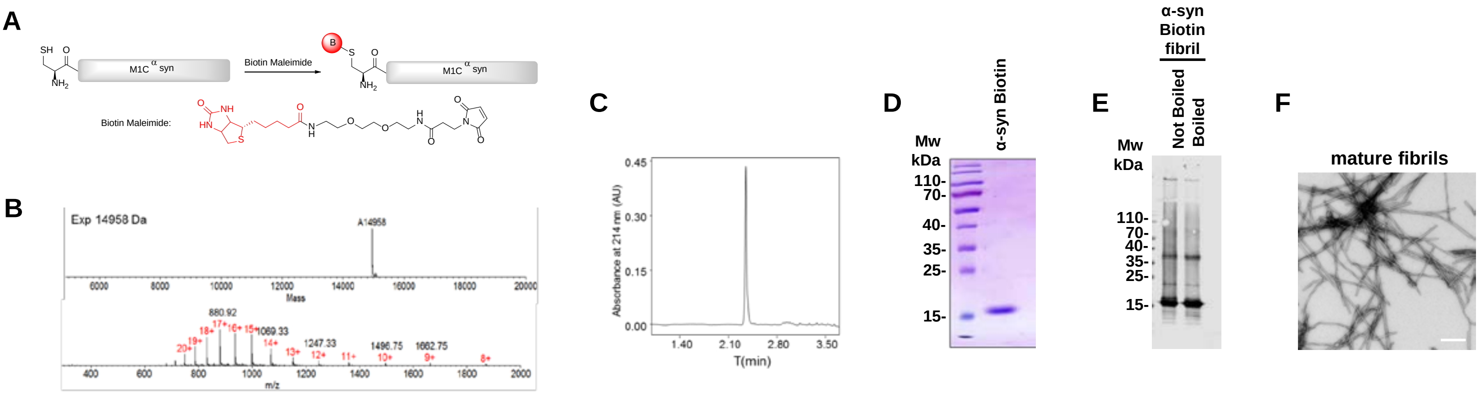


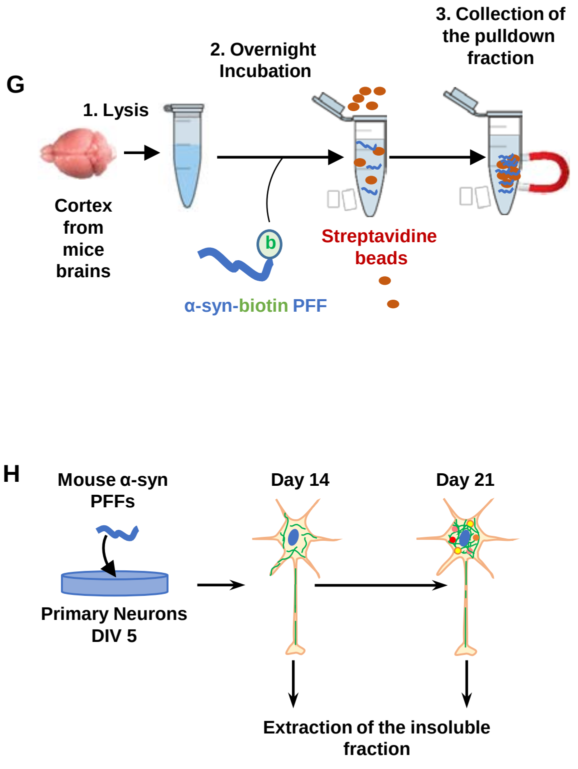


**I**

| C-terminal interactome Proteins name | Insoluble Fraction D14 | Insoluble Fraction D21 | Biotynalated PFF pull down | References |
|---|---|---|---|---|
| Aldoa | + | - | - | McFarland et al., 2008 |
| Ap1b1 | + | - | - | McFarland et al., 2008 |
| Atp5h | + | - | - | McFarland et al., 2008 |
| Gnai1 | + | + | - | McFarland et al., 2008 |
| Gnb1 | + | - | - | McFarland et al., 2008 |
| Gnb2 | + | + | - | McFarland et al., 2008 |
| Hk1 | + | + | + | McFarland et al., 2008 |
| Hspa5 | + | + | - | McFarland et al., 2008 |
| MAP1-B | + | - | + | Jensen et al., 2000 |
| Map1lc3a | + | - | + | McFarland et al., 2008 |
| Myh10 | + | - | - | McFarland et al., 2008 |
| Ndufb11 | + | - | + | McFarland et al., 2008 |
| Rap1a | + | - | + | McFarland et al., 2008 |
| Sdhb | + | - | - | McFarland et al., 2008 |
| Tubb3 | + | + | - | McFarland et al., 2008 |
| Tubb5 | + | - | - | McFarland et al., 2008 |
| Tubb6 | + | - | + | McFarland et al., 2008 |
| Vapa | + | - | + | McFarland et al., 2008 |
| Amph | - | + | + | McFarland et al., 2008 |
| Atp5f1 | - | - | + | McFarland et al., 2008 |
| Csnk1a1 | - | - | + | McFarland et al., 2008 |
| Csnk1d | - | - | + | McFarland et al., 2008 |
| Gabra4 | - | - | + | McFarland et al., 2008 |
| Glul | - | + | - | McFarland et al., 2008 |
| Gnao1 | - | - | + | McFarland et al., 2008 |
| Hsc70 | - | - | - | Gao et al., 2015 |
| Icam5 | - | - | + | McFarland et al., 2008 |
| Immt | - | - | + | McFarland et al., 2008 |
| Immt | - | - | + | McFarland et al., 2008 |
| Kcnab1 | - | - | + | McFarland et al., 2008 |
| Kcnab3 | - | - | + | McFarland et al., 2008 |
| Map1a | - | - | + | McFarland et al., 2008 |
| Map9 | - | - | + | McFarland et al., 2008 |
| Mark1 | - | - | + | McFarland et al., 2008 |
| Mtch2 | - | - | + | McFarland et al., 2008 |
| Ndufa10 | - | - | + | McFarland et al., 2008 |
| Ndufa12 | - | - | + | McFarland et al., 2008 |
| Ndufa13 | - | - | + | McFarland et al., 2008 |
| Ndufa5 | - | - | + | McFarland et al., 2008 |
| Ndufa6 | - | - | + | McFarland et al., 2008 |
| Ndufa8 | - | - | + | McFarland et al., 2008 |
| Ndufa9 | - | - | + | McFarland et al., 2008 |
| Ndufb10 | - | - | + | McFarland et al., 2008 |
| Ndufb4 | - | - | + | McFarland et al., 2008 |
| Ndufs1 | - | - | + | McFarland et al., 2008 |
| Ndufs2 | - | - | + | McFarland et al., 2008 |
| Ndufs3 | - | - | + | McFarland et al., 2008 |
| Ndufs4 | - | - | + | McFarland et al., 2008 |
| Ndufs5 | - | - | + | McFarland et al., 2008 |
| Ndufs7 | - | - | + | McFarland et al., 2008 |
| Ndufs8 | - | - | + | McFarland et al., 2008 |
| Ndufv1 | - | - | + | McFarland et al., 2008 |
| Ndufv2 | - | - | + | McFarland et al., 2008 |
| Phb | - | - | + | McFarland et al., 2008 |
| Poldip3 | - | - | + | McFarland et al., 2008 |
| Ptprd | - | - | + | McFarland et al., 2008 |
| Rab5a | - | - | + | McFarland et al., 2008 |
| Rps2 | - | - | + | McFarland et al., 2008 |
| Samm50 | - | - | + | McFarland et al., 2008 |
| Serf1 | - | - | - | Yoshimura et al., 2017 |
| Slc1a2 | - | - | + | McFarland et al., 2008 |
| Slc1a3 | - | - | + | McFarland et al., 2008 |
| Slc25a1 | - | - | + | McFarland et al., 2008 |
| Slc25a11 | - | - | + | McFarland et al., 2008 |
| Snap91 | - | - | + | McFarland et al., 2008 |
| Stx1a | - | - | + | McFarland et al., 2008 |
| Stxbp1 | - | - | + | McFarland et al., 2008 |
| Syt2 | - | - | + | McFarland et al., 2008 |
| Tfam | - | - | + | McFarland et al., 2008 |
| Trim3 | - | - | + | McFarland et al., 2008 |
| Uqcr10 | - | - | + | McFarland et al., 2008 |
| Uqcrc2 | - | - | + | McFarland et al., 2008 |
| Uqcrfs1 | - | - | + | McFarland et al., 2008 |
| Vamp2 | - | - | - | Buré et al., 2015 |
| Vapb | - | - | + | McFarland et al., 2008 |
| Vdac1 | - | + | - | McFarland et al., 2008 |

Figure S11. Related to Figure 6

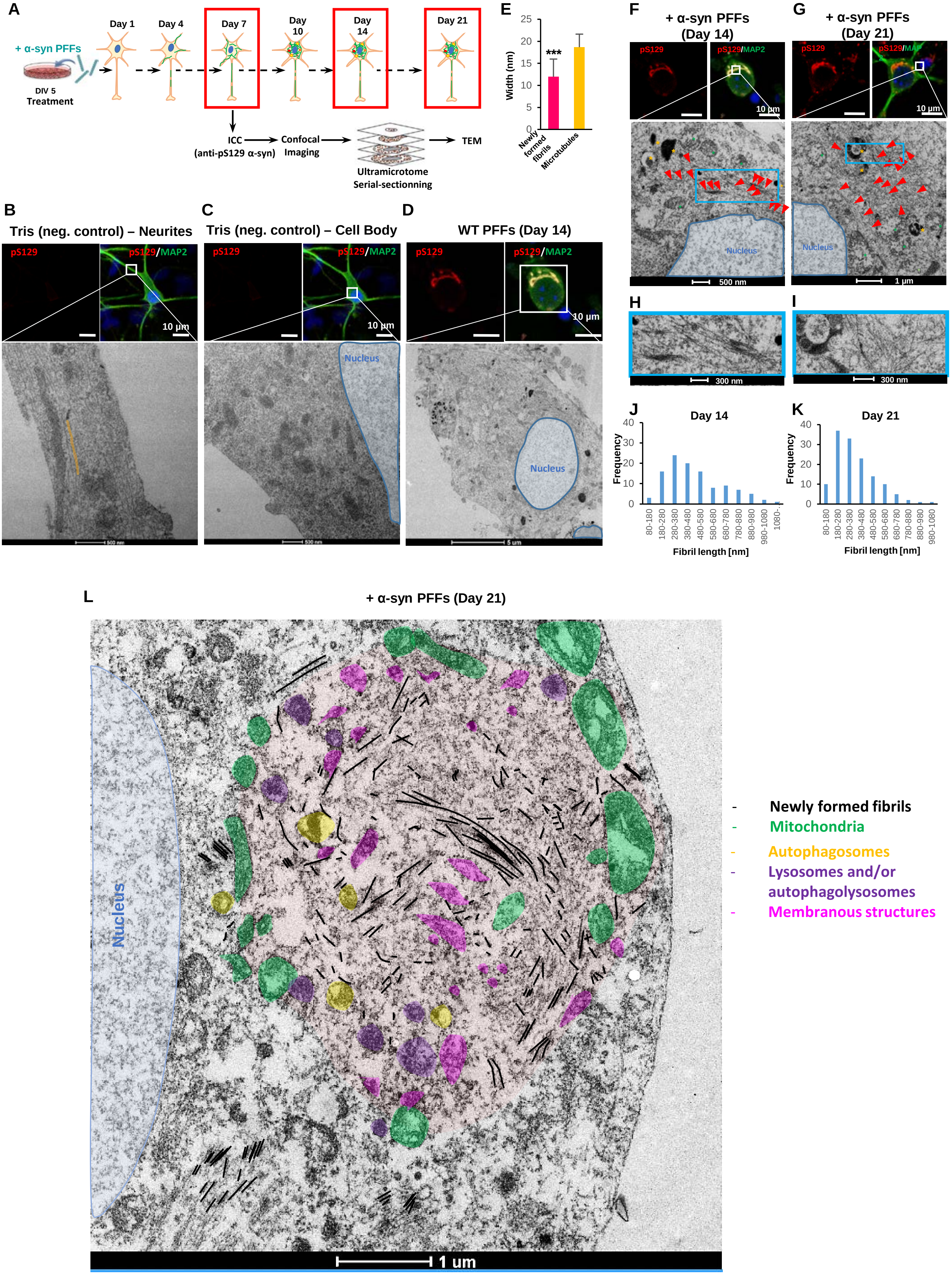

**Figure S12. Related to Figure 7**

**A**

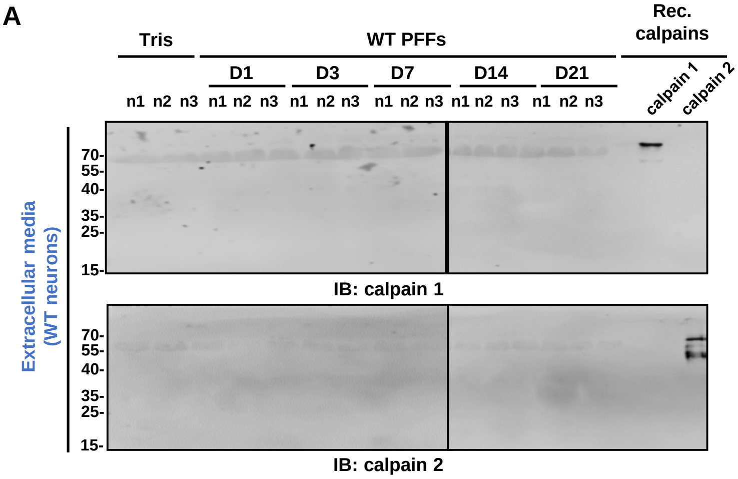


**B**

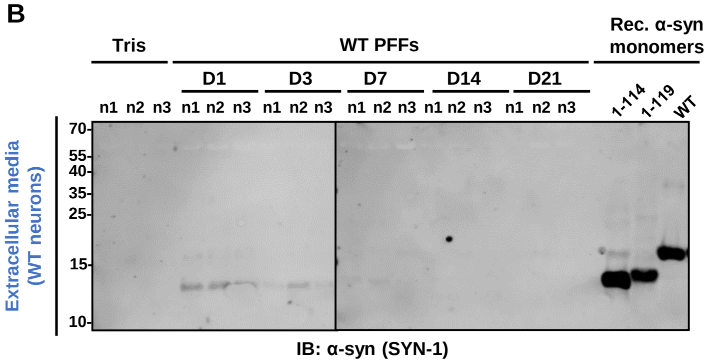


**C**

| Calpain 2 | Fold difference (Tris vs PFFs) | $-\text{Log}_{10}$ (p-Value) |
|---|---|---|
| **Day 7** | 0.04 | 0.11 |
| **Day 14** | 4.05 | 1.67 |
| **Day 21** | 0.98 | 2.77 |

**D**

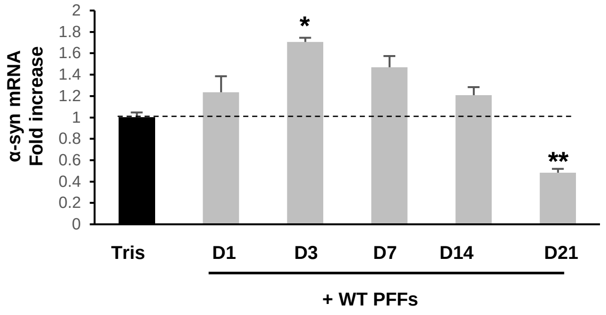


**E**

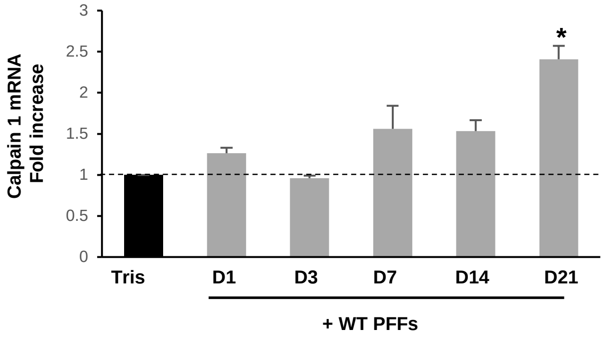


**F**

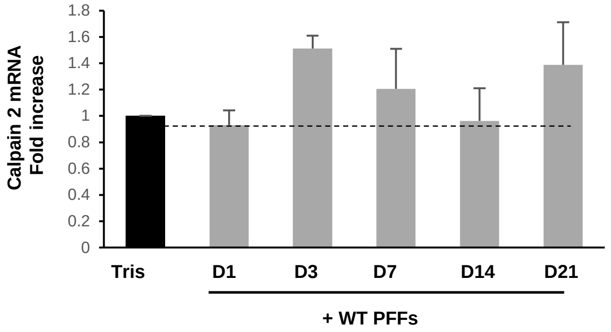


**G**

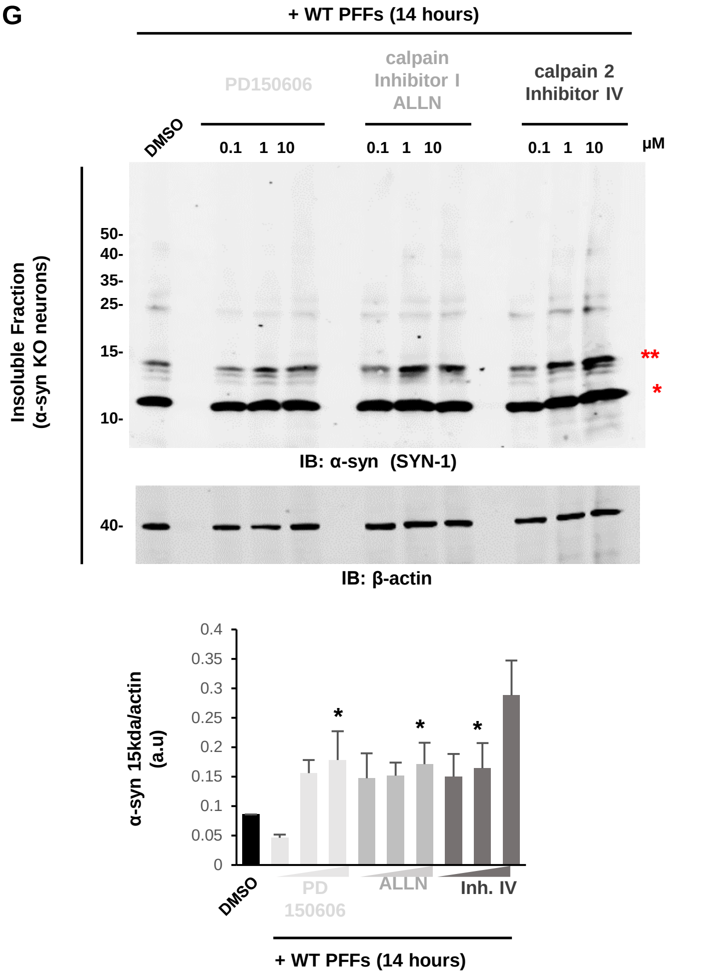

Figure S13. Related to Figure 8

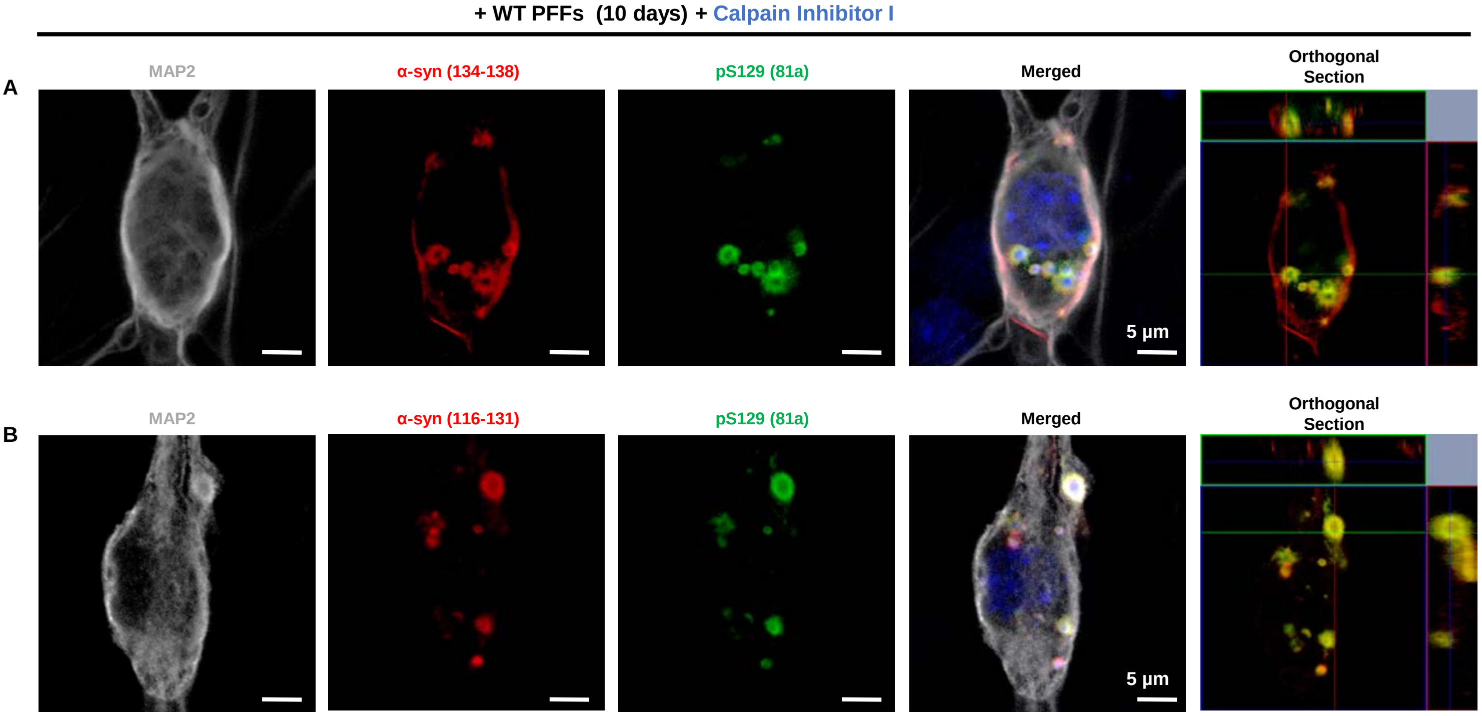

Figure S14. Related to the Discussion

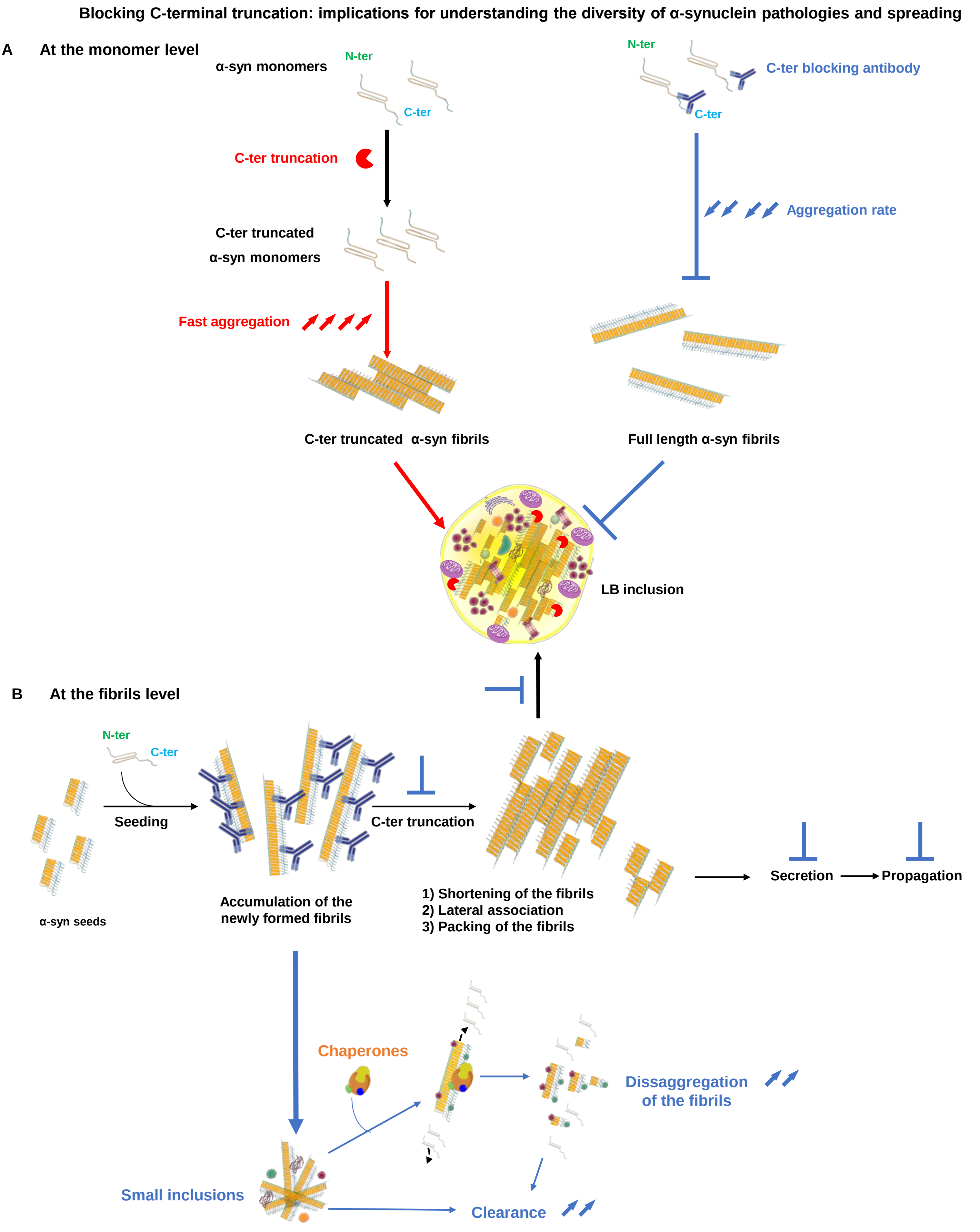
