## Supplementary material for "The making of a Lewy body: the role of α-synuclein post-fibrillization modifications in regulating the formation and the maturation of pathological inclusions": Graphical_Abstract

bioRxiv
THE PREPRINT SERVER FOR BIOLOGY

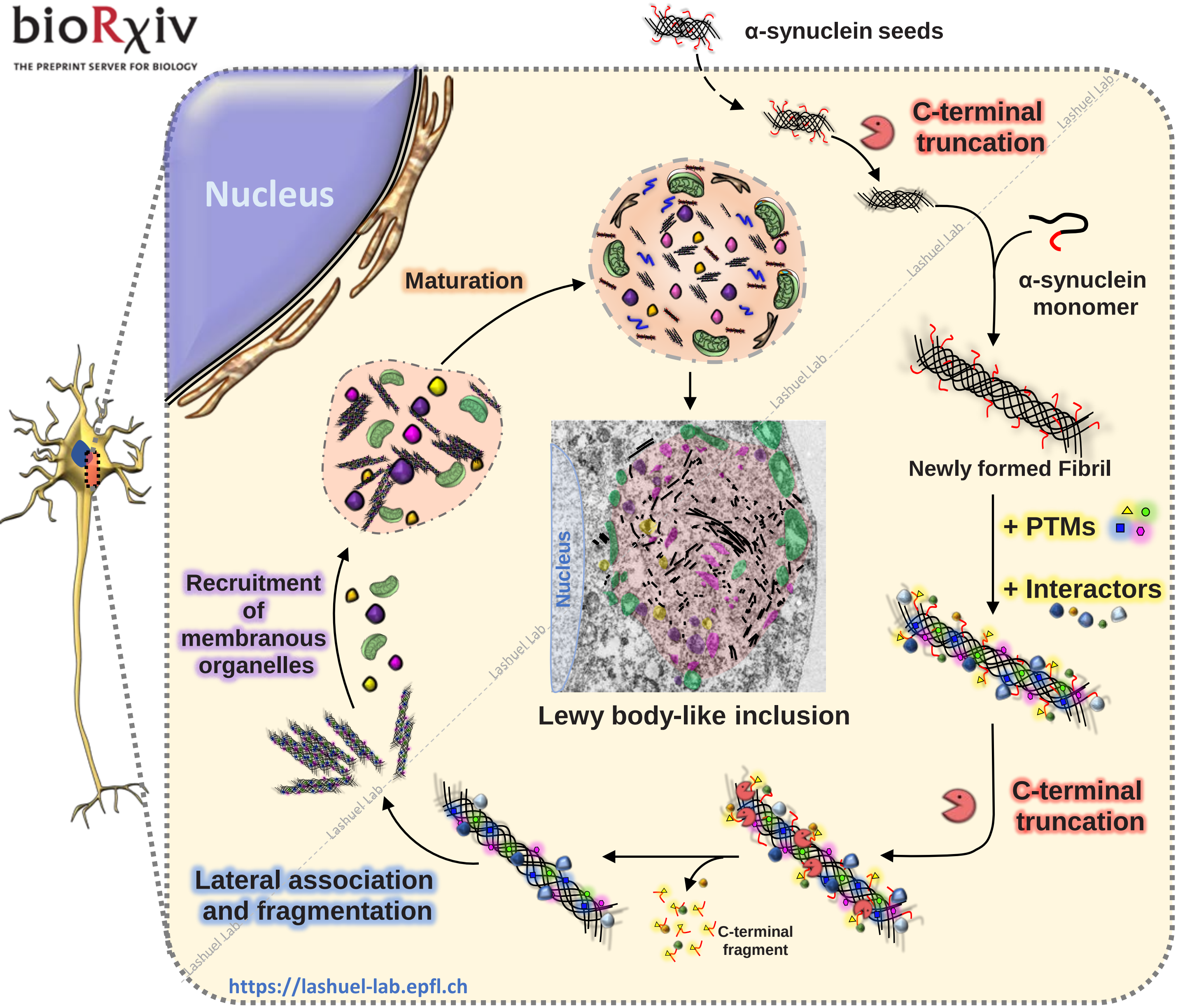


### Highlights

- A neuronal seeding model that recapitulate the process of Lewy body formation.
- Novel insight into the molecular, structural and cellular mechanisms underlying Lewy body formation and maturation.
- Post-fibrillization C-terminal cleavages regulate α-syn inclusions formation by modulating the interactome, processing and lateral association of α-syn newly formed fibrils.
- More tools are needed to enable accurate profiling and assessment of α-syn pathological diversity and spreading in human brain tissues.
